# Alanine-scanning of the yeast killer toxin K2 reveals key residues for activity, gain-of-function variants, and supports prediction of precursor processing and 3D structure

**DOI:** 10.1101/2024.11.22.624868

**Authors:** Rianne C. Prins, Tycho Marinus, Eyal Dafni, Iftach Yacoby, Sonja Billerbeck

## Abstract

Yeast killer toxins (YKTs) are antimicrobial proteins secreted by yeast with potential applications ranging from food preservation to therapeutic agents in human health. However, the practical use of many YTKs is limited by specific pH requirements, low temperature stability, low production yields, and narrow target specificity. While protein engineering could potentially overcome these challenges, progress is hindered by a lack of detailed knowledge about sequence-function relationships and structural data for these often multi-step processed proteins. In this study, we focused on the YKT K2, encoded by the M2 dsRNA satellite virus in *Saccharomyces cerevisiae*. Using alanine scanning mutagenesis of the full open reading frame and structure predictions combined with molecular dynamics simulations, we generated a comprehensive sequence-function map, refined the model for the proteolytic processing of the K2 precursor, and predicted the mature toxin structure. Our findings also demonstrate that K2 can be engineered towards enhanced toxicity and altered target specificity through single-site mutations. Furthermore, we identified structural homology between K2 and the SMK toxin from the yeast *Millerozyma farinosa*. Our cost-effective workflow provides a platform to broadly map YKT sequence-structure-function relationships, facilitating the engineering towards toxin-based technologies. The workflow could also serve as a template to resolve the processing and conformations of other proteins within the secretory pathway – a dynamic multi-step process that is challenging to structurally capture by purification and solving structures of intermediates.

## Introduction

Many yeast secrete antifungal proteins that are lethal to sensitive strains, providing a competitive advantage in ecological niches and shaping microbial communities [1–3]. The genes encoding these yeast killer toxins have been identified on several genetic elements, including on host chromosomes [4,5], on dsDNA virus-like particles [6,7], and on cytoplasmically inherited dsRNA mycoviruses [8]. Some killer toxins target the cell wall, others the plasma membrane or intracellular compartments, to ultimately interfere with vital cellular processes [9]. With a growing threat of antifungal resistance and a need for novel antifungal agents [10], these functionally diverse killer toxins hold potential for various applications, from biocontrol agents for crops and food to therapeutic agents for human health [2,11].

However, there are intrinsic limitations that need to be overcome. The natural production hosts often thrive in ambient temperatures and acidic environments, rendering many killer toxins dependent on low pH and showing low thermal stability, with often low production yields [11]. Since these toxins are ribosomally synthesized proteins, their encoding genes are potentially amenable to engineering towards overcoming these limitations [12]. In addition, it would be of interest to engineer target ranges of these toxins toward narrow- or broad-spectrum applications [11].

K2 is a killer toxin encoded on a dsRNA satellite mycovirus (M2) in *Saccharomyces cerevisiae* [13] (**Figure 1A**). The K2 toxin is produced by commercial wine yeast strains, which may indicate that it can be generally considered safe for consumption [11,14]. The toxin is most active at a temperature of 20-25°C and under acidic conditions - it loses activity completely above pH 5.5 [15]. It is active against several yeasts, including sensitive *S. cerevisiae* strains as well as clinically relevant isolates of *Nakaseomyces glabratus* (previously named *Candida glabrata*), which can have high levels of resistance to current clinical antifungal therapeutics [16]. Therefore, the K2 toxin serves as an ideal model for exploring toxin bioengineering opportunities.

**Figure 1.**
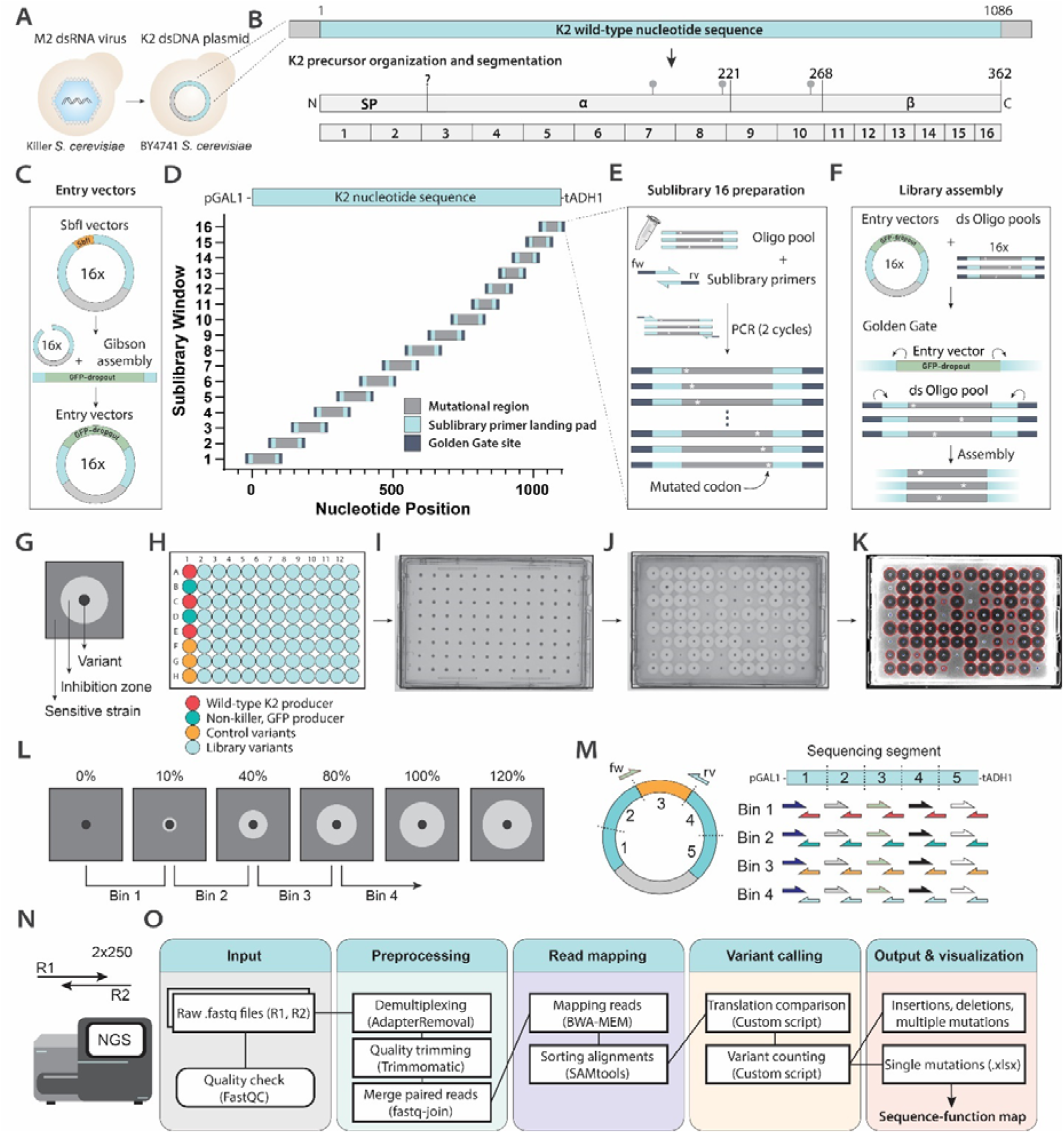
Overview of alanine scanning library creation, variant screening, and next-generation sequencing. (**A**) The K2 killer phenotype is naturally conveyed by the presence of the M2 dsRNA virus in *S. cerevisiae*. Within this study, the sequence encoding K2 was expressed from a pRS423-type dsDNA plasmid. (**B**) The product of the ORF is the K2 precursor, which consists of 362 residues, reportedly organized in a signal peptide (SP) and the α- and β-domains that form the mature toxin. The exact signal peptidase cleavage site is unknown. Two potential canonical dibasic (KR) pKex2 protease cleavage sites are present at residues 220/221 and 267/268. Three potential *N*-glycosylation sites are present at N177, N214 and N261. The ORF was divided into 16 segments to facilitate the cloning of sublibraries using pooled mutant oligonucleotides (**Supplementary Table S1 and S9-S11**). (**C**) Each of the 16 segments was previously replaced by an SbfI restriction site [17], which was used here to insert a GFP-dropout module. (**D**) Each segment sequence was encoded on oligonucleotides in which alanine substitutions were introduced. The mutant oligonucleotides were ordered as 16 sublibrary pools. (**E**) The single-stranded oligonucleotides were double-stranded and type-II restriction sites were added. (**F**) Within a Golden Gate reaction, the GFP-dropout modules were replaced by the respective mutant oligonucleotide pools, introducing the alanine conversions into the wild-type K2 sequence. (**G**) The resulting variants were assessed for formation of a zone of inhibition on a lawn of a sensitive indicator strain, indicating secretion of active toxin. The size of the halo is related to the activity of the toxin. (**H**) *S. cerevisiae* transformants were arrayed into 96-well microtiter plates. In each plate, the first column was dedicated to controls: Three replicates of wild-type K2 producers, two replicates of a non-killer that produces GFP (convenient for determining plate orientation), and three previously constructed control variants with varying expected halo sizes. (**I**) The cultures were first pinned in duplicate onto solid media, under conditions inducing protein expression, to create the source plates. (**J**) From the solid source plates, the colonies were replicated onto an assay plate containing an embedded sensitive strain and halo sizes (zones of inhibition) were imaged after 2 days. (**K**) The areas of the zones of inhibition were quantified using image-processing software [27], adjusted for this purpose. The library was screened in duplicate and the displayed images are representative of the results. (**L**) The different halo area sizes were compared to the wild-type zone of inhibition of the K2 toxin (set to 100%). The variants were divided into four bins based on the area of the halo: Bin 1 (0-10%), bin 2 (10-40%), bin 3 (40-80%) and bin 4 (>80%). (**M**) The K2 sequence was divided into 5 segments for next-generation amplicon sequencing. Amplicons were generated using primer combinations specific for each segment/bin combination that add barcodes and partial Illumina adapters (**Supplementary Table S5**). (**N**) Samples were sequenced using a NGS Illumina platform, yielding 250 bp paired-end reads that cover the entire mutated region. (**O**) Reads were assessed for quality, demultiplexed based on the bin/segment specific barcodes, mapped to the template plasmid, and counts of single, intended, amino acid conversions within each sample were determined using a custom script.

The K2 toxin precursor consists of 362 amino acids (**Figure 1B**). The N-terminal signal peptide directs the downstream amino acid sequence into the secretory pathway and simultaneously functions as the immunity factor that prevents self-killing [17]. Within the early secretory pathway, disulfide bridge formation and glycosylation take place [13]. Within the late Golgi, proteolytic processing occurs by the endopeptidases Kex1p and Kex2p which finally yields the α- and β-subunits that form the mature secreted heterodimeric K2 toxin [13]. Toxin molecules subsequently diffuse into the environment, dock onto 1,6-β-glucan within the yeast cell wall of target cells [18], likely interact with the plasma membrane receptor Kre1p [19] and then act as ionophores to induce membrane permeability [8,20] – a process in which the rather hydrophobic α-subunit seems to play a critical role [17].

The exact molecular mechanism by which the K2 toxin kills cells is however not completely understood. The lack of efficient protocols to purify the secreted K2 toxin from the culture media has hindered structural studies. Previous studies have investigated the importance of only a limited number of residues by site-directed mutagenesis, such as two putative protease processing sites [13] (**Figure 1B**). In addition, the high primary sequence diversity of killer toxins and lack of identified homologous (toxin) sequences have limited the ability to perform sequence comparisons to identify critical conserved residues. Without precise data on precursor processing sites, the prediction of the mature toxin subunits and the associated three-dimensional protein structure is challenging. Gaining insights into sequence-function relationships and the mature toxin structure is essential for advancing our molecular understanding of yeast killer toxins and facilitating rational design in engineering strategies.

To gain insights into the sequence-function relationships of the K2 toxin, we performed a systematic alanine scanning mutagenesis of the full K2 open reading frame (ORF). Here, each substitution into alanine examines the contribution of individual side chains to the fitness of the toxin. To build the protein variant libraries, we used a plasmid-based system (**Figure 1A**), leveraged oligo pools as an affordable way of DNA synthesis to encode all the library variants [21] and assembled the library using Golden Gate cloning [22]. Since K2 is a secreted protein, variants were spatially separated into an arrayed format and screened based on the formation of zones of growth inhibition on solid media plates seeded with a sensitive strain. Recent advancements in protein structure prediction further contributed to the construction of precursor and mature toxin structure predictions [23–25], which were further refined by molecular dynamics simulations.

This workflow yielded a sequence-function map that identified critical residues for toxin activity and also yielded variants with enhanced function and altered species-specificity. Together with map-inspired additional experiments and structure predictions, this enabled us to create a new model for K2 precursor processing, and a prediction for the mature K2 toxin structure. We further identified that the K2 toxin shares structure similarities with the SMK killer toxin, for which a crystal structure is available.

## Results

### 1. Creation of the alanine scanning mutagenesis library, variant screening and sequence-function linkage

To facilitate genetic manipulation, the K2 sequence from the M2 dsRNA virus was expressed from a pRS423-type high-copy plasmid backbone using the galactose-inducible promoter (pGAL1) from the widely used yeast toolkit [26] (**Figure 1A**). We had previously divided the K2 coding sequence into 16 segments and replaced the segments by an SbfI restriction site [17] (**Figure 1B**) (**Supplementary Table S1**) and here, to create entry vectors, replaced each segment by a Golden Gate-compatible GFP-dropout cassette which can be expressed in *Escherichia coli* (**Figure 1C**). The alanine scan involved all codons of the K2 ORF besides the stop codon (362 codons). Native alanine residues were converted into glycine. The complete alanine scanning library was encoded on and ordered as 16 oligonucleotide (oligo) pools (one for each segment) and cloned into the 16 entry vectors using Golden Gate as outlined in **Figure 1C-F** (**Supplementary Table S9-S11**) [17,22].

After transforming each segment-specific sublibrary into *E. coli*, green-white screening was used to select white colonies that carried constructs with a successful exchange of the GFP-dropout module for a mutant oligo. Based on the number of white colonies, all sublibraries were estimated to be complete (assuming a uniform distribution of mutants) (**Supplementary Table S2**). A high completeness was confirmed by next-generation sequencing (NGS) for all sublibraries (99% complete) except for the sublibrary of segment 5, which was subsequently successfully recloned. Only few mutations with no or low counts at this stage were I228A, Y276A, G280A and Q311A, and, consistently, these variants were also missing later in the screened library in yeast (**Supplementary Figure S3**).

The sublibrary plasmids were extracted from the pooled white *E. coli* colonies and used to transform *S. cerevisiae* BY4741, which lacks the original M2 dsRNA satellite virus. Since the K2 toxin is a secreted protein, spatial separation of the variants was required for library screening. The transformants were therefore arrayed into 96-well plates with internal controls in each plate, including three wild-type K2 producers, and subjected to a halo assay to assess toxin activity (**Figure 1G-J**). This coupled toxin activity to a screenable phenotype, since the size of the formed zone of inhibition is related to toxin functionality. The assay provides the most stringent test for function: The toxin needs to be efficiently transcribed, translated, processed, folded, secreted, and needs to be stable and to diffuse in the agar media and interact properly with target cells. Any aspect that was impacted by an alanine conversion should be evident in the resulting zone of inhibition. In total, 1228 yeast colonies were screened, providing a theoretical probability of 0.96 for library completeness (assuming a uniform distribution of mutants) (**Supplementary Table S3**).

The variants created a variety of zone of inhibition (halo) sizes (**Figure 1J**). We quantified the size of each halo using a modified version of the image-processing software CFQuant (**Figure 1K**) [27]. To account for slight plate-to-plate variations in halo sizes, we subsequently normalized the halo areas within each plate to the internal wild-type K2 producer controls, before cross-plate comparison. We determined a confidence interval for wild-type toxin activity between 80-120% of the average wild-type halo size (see **Methods**) and then divided the variant colonies into 4 activity bins from loss-of-function to wild-type activity (**Figure 1L**).

We observed that 13.2% of all the screened colonies showed loss-of-function (bin 1, 0-10% of wild-type halo size), 9.7% of colonies showed a major negative effect (bin 2, 10-40%), 26.1% showed a moderate negative effect (bin 3, 40-80%), and 49.8% of the colonies retained wild-type level activity (bin 4, >80%) (**Supplementary Figure S1, Table S4**). Interestingly, some colonies (1.3%) created increased halo sizes (>120%), in particular with mutations towards the C-terminal region.

To create genotype-phenotype links on residue-level, we combined the colonies based on the four activity bins and prepared samples for NGS analysis (**Figure 1L-N**). Since the K2 ORF (362 codons) was longer than our NGS read length, the ORF was divided into 5 sequencing segments which were eventually amplified from the samples by barcoded primers, each sufficiently short to be covered by 250 bp paired-end NGS reads (**Figure 1M and N**) (**Supplementary Table S5**).

The sequencing yielded a total of 305.651 paired-end reads. The reads were assigned to the respective bin/segment combination by demultiplexing based on the dual barcode sequences. After quality filtering, paired-end reads merging and mapping, 243.321 reads (79.6%) were yielded for input into a custom script for computational analysis (**Figure 1O**). The custom script performed further stringent filtering to yield only high-quality sequencing reads and subsequently performed variant calling and counting of occurrences of single-amino acid mutations. This yielded a total of 224.328 read sequences that passed filtering (92.2% of input reads), of which 180.222 reads contained single, intended mutations (80.3% of used reads). The resulting coverage per variant is displayed in **Supplementary Figure S2A**.

The distribution of read counts of single, intended mutations followed a Gaussian distribution (**Supplementary Figure S2B**). The distribution had a relatively wide standard deviation, indicating that variants were represented in different quantities. This could be caused by a varying oligo distribution within the synthesized oligo pools and/or the stochastic process of colony selection, resulting in varying variant representation numbers on the assay plates, and/or differences in amplicon ratios arising during the sample preparation process. However, only 11 variants did not yield any reads, and only 11 variants were significantly overrepresented. 50% of positions had variant counts between 243 and 684 reads (**Supplementary Figure S2B**).

The K2 variants were subsequently linked to a functional score. Loss-of-function (i.e. variant was found in bin 1) was assigned a score of 1, whereas wild-type activity (i.e. variant was found in bin 4) was assigned a score of 4. We found that the reads of some variants were divided over two adjacent bins, in which case we assumed that the variant had a functionality on the border of the bins and we therefore averaged the functional score (resulting in scores of 1.5, 2.5 or 3.5, respectively for 3, 16, and 40 variants). In this way, we were able to determine functional scores for 340 out of the 362 positions (93.9%), rendering the library near-complete (**Supplementary Table S6 and S7**). A score of 0 indicated that the variant data did not pass the quality thresholds: 22 positions (6.1%) were excluded from analysis, either because of no or too few read counts (12 cases), or read counts were significantly spread over two non-adjacent bins (10 cases). In case of the latter, we speculate that one clonal population may have contained two plasmids, was a mixed colony, or contamination of cells or barcoded primers occurred during the handling process.

From the resulting data we created an alanine scanning-based sequence-function map of the K2 killer toxin. We created maps to determine the global distribution of functional scores along the K2 ORF (**Figure 2A and B**) as well as a detailed per-residue function map (**Supplementary Figure S3**).

**Figure 2.**
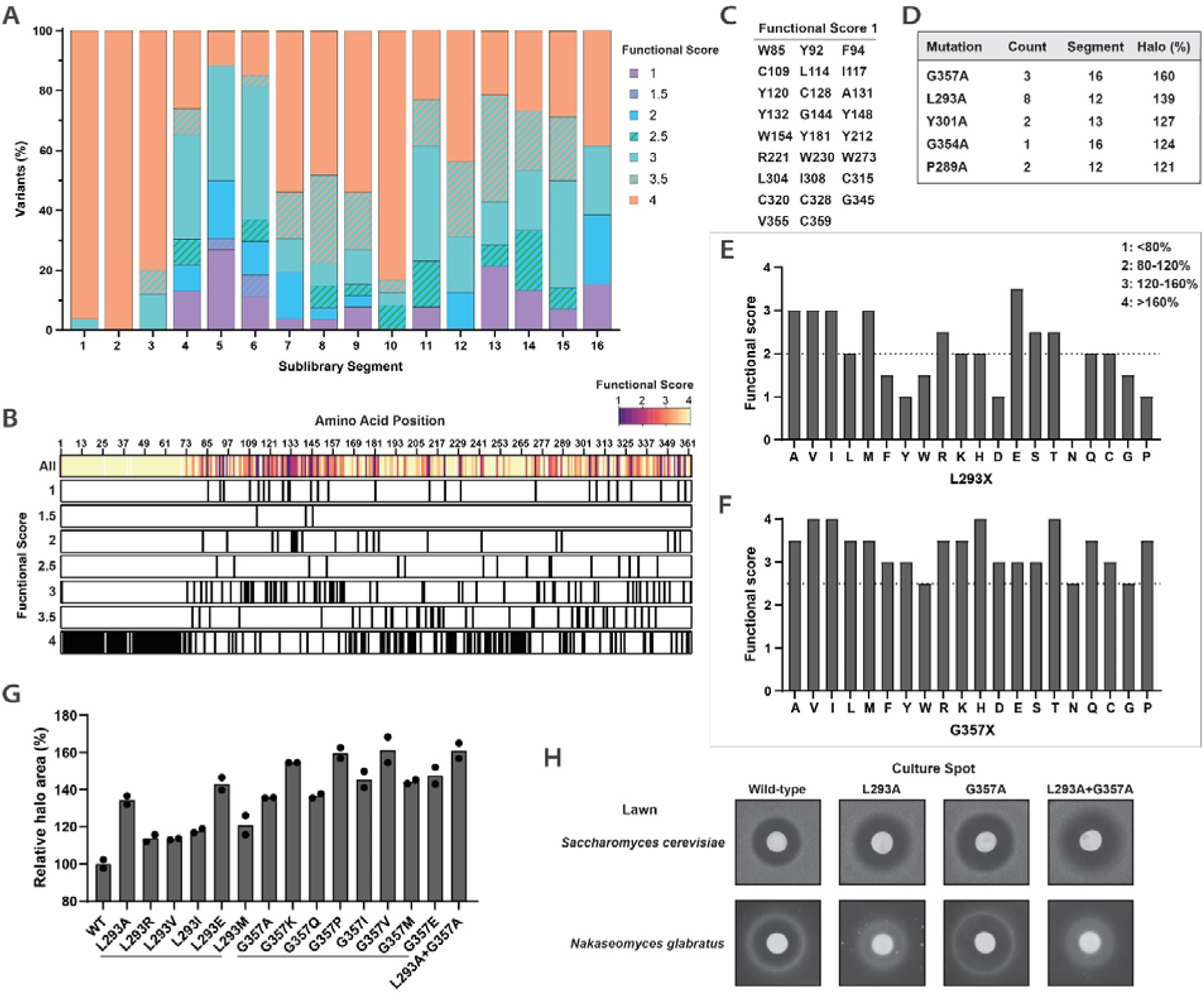
Variant functional scores distribution along the K2 ORF and gain-of-function variants. (**A**) Distribution of the functional scores per sublibrary segment of the K2 ORF (**Supplementary Table S1**). Per segment, the percentage of all positions within that segment with a given functional score is indicated. (**B**) Functional scores displayed along the K2 ORF for individual positions. (**C**) Residues with functional score 1 (loss of function). (**D**) 16 colonies with increased halo sizes were sequenced, and several mutations associated with gain of function were identified. For the relative halo area size, the wild-type halo was set to 100%. (**E**) The determined functional scores of the saturation mutagenesis library of position 293 are displayed. The horizontal line indicates the wild-type. Bin borders specific for figure panels E and F are displayed in the right top corner. (**F**) The determined functional scores of the saturation mutagenesis library of position 357 are displayed. The horizontal line indicates the wild-type. (**G**) A number of plasmids was extracted from library variants, transformed in *E. coli*, purified from *E. coli* and reintroduced into *S. cerevisiae.* The resulting transformants were screened in duplicates for production of a zone of inhibition, of which the area was quantified after 2 days of growth displayed relative to the average of the wild-type, which was set to 100%) ((**Supplementary Figure S5**). WT; plasmid encoding wild-type K2. Dots indicate the individual replicate values, while the bar indicates the mean. (**H**) The OD_600_ of *S. cerevisiae* overnight cultures - expressing either wild-type K2, K2 with an L293A or G357A mutation or L293A+G357A - was normalized and 5 µL was spotted on top of either a *S. cerevisiae* Δ*pbs2* or *N. glabratus* indicator lawn. Zones of inhibition were imaged after 2 days of growth.

### 2. Alanine scanning results in a range of phenotypes from loss of function to gain of function

We found that 50.0% of the analyzed amino acid positions (170/340) resulted in wild-type toxin activity (functional score 4) after conversion into alanine (or alanine to glycine) (**Supplementary Figure S3, Table S6 and Table S7**). These respective amino acid sidechains were therefore not individually critical for K2 toxin function. This corresponds with the fact that alanine substitutions are considered relatively mildly disruptive compared to some other amino acids, and will mainly identify key residues [28].

Many positions within the N-terminal region were highly tolerant to alanine conversions (**Figure 2A and B**): Sublibrary segments 1 and 2 (corresponding to residues 1-54) contained 96% and 100% of their analyzed variants, respectively, in the wild-type activity bin (functional score 4) (**Supplementary Table S7**). This is consistent with previous observations showing that the N-terminus acts as a signal peptide and is important for self-immunity, but that the individual residues are not essential for K2 toxin activity [17]. Since signal peptides of *S. cerevisiae* often contain alanine, and are dependent on overall features rather than specific amino acid positions [29], it was not surprising that active toxin secretion was not affected by alanine conversions within this region. The M1A mutation resulted in a decrease in halo size, further consistent with previous observations [17].

On the other hand, 7.7% (26/340) of the analyzed positions lost toxicity after conversion into alanine, indicating that these sidechains are highly important for toxin function (functional score 1) (**Figure 2B and 2C**). From the 9 cysteines in the K2 precursor, 6 were found to be essential for toxin activity (**Supplementary Table S6**). Further, many of these residues were hydrophobic and aromatic; 4 of the 5 tryptophan (W) residues lost function when mutated into alanine (W156 had functional score 3). Of the 26 tyrosine (Y) residues, 6 could not be converted into alanine without loss of function. Two glycine residues could not be converted into alanine (G144, G345), and one alanine residue (A131) could not be converted into glycine without loss of function.

When plotting the distribution of functional scores along the K2 ORF, the overall importance of certain regions became visible (**Figure 2A and 2B**). Sublibrary segments 5 and 6 which are located within the α-subunit (corresponding to residues 109 to 162, **Supplementary Table S1**), part of the ultimate toxic subunit [17], contained the least intra-segment percentage of variants with wild-type activity (only 12% and 15% with functional score 4, respectively) and a high percentage of loss-of function variants (27% and 11% with functional score 1, respectively) (**Supplementary Table S7**). In contrast, sublibrary segment 10 was relatively tolerant to alanine conversions compared to other α- and β-domain segments (83% of positions had functional score 4).

Remarkably, sublibrary segments 12, 13 and 16 contained gain-of-function variants with increased halo sizes (**Supplementary Figure S1**).

### 3. Single-residue substitutions yield gain-of-function variants with higher toxicity and altered target specificity

We investigated these gain-of-function variants further. First, we used Sanger sequencing to directly identify the genotype of these colonies. We found several variants with the same mutations, all located within the β-domain (**Figure 2D**). While P289A, Y301A and G354A mutations resulted in relatively small increases in activity (121-127%), the L293A and G357A mutations showed an average halo size of 139% or 160%, respectively, compared to wild-type toxin.

We then looked further into the L293 and G357 positions to identify whether the increase in halo size was a specific result of a conversion into alanine, or whether other amino acid substitutions could lead to the same effect. The modular design of the sublibraries facilitated the swift creation of additional libraries for these two positions. We designed oligo pools encoding saturation mutagenesis and cloned and screened the two resulting libraries in the same way as described above (**Figure 1**). Many of the variants created large zones of inhibition. For preparation of these sequencing samples, we therefore used bin borders here that focus more on the variants with enhanced function: Bin 1 contained all variants <80% of wild-type (loss-of-function), bin 2 those of 80-120% (wild-type), bin 3 those of 120-160% (gain of function) and bin 4 those with halo sizes >160% compared to wild-type (significant gain of function) (**Supplementary Table S8**). NGS yielded 48.046 reads in total. After processing, the lowest number of read counts for a variant was 157, enabling assignment of functional scores with confidence (**Supplementary Figure S4**). One variant (L293N) did not yield any reads, and was likely missing from the screened library. The functional score was again related to the bins (i.e. a functional score of 1 indicated that the variant was detected in bin 1, a functional score of 4 indicated presence in the bin 4 sample, and a functional score of 2.5 indicated that reads were found in both bin 2 and bin 3).

Since the oligonucleotide pools encoded a complete saturation mutagenesis, including the wild-type sequence, these variants provided an additional internal control for wild-type toxin activity. For the assay plate containing the variants of position 293, the wild-type was identified in bin 2, as expected. However, for the plate containing mutations of position 357, we found that the wild-type was present in bin 2 but also partially in bin 3, and we therefore assigned it a functional score of 2.5. We attribute this discrepancy to the fact that this plate contained a high number of large, sometimes overlapping halos, which probably resulted in less accurate quantification of some halo sizes.

Based on these data, we determined that multiple other amino acid substitutions at positions 293 and 357 also resulted in increased halo sizes (**Figure 2E and F**). For position 293, especially L293E, but also L293V, L293I and L293M showed increased halo sizes besides L293A (**Figure 2E**). Aromatic sidechains (F, Y, W) were not well tolerated at position 293 and resulted in loss of function, as well as residues that alter conformational stability of the backbone (G, P) (**Figure 2E**). For position 357, especially G357V, G357I, G357H, and G357T displayed increased halo sizes, but also G357L, G357M, G357R, G357K, G357Q and G357P, besides G357A (**Figure 2F**). Position 357 appeared rather tolerant to amino acid substitutions, with no loss-of-function mutations compared to the wild-type but several substitutions that led to an increase in the halo size. Overall, mutations at position G357 led to larger increases in halo size than those at position L293.

We further validated that the effects were due to the single amino acid substitutions by sequencing a number of individual colonies from the library source plate using colony PCR products, extraction of plasmids from different variants, reintroduction of the plasmids into fresh non-killer yeast, and assessment of the toxic phenotype in a manual halo assay (**Supplementary Figure S5**). We quantified the sizes of the resulting halos (**Figure 2G**). Consistent with the previous results, L293A and G357A again resulted in increased zones of inhibition. For position 293, variant L293E showed the largest increase in halo size, consistent with the NGS data. For position 357, all tested variants again resulted in an increase in the halo size, especially G357K, G357P, and G357V.

In addition, we constructed a variant containing both the L293A and G357A mutations (L293A+G357A). This variant demonstrated an increased zone of inhibition, larger than the two separate mutations, suggesting that the two mutations may affect a different aspect of toxin activity (**Figure 2H**). We further tested the L293A, G357A and the combined L293A+G357A variants against a second species that is targeted by K2 – the human pathogen *N. glabratus*. Here, the G357A mutation also resulted in an increased halo size, but the L293A mutation and the combined L293A+G357A variant demonstrated a reduced activity against *N. glabratus* (**Figure 2H**). In fact, multiple other L293 and G357 mutations that showed an increase in activity against *S. cerevisiae* resulted in reduced toxicity against *N. glabratus* (**Supplementary Figure S5**). Within our libraries we did not find mutations that had the opposite effect.

The different specificity against different target species is an indication that not simply a difference in expression/secretion or diffusion of the toxin through agar is causing the observed effect, but possibly something mechanistic in the interaction with the target cell. Together, these results support that K2 can be engineered towards enhanced killing activity and an altered target spectrum via mutagenesis.

### 4. Residues L29, F31 and F32 potentially play a key role in immunity

On the halo assay plates from sublibrary segments 1 and 2, we further noticed that a few colonies had consistently smaller colony area sizes than average on both replicate plates. We used Sanger sequencing to determine the individual genotypes of these colonies, and found mutations L29A (3 colonies, average colony size 36%, average halo size 87%), F31A (2 colonies, average colony size 42%, average halo size 87%), and F32A (1 colony, colony size 39%, halo size 85%). Because these colonies, despite being small, still produced large zones of inhibition, we speculated that the small-colony phenotype could be a consequence of suicidal behavior. It has been shown previously that mutagenesis of this N-terminal region can result in loss of immunity which can lead to a reduced biomass of the producer spot [17], suggesting that residues L29A, F31A and F32A may play key roles in establishing self-protective immunity.

### 5. The K2 precursor contains a putative proregion with a Kex2p cleavage site after position R79

Next, we aimed to use the alanine scanning data as a basis to address key questions regarding precursor processing and folding of the mature heterodimeric K2 toxin.

The precursor undergoes proteolytic cleavage by a signal peptidase early in the secretory pathway and by the Kex1p and Kex2p proteases in the late Golgi. An early study proposed that the K2 precursor was proteolytically processed after residue R221 by Kex2p, after which Kex1p removes the basic residues K220 and R221, but that residues K267/R268 may be involved in toxin activity rather than processing [13] (**Figure 1B**). Based on the R221 cleavage site, it has been suggested that the α-subunit consists of 172 residues and that the β-subunit consists of 140 residues [30]. However, we here further refine boundaries for the α- and β-domains based on novel data.

We began by examining the N-terminal precursor processing, focusing on determining the starting point of the mature α-domain. As mentioned, the N-terminal region was highly tolerant to the introduced mutations. Between positions 2 and 72, all analyzed variants resulted in wild-type level activity (67 out of 71 positions, with 4 residues not being present in the library), while all analyzed variants between residues 73 to 81 had at least a functional score of 3 (8 out of 9 positions, with 1 residue not being present in the library) (**Supplementary Figure S3, Table S7**). We had previously established that residues 1-54 were sufficient to function as a secretion signal and in addition encode a functional immunity factor [17]. We had proposed that the signal peptide might be cleaved by signal peptidase after position 54 [17], but it was not clear if the α-domain would start directly after that residue.

Besides a signal peptide (preregion), other precursors of dsRNA-encoded yeast killer toxins from *S. cerevisiae* often contain a proregion as well, which is removed later within the secretory pathway. Both the K1 and K28 toxin precursors – which are also encoded on dsRNA mycoviruses in *S. cerevisiae* - contain such a proregion (or δ-subunit). Because we found that many side-chains of residues directly after position 54 were not critical for toxin function, we hypothesized that a similar proregion may be present in the K2 precursor.

Therefore, in addition to the alanine scanning data of individual positions, we first determined whether deletion of the entire respective region affected α-subunit toxicity. The α-subunit is suicidal to producer cells when fused to the yeast α-mating factor (αMF) signal peptide [17]. We created N-terminally truncated versions of the region containing the α-subunit to determine the minimal sequence required for toxicity. Using the suicidal phenotype as a readout, we determined that the region up to position 80 was not critical for α-subunit toxicity (**Figure 3A**). Expression of residues 90-219 resulted in a reduced suicidal phenotype, whereas residues 100-219 did not yield a functional toxic α-subunit (**Figure 3A**). In addition, expression of the different α-subunit regions fused to the native K2 signal peptide (here residues 1-54), which also confers immunity to the K2 toxin [17], did not result in a suicidal phenotype (**Figure 3B**), indicating that the deleted residues were also not involved in immunity.

**Figure 3.**
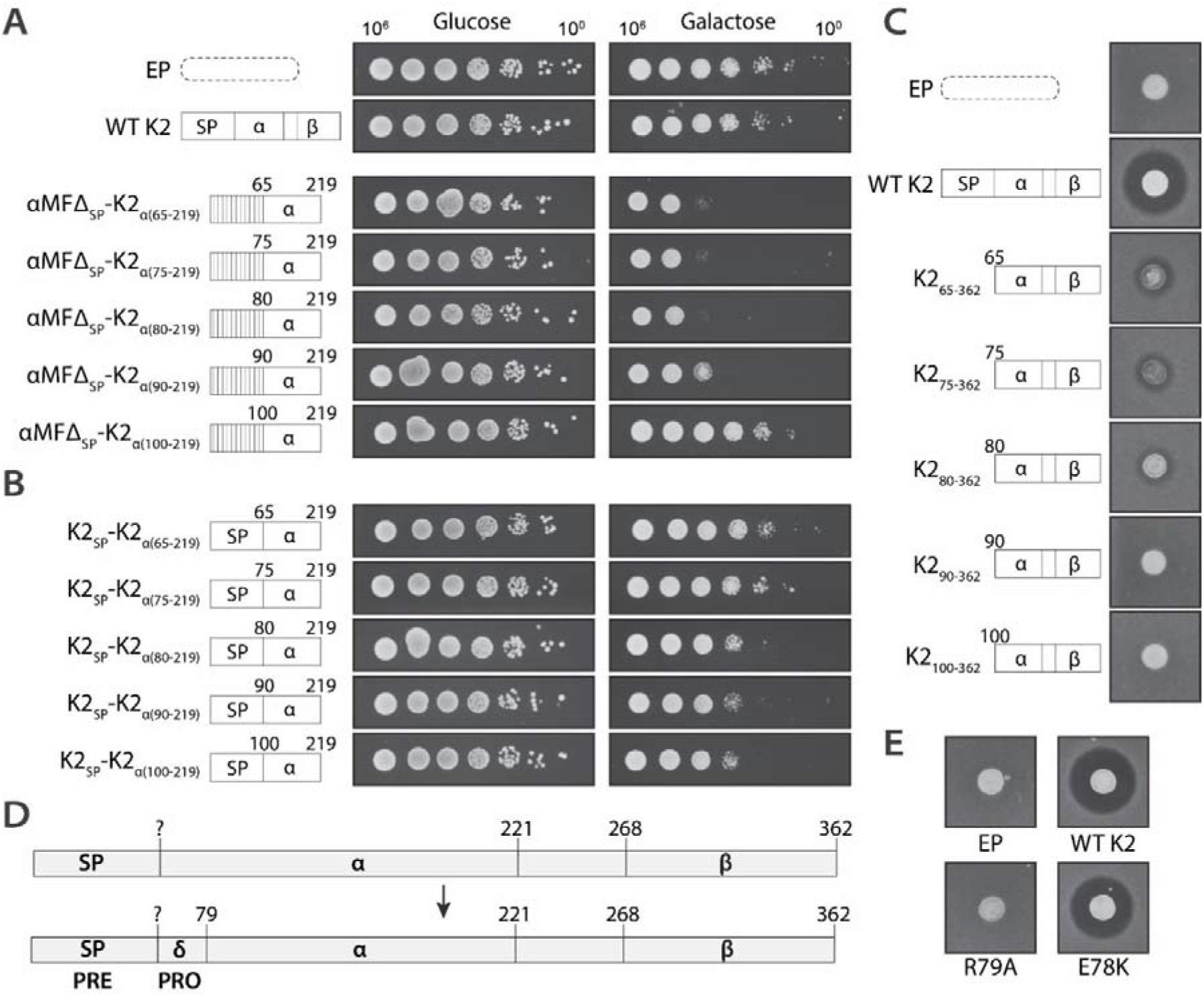
The K2 proregion (δ-domain). The suicidal effect of expression of truncated K2 cytotoxic α-subunit was assessed when the sequence was either fused to (**A**) αMF_SP_; a signal peptide of the α-mating factor, or (**B**) K2_SP_; native K2 signal sequence region consisting of residues 1-54, conferring immunity to wild-type K2 toxin. Cell cultures normalized for optical density and were diluted 10-fold (∼10^6^-10^0^ cells) and spotted onto plates with non-inducing (glucose) or inducing (galactose) conditions. Plates were incubated for 3 days before imaging. EP; empty plasmid. WT K2; wild-type K2. (**C**) Cell cultures with the indicated constructs were spotted on top of agar containing a sensitive indicator strain. Zones of inhibition formed by active toxin secretion were imaged after 2 days. (**D**) Schematic overview of the new proposed K2 precursor domain organization, adding the proregion (δ-domain) between positions ∼54 and 79. (**E**) The remaining toxin activity of variants R79A and E78K was tested in a halo assay. Cell cultures, normalized for optical density, were spotted on top of agar containing a sensitive strain, and the formed zones of inhibition were imaged after 2 days.

Second, we also expressed similar N-terminally truncated versions of the full K2 sequence to determine the impact of the truncated residues on mature toxin secretion. We observed that a small amount of active toxin secretion was still achieved after expression of residues 80-362, while no zone of inhibition was detected after expression of residues 90-362 (**Figure 3C**). The sequence between residues 80 and 90 (“GGFQAWAVGAG”) contains some rather hydrophobic residues, which we speculate may be responsible for the observed remaining – albeit inefficient - toxin secretion.

In conclusion, the region 1-80 seemed involved in toxin secretion, but not required for ultimate toxin activity. Because residues 1-54 had been shown to be sufficient to function as a secretion signal (preregion) [17], and because active toxin (although inefficiently) could be secreted from position 80, we hypothesized that the region in between (residues ∼54-80) may serve as a proregion. If this is the case, then we expect an appropriate cleavage site to be present that would release the proregion from the remainder of the precursor. In the case of two other dsRNA-encoded killer toxins from S. cerevisiae, K28 and K1, Kex2p cleaves the proregion after residues ‘ER’ (K28, position 48-49) [31], or ‘PR’ (K1, position 43-44) [32]. The Kex2p protease is dependent on an arginine at position P1, but while K or R is preferred at position P2 to form a canonical dibasic site, several other amino acids can be present, resulting in varying cleavage efficiencies [33]. In support of our hypothesis, K2 contains an ‘ER’ sequence at position 78-79. Therefore, we propose that a proregion is present within the K2 precursor, cleaved after residue R79 by Kex2p (**Figure 3D**).

For the K1 toxin, it has been determined that an R44A mutation resulted in loss of toxicity due to disruption of the proregion cleavage site [34]. Unfortunately, variant R79A was one of the variants within our library that did not pass the set quality thresholds. We therefore separately cloned this variant, assayed the production of a zone of inhibition, and found that the R79A variant as expected leads to loss of function (**Figure 3E**). An E78K mutation, which would yield a canonical dibasic ‘KR’ site, resulted in a wild-type level active toxin secretion (**Figure 3E**), similar to the E78A mutation in our library (**Supplementary Figure S3**), indicating that this P2 site is flexible towards processing efficiency. A saturation mutagenesis study had shown that compared to a ‘KR’ site, the ‘AR’, ‘ER’ and ‘PR’ sites resulted in approximately 10% cleavage efficiency [33].

Interestingly, if this is the case, K1, K28 and K2 (all are toxins encoded on *S. cerevisiae* dsRNA mycoviruses) use only a moderately processed recognition signal for Kex2p at the proregion cleavage site, indicating there may be a underlying functional relevance selecting for moderate cleavage efficiency [33,35]. In addition, the K74 toxin from *Saccharomyces paradoxus* (encoded on the M74 dsRNA virus) contains a similar genetic organization and also contains a putative noncanonical proregion ‘VR_36_’ Kex2p cleavage site [36]. An R36K mutation resulted in secreted but inactive K74 heterodimers [36]. It has been speculated that the proregion of these toxin precursors may be involved in inactivation of the toxin until it is secreted [34].

Summarizing, our data suggest the presence of a K2 proregion (δ-domain), which is cleaved in the secretary pathway via a low-efficiency Kex2p processing site, and that the α-subunit only starts from residue 80 onwards (**Figure 3D**).

### 6. K2 precursor structure prediction indicates fold similarities with the SMK toxin from the yeast *Millerozyma farinosa*

After refining the model for N-terminal precursor processing, we used the sequence-function data in combination with protein structure predictions to gain insights into further processing of the precursor. Since precursor processing dictates the subunits of the ultimately secreted mature toxin, this knowledge is essential for predicting the structure of the mature K2 toxin.

Given YKTs often encode several cysteine residues that form stabilizing disulfide bridges, we started with the question which cysteine residues were essential for disulfide bridge formation. The K2 precursor contains a total of 9 cysteines. The alanine scan pointed towards 6 of those being critical for function - located within residues 80-362 - suggesting that three disulfide bridges are formed. One cysteine residue, C300, could be mutated without loss of function (**Figure 2C**). Since disulfide bridges are formed early within the secretory pathway, where Kex2p-dependent proteolytic processing has not yet occurred, we hypothesized that it should be possible to confidently determine disulfide bridge formation in a predicted structure of the K2 precursor. The average pLDDT score of the resulting precursor structure was 79.0 between residues 80 and 362 (**Figure 4A**). Consistent with our pre-proregion predictions, residues 1-79 formed a long extended loop (pLDDT 19.0). Structural analysis subsequently revealed three intrasubunit disulfide bridges, one within the α-subunit (C109-C129) and two within the β-subunit (C315-C320 and C328-C359), with residue C300 not involved in a disulfide bridge (**Figure 4A**). Therefore, the library data regarding cysteines aligned with the predicted structure of the full K2 precursor.

**Figure 4.**
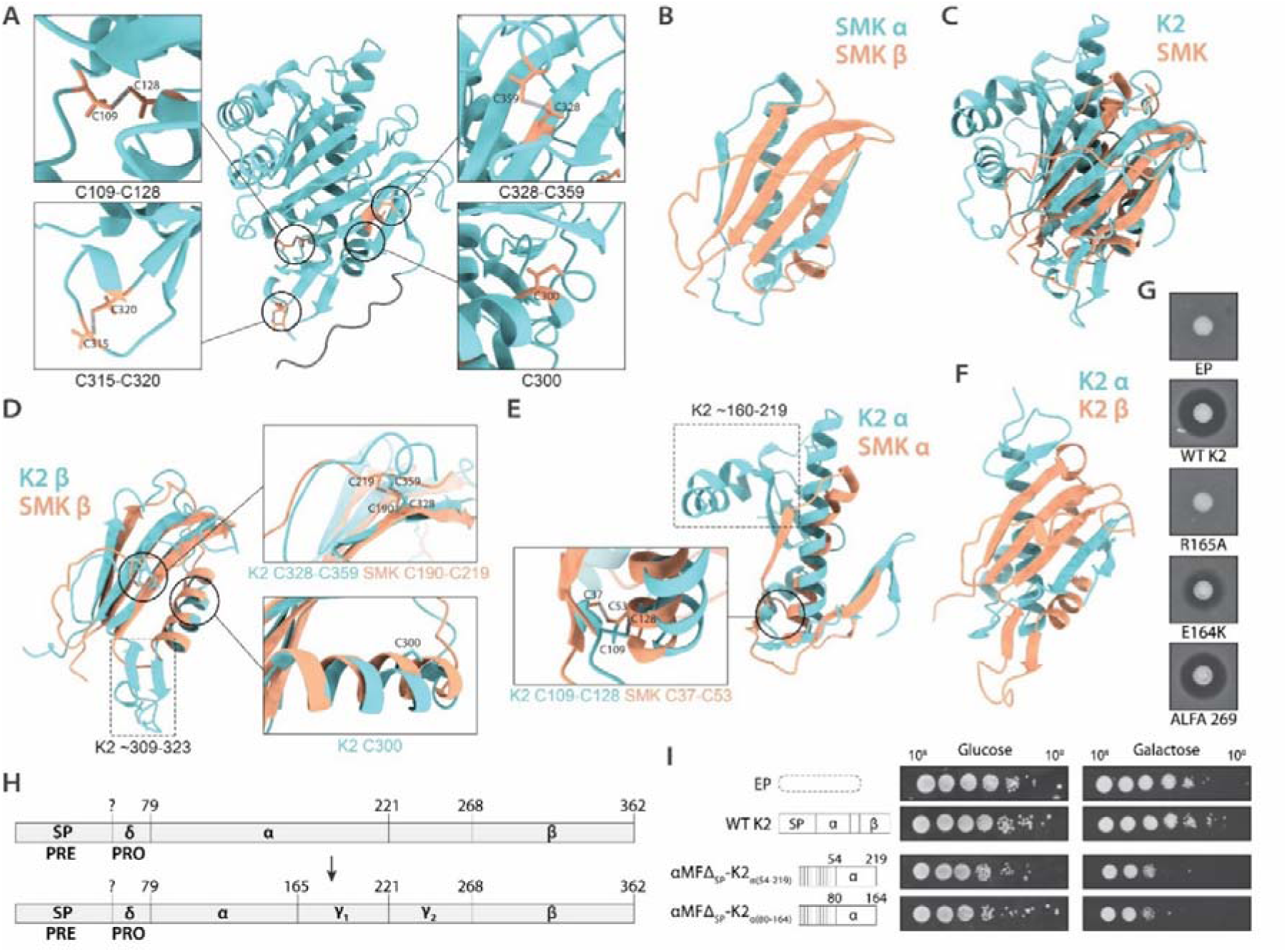
Towards a structural model of the mature K2 toxin. (**A**) The K2 precursor structure was predicted using AlphaFold. For visibility the N-terminus was truncated here up to position 68 (grey). The six essential cysteines form disulfide bridges, whereas the non-essential C300 is not involved in disulfide bridge formation. (**B**) The SMK toxin structure (chain A and B, PDB ID: 1KVE). (**C**) The predicted precursor structure of the K2 toxin (the N-terminus was truncated up to position 80) superposed with the mature SMK toxin structure. (**D**) The β-subunit of the SMK toxin and of the predicted K2 precursor were superposed. While both contain a disulfide bridge connecting the C-terminal loop to the β-sheet, the non-essential C300 residue in K2 is not conserved in the SMK structure. K2 contains an extra loop (marked in the dashed selection) between residues ∼309-323. (**E**) The SMK α-subunit and the K2 α-subunit from the predicted precursor structure were superposed. Cysteine bridges within the α-subunit are highlighted in the box. The K2 α-subunit (here residues 80-219) contains additional residues compared to that of SMK (marked in the dashed selection). (**F**) The predicted mature K2 toxin structure. The α-subunit (residues 80-164) and β-subunit (residues 269-362) form the mature toxin. (**G**) The indicated strains were spotted onto a sensitive background strain to determine toxin activity. EP; empty plasmid. WT K2; wild-type K2. (**H**) The proposed gene organization of the K2 toxin includes a γ-domain. (**I**) The suicidal phenotype of the α-subunit was determined from expression of residues 54-219 or 80-164. Cell cultures normalized for optical density and were diluted 1:10 (∼10^6^-10^0^ cells) and spotted onto plates with non-inducing (glucose) or inducing (galactose) conditions. Plates were incubated for 3 days before imaging.

Interestingly, K2 did not appear to contain intersubunit disulfide bridges connecting the α-subunit and the β-subunit. This was surprising, given several other *S. cerevisiae* dsRNA virus-encoded toxins do contain such intersubunit disulfide bridges, hypothesized to be critical in holding the heterodimer together. For example, the K1 toxin contains a disulfide bridge between C92 (α-subunit) and C239 (β-subunit) [37], and the K28 toxin contains a disulfide bridge between C56 (α-subunit) and C333 (β-subunit) which is reduced once it enters the cytosol to release the cytotoxic α-subunit [38].

Interestingly, the structure of the SMK (or SMKT) killer toxin produced by the halotolerant yeast *Millerozyma farinosa* (formerly *Pichia farinosa*) has been determined experimentally and – similar to K2 – does not contain disulfide bridges in between subunits, but only within its α- and β-subunit, and non-covalent interactions maintain the heterodimer interaction [39]. The SMK precursor consists of 222 amino acids and is organized into an N-terminal preregion followed by α-γ-β-domains. The α-subunit (residues 19-81) and β-subunit (residues 146-222) form the mature heterodimeric toxin (**Figure 4B**) [40]. When we compared the predicted structure of the K2 precursor to the structure of the mature SMK, we noticed remarkable similarities in the fold topology (**Figure 4C**) and the hydropathy profiles of SMK and K2 (**Supplementary Figure S6**). However, the toxins do not share a high sequence identity (**Supplementary Figure S7**).

The overall fold of the SMK β-subunit and the respective region in the K2 precursor (residues 269-362) was very similar, containing a split left-handed βαβ-motif (**Figure 4D**) and we hereafter refer to this region as the K2 β-subunit (see refinement of the β-domain in **Section 7**). Further, both β-subunits contained a conserved stabilizing disulfide bridge at the same position within the structure (C328-C359 in K2, C190-C219 in SMK) (**Figure 4D**). Where K2 encoded the non-critical C300 in the β-subunit α-helix, SMK encoded M173, indicating this non-critical cysteine is also not conserved in the SMK toxin (**Figure 4D**). The K2 β-subunit contained a larger loop between residues 309 and 323 than the SMK toxin, in which an additional disulfide bridge is present (C315-C320). This loop is mainly responsible for the difference in size of the two subunits (the SMK β-subunit consists of 77 residues, the K2 β-subunit counts 94 residues) (**Figure 4D**).

The α-subunit of K2 and SMK also showed similarities, both containing another split left-handed βαβ-motif (**Figure 4E**). The α-subunit provides a central hydrophobic α-helix that is largely shielded from the environment by the surrounding β-sheet (**Figure 4B and C**) (**Supplementary Figure S6**). Our alanine scan had shown that the amino acid sequence of this central hydrophobic helix (in K2 consisting of residues 125-148, residing within sublibrary segments 5 and 6), was highly important for K2 toxicity since this region only contained 3 library variants without reduced functionality (**Supplementary Figure S3**). Similar to the β-subunit, both SMK and K2 contained a conserved disulfide bridge within the α-subunit, formed by residues C37-C53 in SMK and C109-C128 in K2 (**Figure 4E**).

Interestingly, there are also striking functional similarities between SMK and K2. The SMK toxin has an optimum activity at pH 2.5-4.0 which steeply decreases with increasing pH and is completely lost at pH 6.0 [41]. It was suggested that SMK acts by formation of ion channels [41], and later it was demonstrated that it associates with and affects membranes of sensitive cells and induces calcein leakage from liposomes *in vitro* [42]. Overall, these results pointed to a functional and structural relatedness of the two toxins making it reasonable to use the SMK crystal structure as an additional guide to further study the structure of the mature K2 toxin.

### 7. Alanine scanning data together with structure predictions indicates the presence of a γ-domain with a Kex2p cleavage site after R165

While the precursor structure represented the putative structure present in the early secretory pathway, the mature toxin is processed and secreted as an α/β-heterodimer [13]. Next, we sought to generate structure predictions of the mature secreted K2 toxin to understand sequence-function relationships in more detail.

To predict the structure of the mature toxin it is crucial to understand how the precursor is exactly processed within the region 80-362. The K2 precursor contains two dibasic ‘KR’ residues at positions 220-221 and 267-268 that form canonical sites for Kex2p/Kex1p-dependent processing (**Figure 1B**). Consistent with previous data [13], we observed that the R221A toxin variant was not functional (score 1), indicating that cleavage at this KR site is critical for toxin function. K220A showed a minor decrease in toxicity (score 3.5), suggesting that a less efficient ‘AR’ cleavage site suffices to maintain a high activity level but that efficient cleavage is necessary for full activity [33]. In contrast, the R268A variant showed reduced activity (score 2.5) while the K267A variant had wild-type activity (score 4) (**Supplementary Figure S3**), indicating that the R268 cleavage site highly enhances the toxin function, but that the toxin is still active to some extent without it.

Together with the data on a putative proregion up to position R79, we therefore defined that the α-subunit consisted of residues 80 to 219, and the β-subunit of residues 269-362. We used these subunit boundaries and generated a structure prediction of the resulting K2 heterodimer (average pLDDT score of 71.7) (**Supplementary Figure S8A**). However, we noticed a rather low-confidence region within the α-subunit, approximately between residues 160-219 (**Supplementary Figure S8B**). This corresponded to a region within the α-subunit that was also not present in the SMK toxin (**Figure 4E**). We therefore hypothesized that another cleavage site may be present which would lead to removal of this region.

Upon closer inspection of the amino acid sequence, we noticed that residues 164-165 form another putative non-canonical ‘ER’ Kex2p cleavage site. We therefore subsequently generated a structure prediction using α-subunit residues 80-164 and β-subunit residues 269-362 (**Figure 4F**) (**Supplementary Figure S9**). The local confidence of this structure was increased (pLDDT 79.8, pTM 0.815, ipTM 0.847), and the overall topology more similar to the SMK toxin structure (**Figure 4B and F**). In the initial predicted structure of the full K2 precursor (**Figure 4A**), the 164-165 predicted Kex2p cleavage site is situated on a loop between elements of secondary structure and is exposed to the solvent (as well as those around position R79, R221 and R268), increasing the likelihood of this site representing a true Kex2p cleavage site accessible to Kex2p. Unfortunately, the R165A variant was missing in our mutagenesis dataset (**Supplementary Figure S3**). We therefore cloned this variant separately and found that it resulted in loss of function, as expected for an important Kex2p cleavage site (**Figure 4G**). The E164A mutation resulted in wild-type level toxicity (**Supplementary Figure S3**). An E164K mutation resulted in a reduced function, suggesting that this site may prefer an ‘ER’ or ‘AR’ cleavage site over a ‘KR’ cleavage site (**Figure 4G**).

Based on these data, we propose that the K2 α-subunit consists of residues 80-164 (**Figure 4H**). Simultaneously, we propose a γ-subunit stretching residues 166-266 with an internal cleavage site at position 219-220 (resulting in regions γ_1_ and γ_2_). Residues 269-362 form the β-subunit. To further support this model, we confirmed that expression of residues 80-164 fused to the αMF signal peptide resulted in a functional suicidal phenotype, similar to expression of residues 54-219 which were shown previously to generate a cytotoxic α-subunit [17] (**Figure 4I**).

The novel proposed gene organization of the K2 ORF, including a proregion and γ-region, is similar to the SMK and K1 toxin gene organizations (**Figure 5A**). Especially the γ-subunit resembles the one of K1 which also contains an internal cleavage site [32]. However, while the internal cleavage site within the K2 γ-subunit appears to be essential for toxin function, the internal cleavage site within K1 was determined to be nonessential since an R188A mutation still led to functional toxin secretion [32].

**Figure 5.**
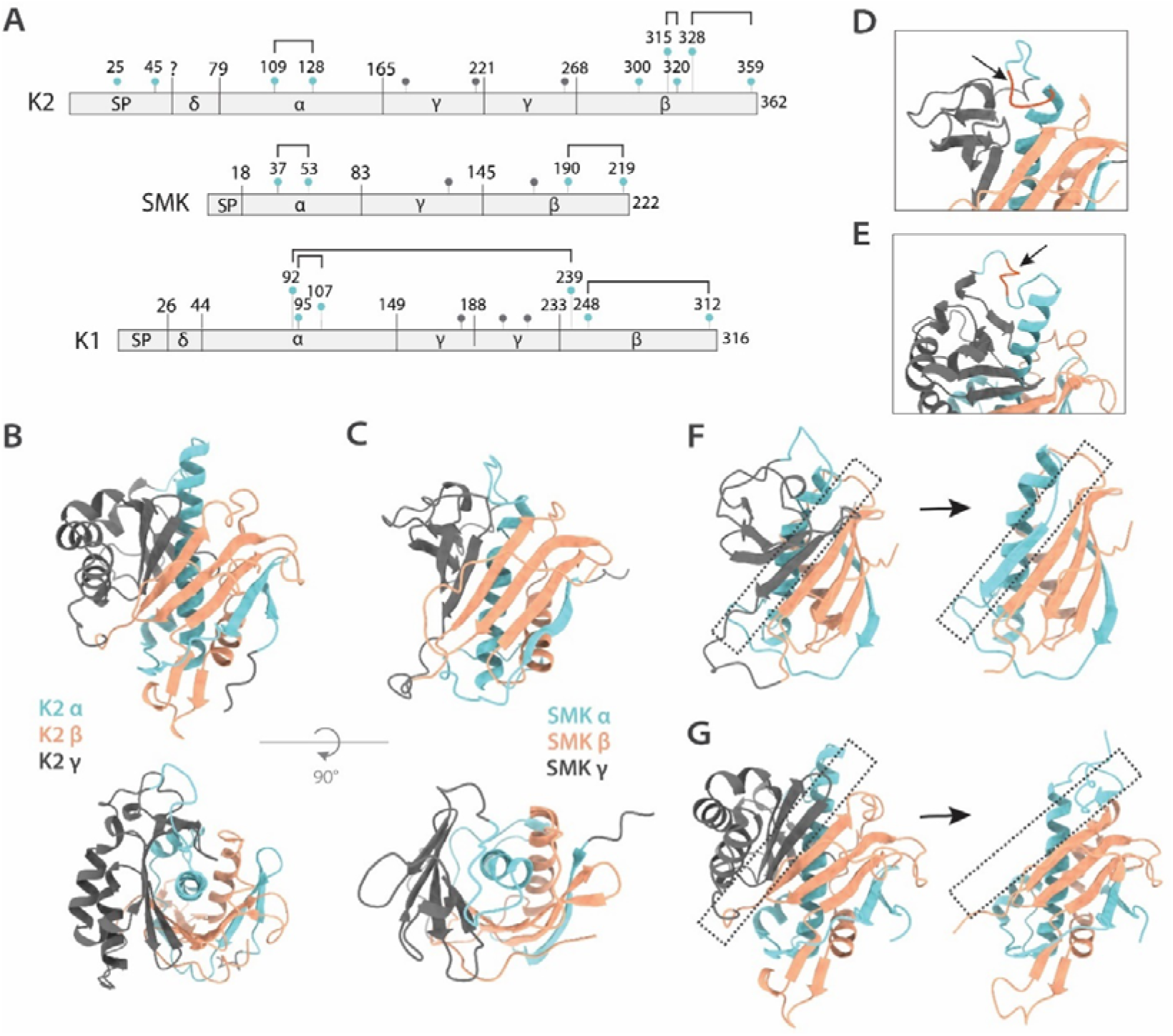
Toxin precursor structures. (**A**) Schematic overview of the K2, SMK and K1 precursor organizations. Potential glycosylation sites are indicated with grey pins, and cysteine residues are indicated with blue pins. Bridges indicate disulfide bridge formation. (**B**) The K2 precursor structure is shown, together with the top view (the preproregion was truncated). (**C**) The SMK precursor structure is shown, together with the top view (the preregion was truncated). (**D**) The GGGG motif in the SMK precursor loop that connects the α-domain to the γ-domain is indicated in red. (**E**) The GGPGG motif in the K2 precursor loop that connects the α-domain to the γ-domain is indicated in red. (**F**) In SMK, the β-strand of the γ-subunit that completes the β-sheet is replaced by a β-strand of the α-subunit after removal of the γ-subunit (indicated by the dashed boxes). (**G**) The γ-subunit in K2 also donates a β-strand to the β-sheet, but in the mature K2 toxin predicted structure we only observed formation of a short β-strand by the α-subunit at the same position.

In addition, the fact that an R268A variant showed reduced toxin activity and not complete loss of activity was suggesting that an N-terminal extension of the β-subunit may be tolerated to some extent. The site may therefore allow the insertion of an epitope tag for toxin purification or localization strategies. We tested this hypothesis by insertion of the ALFA-tag [43] at position 269 in the wild-type sequence, which resulted in a K2 variant that still retained functional toxicity (**Figure 4G**).

### 8. The γ-domain contains critical residues but the N-glycosylation sites are not essential

The proposed γ-domain (residues 166-266) corresponds to sublibrary segments 7 to 10 (which encompass residues 163-268) (**Supplementary Table S1**). Within segments 4 to 16 (corresponding to residues 82-362), the number of variants with functional score 4 was higher in each of the segments 7-10 compared to each of the other segments (**Figure 2A**) (**Supplementary Table S7**). Still, there were residues within this proposed γ-domain that were critical (Y181, Y212, R221, W230 -functional score 1) or strongly important (i.e. F171, L176, L180, G183, Y211, Y242 – functional score 2) for toxin function (**Supplementary Figure S3**), potentially by impacting proper folding and maturation of the toxin precursor, indicating that the γ-domain plays a critical role in function, rather than being a simple linker.

For the K1 toxin it had been determined that the γ-domain contains 3 potential N-glycosylation sites, but that the final mature toxin does not contain glycosylation [34] (**Figure 5A**). For the K74 toxin from *S. paradoxus*, which shares a similar gene organization, the secreted heterodimer was also determined to lack glycosylation - all glycosylation sites are also present within the excised γ-domain [36]. There are also three potential N-glycosylation sites within the K2 precursor at positions N177, N214 and N261 (**Figure 5A**). According to the novel proposed organization of the K2 ORF, these N-glycosylation sites would all be present within the γ-domain, similar to K1 and K74, in which case we would not expect the mature K2 toxin to be glycosylated. For the K1 toxin, it was further determined by mutagenesis that none of the three N-glycosylation sites within the γ-domain were required for either toxicity or immunity [34]. In our data for the K2 toxin, we similarly found that the N177A and N261A variants resulted in wild-type activity (score 4). N214A resulted in only a slight loss of activity (score 3.5). Although data from a previous study suggested glycosylation of one of the toxin domains [13], the data here indicated that (at least individual) glycosylation of these positions was not critical for toxin function.

### 9. SMK precursor reconstruction shows further similarities with the K2 precursor structure and indicates conformational reorientation of the **α**-subunit C-terminus after excision of the **γ**-domain

Within the mature SMK toxin structure - determined in 1997 - the C-terminus of the α-subunit and the N-terminus of the β-subunit are close together (**Figure 4B**), which led the authors to hypothesize that the γ-subunit forms an independent domain next to the mature SMK toxin subunits [39]. AlphaFold now gives the opportunity to also reconstruct the SMK precursor structure.

In the resulting structure (average pLDDT of 81.5), we indeed observed that the γ-subunit was predicted to be located on one side of the precursor structure, similar to the conformation in the K2 precursor (**Figure 5B and C**). The γ-subunit of SMK (61 amino acids) is smaller than the γ-subunit from K2 (101 amino acids). While the γ-subunit of SMK only consisted of β-strands, the γ-subunit of K2 consisted of both α-helices and β-strands. From the top view, the central hydrophobic helix from the α-subunits is visible, with the γ-domains located on one side of the proteins (**Figure 5B and C**).

These structures can help to reconstruct the further sequence of events in precursor processing towards the mature toxin, after excision of the γ-domain. While most of the structure appears to stay nearly identical between the precursors and the mature toxins, one region visibly undergoes changes: The loop that connects the α-domain to the γ-domain (**Figure 5D and E**). The SMK toxin encodes a GGGG motif within this region (G72-G75) (**Figure 5D**, in red), and the K2 toxin encodes a similar GGPGG motif (G157-G161) (**Figure 5E**, in red) (**Supplementary Figure S7**). Mutation into alanine of several of these residues resulted in reduced toxin functionality (GGPGG had functional scores 3, 2, 3, 4, 3, respectively) (**Supplementary Figure S3**), indicating that the region performs a function for K2 toxicity. Being present in the connecting region between the α-domain and γ-domain, we speculate that this motif may provide a required flexibility in the loop connecting the two domains (**Figure 5D and E**).

The γ-subunit is arranged in such a way that it is ready for cleavage within the secretory pathway (**Figure 5B and C**). Prior to cleavage, within the SMK precursor, the γ-subunit provides a β-strand that completes the β-sheet formed by the α-subunit and β-subunit (**Figure 5F**, left panel). Comparing the SMK precursor structure and the mature SMK toxin structure, it appears that this β-strand is replaced by a β-strand formed by the C-terminus of the α-subunit after the γ-subunit is proteolytically removed (**Figure 5F**). The K2 γ-subunit also provided a β-strand to the β-sheet in the predicted precursor structure, but the K2 α-subunit C-terminus only formed a small replacing β-strand in our structure predictions of the mature toxin while the remainder of the C-terminus was pointing into the solution (**Figure 5G**).

### 10. Molecular dynamics simulations confirm overall stability of the predicted mature K2 toxin structure and find a more stable conformation of the α-subunit C-terminal region

To confirm that the predicted structure of the mature K2 toxin is stable in solution, and to assess if the C-terminus of the α-subunit could possibly find a more stable conformation, we performed molecular dynamics (MD) simulations. As a complementary dataset, we also performed the simulations using the SMK toxin, since the structure was experimentally determined at acidic pH (pH 3.5 [39]) it should be stable in MD simulations.

While the SMK structure and the predicted K2 mature toxin structure present the overall fold of the toxins, the protonation states do not reflect those expected at an acidic pH - where the toxins are active - because hydrogen atoms are missing in the models. Therefore, we used the PROPKA algorithm [44] to assign protons to the titratable amino acid side chains of SMK and K2. This is important, as it was shown for the SMK toxin that protonated side-chains of glutamic acid and aspartic acid residues can participate in the hydrogen bond network and may play a role in pH-dependent toxin stability [39].

We performed three independent MD simulations of 500 ns in the protonated state as expected at pH 4.0. The radius of gyration (Rg) and number of hydrogen bonds over time were both stable, indicating that the structures overall remained folded and compact (**Supplementary Figure S10**), further supporting the accuracy of the predicted mature K2 toxin structure. For the K2 toxin, we observed some fluctuation in the Rg and the root mean square deviation (RMSD). The root mean square fluctuation (RMSF) indicated that there were regions with more conformational flexibility, in particular the C-terminus of the K2 α-subunit (**Supplementary Figure S10A-C, I and J**).

We inspected the trajectory further, here displaying 33 superposed frames that were extracted equidistantly in time from each generated trajectory between 20-500 ns (**Figure 6A and B**). These frames create a representation of the sampled conformations over the course of the trajectory. The SMK toxin conformation remained highly stable over time, as expected (**Figure 6A**). While the overall structure of K2 also remained stably folded, we observed that the C-terminal loop of the K2 α-subunit behaved differently across the three replicates (**Figure 6B**). In replicate 1, the loop spent a majority of the time in a position on top of the β-sheet formed by the β-subunit (**Figure 6B**, r1). In replicate 2, the loop was mostly folded back onto itself and remained positioned closer to the initial conformation in the middle (**Figure 6B**, r2). In replicate 3, the loop moved towards a position closer to the conformation as observed in SMK, positioning parallel to the β-sheet formed by the β-subunit (**Figure 6B**, r3).

**Figure 6.**
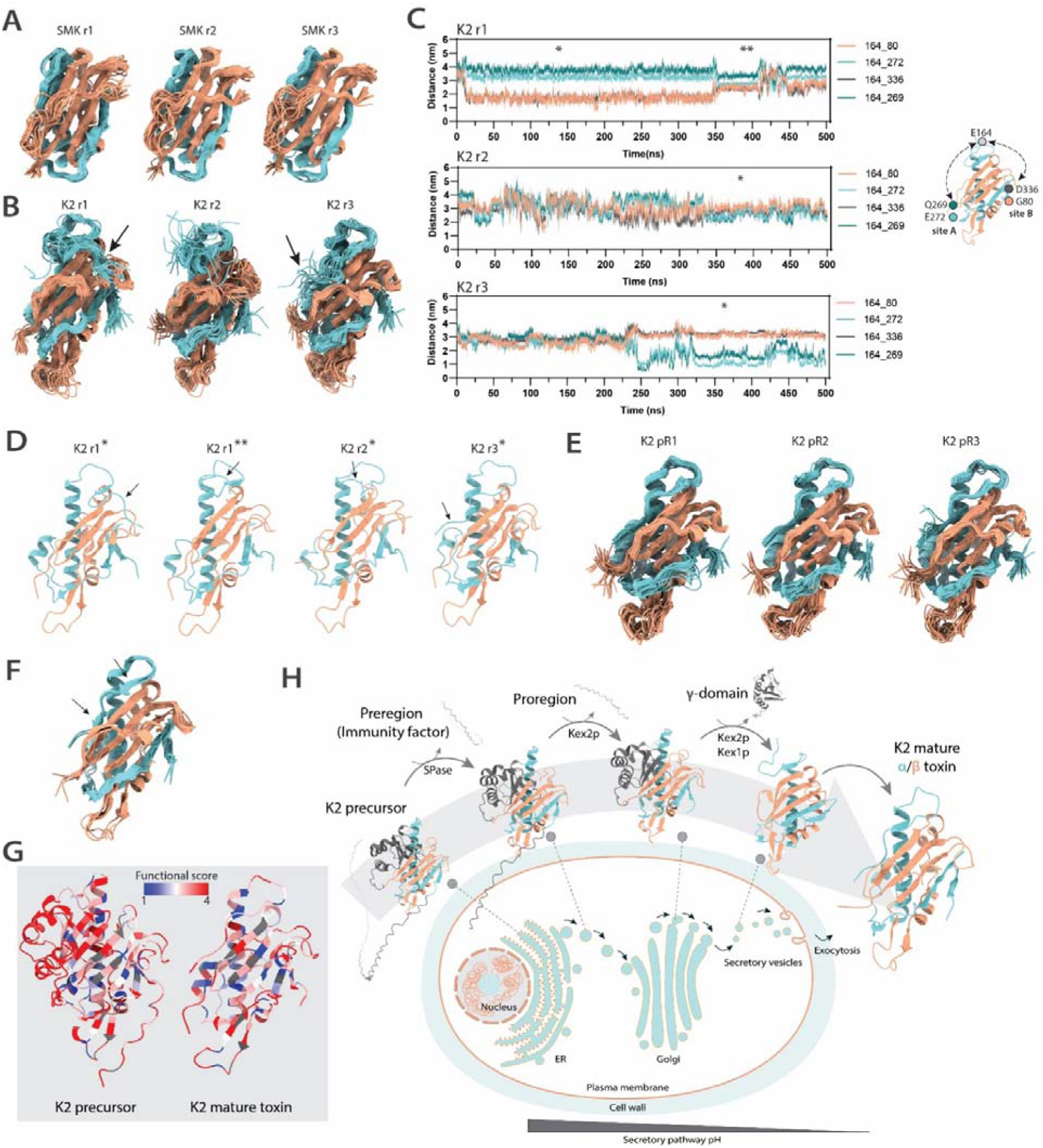
Molecular dynamics of mature toxin structures. (**A**) Displayed are 33 superposed frames from each replicate trajectory of the SMK toxin (r1-r3), equidistant in time, between 20-500 ns. (**B**) Displayed are 33 superposed frames from each replicate trajectory of the K2 toxin (r1-r3), equidistant in time, between 20-500 ns. The arrow indicates the position of the C-terminal region of the α-subunit (**C**) The distances between residue 164 and residues 269 and 272 (‘site A’) and 80 and 336 (‘site B’) are shown over time in the three replicates. The timeframes of cluster centroids are indicated by asterisks. (**D**) Cluster centroids from K2 r1-r3 are shown: *, largest cluster, centroid; **, second largest cluster centroid. The arrow indicates the location of the C-terminal region of the α-subunit. (**E**) Displayed are 29 superposed frames from each replicate trajectory of the K2 toxin with altered protonation state (pR1-pR3), equidistant in time, between 20-300 ns. (**F**) The centroids of the largest clusters from pR1-3 were superposed, two fragments of β-sheet formation are indicated by arrows. RMSD: 1.155 ± 0.085 Å. (**G**) Functional scores visualized on top of the precursor and the mature toxin structure. Residues with score 0 – not present in the library-were colored grey. (**H**) Overview of K2 processing from the precursor to the mature K2 toxin. Within the ER, the signal peptide (preregion) is cleaved off and forms the immunity factor, protecting the cell against the K2 toxin. Within the Golgi, the Kex2p and Kex1p proteases cleave at several processing sites, releasing the proregion and the γ-domain. The resulting α- and β-subunits form the mature secreted K2 toxin, after the C-terminal region of the α-subunit adopts a position parallel to the β-sheet formed by the β-subunit.

We also observed these movements when determining the distance between the Cα atom of residue E164 (the end of the α-subunit) and the Cα atoms of residues Q269 or Q272 (site ‘A’), or between E164 and residues D336 and G80 (site ‘B’) (**Figure C**). While the loop in replicate 1 rapidly found the position, moving towards site B, the position was not stable, since around 350 ns it moved back towards the initial starting position more in the middle of site A and site B (**Figure 6C**, r1). For replicate 2, the loop pivoted around the initial position, remaining relatively in the middle of site A and site B (**Figure 6C**, r2). In the trajectory of replicate 3, the loop moved towards site A around ∼250 ns where it remained located for the remainder of the trajectory (**Figure 6C**, r3). Subsequent cluster analysis established a number of groups with similar structures, from which the centroids were extracted. These centroid structures of the largest clusters (the timeframe of which are indicated with * or, for a second-largest cluster, with **, in **Figure 6C**) show representative structures of these conformations (**Figure 6D**).

While the position of the α-subunit C-terminal region in replicate 3 towards the end of the trajectory was more similar to the conformation observed in the SMK toxin (**Figure 4B**, **Figure 6A**), there was still fluctuation in the position (**Figure 6C**, r3, **Supplementary Figure S10J**). We reasoned that this could be due to the protonation state. When we used the initial predicted mature K2 toxin structure as input for the protonation state prediction by PROPKA (**Figure 4F**), D270 and the C-terminal end of residue E164 were predicted to be unprotonated, and thus carried a negative charge in this MD simulation. Since the environment of E164 had changed due to the conformational change, and since residue D270 seemed potentially located close to the putative new location of the α-subunit carboxyl-terminus, we hypothesized that the initially predicted pKa may no longer be accurate for these residues. To potentially allow for formation of more stabilizing interactions within this region, we used the centroid structure from replicate 3 as input (**Figure 6D**, r3*) for another set of 300-ns triplicate MD simulations, here protonating D270 and the carboxyl-terminus of E164. The Rg, RMSD, and number of hydrogen bonds rapidly converged in these trajectories and remained stable throughout the rest of the simulation (**Supplementary Figure S11**).

We inspected these trajectories further, here displaying 29 superposed frames that were extracted equidistantly in time from each generated trajectory between 20-300 ns (**Figure 6E**). These frames show that the α-subunit C-terminal region, located parallel to the β-sheet formed by the β-subunit, is highly stable over the course of the timeframes. We also obtained the centroid structures of the main clusters, and superposed these centroid structures, indicating their high similarity (**Figure 6F**).

We can now project the functional scores resulting from the alanine scanning library on top of the K2 precursor structure and the K2 mature toxin structure with the MD-adjusted conformation (**Figure 6G**). Structure analysis shows insertion of the E164 residue between W273 (functional score 1) and Y352 (between which there is a pi-interaction). M162 inserts into a hydrophobic pocket and potentially stabilizes the loop between residues 110-122 (which had a relatively lower pLDDT score in the initial predicted structure (**Supplementary Figure S9B**)). Fragmented β-strand formation is established by backbone interaction between residues Q155 and E281 and between residues G161 and Y276 (**Figure 6F**). We observed that the sidechain of residue D270 remains pointing towards the solution, and as such the protonation of this residue arguably had less impact on structure stability. The functional score of the D270A mutation was 4, indicating that the functional group could be converted without loss of function.

We had noticed above that the E164A mutation was functional but that the E164K conversion led to reduced toxin functionality as determined by halo assay (**Figure 4G**, **Supplementary Figure S3**). While the E164A mutation retains the A164 as the C-terminal residue, the lysine residue resulting from the E164K mutation would in principle be removed by Kex1p after Kex2p cleaves after R165 within the secretory pathway, causing the α-subunit to shorten by one residue and to end with residue A163. The A163 residue would likely be unable to reach the same site occupied by E164. Thus, the potential stabilizing role of E164 or A164, rather than the cleavage site efficiency, may dictate the functional effect observed from the E164K mutation. In case Kex1p would fail to remove the lysine residue, while conversion of the protonated glutamic acid residue to alanine retains a neutral sidechain, this would lead to introduction of a positive charge. Such positively charged residue may also not be well tolerated at this position. While the K2 precursor is processed at several sites, the R165 cleavage site is the only site where the new C-terminus remains present in the mature K2 toxin and exposes a new C-terminal residue with a potential function in toxin stability or action.

We therefore propose that the K2 mature toxin structure is topologically more similar to SMK than the initial prediction indicated (**Figure 4F**). In fact, we speculate that the initial structure prediction (**Figure 4F**) may represent an early conformation of the α/β-toxin after proteolytic excision of the γ-domain. After the γ-domain is removed, the C-terminus of the α-subunit adopts the site which was previously occupied by the γ-domain, a process represented by the here obtained MD trajectory (a rearrangement that can occur within ∼250 ns (**Figure 6B**, r3)) (**Figure 6H**).

### 11. Further linking the sequence-function map to the toxin structure models

With the sequence-function map and 3D-models of the K2 precursor and mature toxin structures, further linkage of the sequence-function and structural data is now possible (**Figure 6G**). We had found several library variants that showed complete loss of function (**Figure 2C**). We so far have shown that the critical six cysteines are involved in disulfide formation within the α- and β-subunits, and that R79, R165 and R221 are critical for proteolytic processing. We observed that residues Y181 and Y212 (functional score 1), and W273 and Y350 (functional score 1 and 3) form pi-interactions that may be important for structure stability (similar to Y211 and Y242, both functional score 2). G345 may be critical because it is located right next to the C328-C359 bond that locates the C-terminus of the β-subunit close to the β-sheet in which G345 is located, potentially causing steric hinder when mutated to alanine. G144 is also located in a packed area, potentially also leading to steric hinder when mutated into alanine. Several other critical hydrophobic residues are buried and may perform important stabilizing functions in hydrophobic contacts.

## Discussion

While yeast killer toxins hold potential as antimicrobials, lack of sequence-function and structural data of these often multi-step processed proteins limits engineering strategies to overcome their sometimes inherent limitations. Here, we used the K2 toxin from *S. cerevisiae* as a model and generated an alanine scanning-based sequence-function map which revealed key residues for toxin action, gain-of-function variants, and variants with altered target specificity. Together with the library data, subsequent genetic analysis and structure predictions revealed previously overlooked proteolytic processing sites, leading to the structure prediction of the mature K2 toxin, which has a similar gene organization as the K1 toxin and a similar fold topology as the SMK toxin. Molecular dynamics simulations further refined the mature toxin structure and indicate how the α-subunit C-terminus rearranges after the excision of the γ-domain.

We report several mutations that led to increased zones of inhibition, in particular at positions 293 and 357. A few other reports of naturally occurring killer toxin variants also indicated enhanced functions. A K1 killer toxin variant (BJH001) with the two mutations I103S and T146I increased the cytotoxicity against the yeast *Kazachstania africana* [45]. A few previous reported K2 variations also influenced the activity spectrum [14]. These data indicate that a limited number of mutations can alter the toxicity to specific species of yeast. While several mutations at positions 293 and 357 resulted in enhanced activity against *S. cerevisiae,* these variants had not been reported in natural isolates yet. Perhaps in nature, toxicity towards an array of yeast species (i.e. wild-type K2 spectrum includes *N. glabratus*) is favored over high toxicity towards the own species (i.e. the L293A variant shows increased activity against *S. cerevisiae*, but little against *N. glabratus*) (**Supplementary Figure S5**). These data suggest room for engineering of natural occurring variants towards increased activity specifically against pathogenic or spoilage yeast. Alternatively, with more sequenced dsRNA virus genomes more natural variants with altered activity ranges may be detected [1]. The data presented here show that site-directed mutagenesis libraries can further provide mutants with enhanced functions, even by single-amino acid conversions.

We further defined the processing of the K2 precursor with cleavage sites at R79 and R165, which had so far been overlooked [46]. Because killer toxins can use non-canonical Kex2p cleavage sites, but are dependent on a P1 arginine, we found it is a useful approach to mutate the P1 site to determine its essentiality. Within residues 60-362, K2 encodes 10 arginine residues, with the following functional scores after alanine conversion: R63 (score 4), R79 (score 1), R165 (score 1), R221 (score 1), R224 (score 4), R244 (score 4), R264 (score 4), R268 (score 2.5), R287 (score 4), R318 (score 3.5). These data point to the potential involvement of R79, R165, R221, R268 in important processing sites. It has been previously proposed that K2 may also contain a disulfide bridge between the α- and β-subunit [20,46], and that K2 does not contain a γ-domain, or that the γ-domain is not released [47]. However, we here presented evidence that indicate no intersubunit disulfide bridge between the α- and β-subunit, similar to the SMK toxin, and that there is a γ-domain which is processed and released.

There is increasing evidence for the existence of killer toxin families. For the K1 toxin it has recently been shown that there is a family of biologically active K1-like killer toxins [48]. We here found that the K2 precursor and mature toxin structure showed similarities to the SMK toxin from *M. farinosa* [40,41], which has been described as an emerging human pathogen in immunocompromised patients [49]. The SMK toxin (genome encoded) has further been compared to the monomeric KP4 toxin from *Ustilago maydis* (dsRNA virus encoded), a calcium channel inhibitor, showing that the folding topology is highly similar (**Supplementary Figure S12**) [39]. It was noted that the SMK and KP4 structures contain two left-handed split βαβ-motifs, which are rare in other proteins [39]. The K2 model shares this structural motif with SMK and KP4. It has further been noted that K2 and the K66 toxin from *Saccharomyces paradoxus* (dsRNA virus encoded) both contain a DUF5341 domain [47], which is found only in *Ascomycota* yeast and in mycoviruses. We found that the predicted structure of the mature heterodimeric K66 toxin also shares fold similarities with K2, containing a hydrophobic central α-helix surrounded by a β-sheet (**Supplementary Figure S12C**). While the SMK toxin does not rely on β-1,6-glucans (a *kre1* deletion mutant is sensitive) [39], K66 has been shown to interact with β-1,6-glucans similar to K2 [47]. It will be interesting to elucidate the evolutionary and/or functional relationships between these killer toxins.

Besides the interest in YKTs as antifungals, the study of the synthesis, structure, processing and secretion of yeast killer toxins have also strengthened our understanding of essential eukaryotic cellular processes such as post-translational protein processing within the secretory pathway, the pathways involved in production of YKT cell wall receptors such as β-1,6-glucan, and virus-host interactions [50]. As such, several genes are named after killer toxins, for example the *MAK* (maintenance of killer) genes and the *SKI* (super killer) genes, and *KRE* (killer resistant) genes [50]. The K2 toxin here indicates how regions of the protein adjacent to excised domains may adjust to a new position after Kex2p-dependent cleavage of those domains. Since many eukaryotic proteins are processed by Kex2p or homologs (such as furin) within the secretory pathway, such proteins may show similar dynamics.

One limitation of yeast killer toxins is their dependence on often low pH conditions. The SMK α-subunit is relatively hydrophobic (**Supplementary Figure S6**), has low solubility in medium and precipitates at neutral pH [40]. Containing no intersubunit disulfide bonds, the subunits dissociate under neutral pH conditions [39,40]. The β-subunit loses secondary structure and the toxin cannot be regenerated [51]. Such inability to regenerate functional toxin after incubation at elevated pH has also been shown for the K2 toxin [15], suggesting that similar molecular factors may be underlying this phenotype. The SMK structure indicated that the pH-dependent stability of the heterodimer may be the result of interactions between carboxyl groups of protonated aspartic acid and glutamic acid residues, which act stabilizing under acidic conditions [39]. Under neutral conditions, these groups would carry negative charges, causing electrostatic repulsion between the sidechains that destabilize the structure [39]. In particular, D76-D213 (intersubunit), D161-D195 (β-subunit), and E158-D199-D222 (β-subunit) were determined to be involved in such interactions [39]. The K2 mature toxin structure provides a first step towards analysis of pH-dependency, potentially facilitating a strategy towards protein engineering of stability in a broader pH range.

The pH difference between ER (pH ∼7.4) and relatively acidic Golgi (pH ∼5.8), towards the secretory vesicles (pH ∼5.5) can influence the association or dissociation of proteins [52,53]. It is currently unclear how pH-dependent YKTs can maintain a stable conformation within the secretory pathway, where initially pH conditions exist that would render the mature toxin inactive and potentially unstable. Speculatively, the proregion and γ-domain play a role in stabilizing the structure towards the late Golgi where the pH is sufficiently lowered (**Figure 6H**). Further studies on the toxin precursor compared to the mature toxin might reveal insights into potential additional functions of the γ-domain besides linking the α- and β-subunit.

The data presented here describe several novel findings related to the K2 killer toxin and can provide useful information for follow-up site-directed mutagenesis experiments. We further showed that an epitope tag can be tolerated at the N-terminus of the β-domain, providing a tool for future studies. Ultimately, a better understanding of relationships between genotype and phenotype of yeast killer toxins, combined with structural data, is needed, together with a better understanding of the molecular processes underlying the killing mechanism. This study shows, using K2 as a model, that recent developments in high-throughput oligonucleotide synthesis combined with library screening, high-throughput sequencing and protein structure prediction can help to study multi-step processed proteins, an important and universal process happening in the secretary pathway which was before the structure-prediction age only accessible for very few proteins with structural information available about precursors or intermediates [54,55].

## Materials and Methods

### Strains

For plasmid constructions and maintenance of all constructs *Escherichia coli* DH5α was used (Thermo Scientific, #18265017). Recombinant proteins were expressed in the yeast strain *Saccharomyces cerevisiae* BY4741 (MAT*a leu2*Δ*0 met15*Δ*0 ura3*Δ*0 his3*Δ*1*) (derived from ATCC, 4040002) [56]. The BY4741 Δ*pbs2* deletion mutant with increased sensitivity to the K2 toxin [57] was obtained from the yeast gene deletion collection (Thermo Scientific, 10277124) and used as a sensitive background strain in halo assays. *Nakaseomyces (Candida) glabratus ySB040* (clinical isolate, gift from Dr. Daniel Green, Columbia University Medical center) was used for experiments with *N. glabratus*.

### Materials

Media and buffer components were obtained from BD Bioscience (Franklin Lakes, NJ, USA) and Sigma Aldrich (Darmstadt, Germany). Sterile, transparent round-bottom microtiter plates were obtained from Corning (Corning Inc.). Black clear-bottom 96-well microtiter plates were obtained from Thermo Scientific. A Singer ROTOR benchtop robot was used for transfer and re-plating of arrayed yeast cultures and colonies. One-well ROTOR PlusPlates and Singer RePads for robotic transfer were obtained from Singer (SingerInstruments).

### Culture conditions and transformation

*E. coli* cultures were incubated at 37°C/200 rpm in LB medium, which was supplemented with chloramphenicol (25 µg/mL) or ampicillin (100 µg/mL) were necessary. Solid media were supplemented with 2% agar and incubated at 37°C. Preparation and transformation of chemically competent *E. coli* were performed using standard CaCl_2_ and heat-shock procedures.

Before transformation, *S. cerevisiae* cultures were grown at 30°C on plates containing YPD (1% yeast extract, 2% peptone, and 2% glucose). Transformants were obtained using the LiAc-PEG protocol [58]. Transformants were subsequently selected and cultivated in minimal Synthetic Defined (SD)-AS/Urea media (SD), which unless otherwise mentioned lacked histidine [59]. For selection and maintenance, SD media with 2% glucose was used as the carbon source. For assay conditions, cells were grown in SD media with 1% sucrose as the carbon source, 0.5% galactose was added for induction of protein expression, and media was buffered with 0.5x of McIlvaine buffer at pH 4.6 as described earlier [59]. The K2 toxin is most active in acidic conditions and at a temperature of 25°C [15]. Cells were either grown on solid media supplemented with 1.6% agar, in 3 mL liquid media within glass culture tubes, or arrayed in 96-well microtiter plates containing 200 µL of liquid medium. One-well ROTOR plates were filled with 30 mL of solid media.

### Cloning of GFP-dropout entry vectors

All plasmids and primers used within this study are listed in **Supplementary Table S9** and **S10**, respectively. In a previous study, the K2 ORF was divided into 16 segments (**Supplementary Table S1**), and each segment was replaced by an SbfI-restriction site, resulting in plasmids pRP035-pRP050 [17]. These plasmids contain a pRS423-type high-copy backbone, and a galactose-inducible promoter. To create the mutagenesis library we used a method based on oligonucleotide pools and Golden Gate cloning [22]. First, we generated entry vectors by using the SbfI-sites to insert a GFP-dropout module (**Figure 1C**). For this purpose, plasmids pRP035-pRP050 were digested with the SbfI restriction enzyme and treated with alkaline phosphatase (Thermo Scientific, EF0654). A GFP-dropout module, which can be expressed in *E. coli*, was amplified from plasmid pYTK047 from the Yeast ToolKit (Addgene, Kit#1000000061) [26]. DNA amplification was performed using 2X Phusion Hot Start II HF PCR Master Mix (Thermo Scientific, F565L). The primers used for this amplification are segment-specific, and add a Type II restriction enzyme (BsaI) site to the GFP-dropout module. DNA purification was performed using the GenElute PCR Clean-Up Kit (Sigma-Aldrich). The amplicons and the digested plasmids (∼50 ng) were assembled using GeneArt Gibson Assembly HiFi Master Mix (Thermo Scientific, A46627) for 1 hour at 50°C, resulting in plasmids where the K2 segments were replaced by a GFP-dropout module (pRP139-pRP154). The reactions were transformed into *E. coli*, green colonies were selected and the plasmids were purified using the GenElute Plasmid Miniprep Kit (Sigma-Aldrich). The plasmid sequences were verified by Sanger Sequencing.

### Oligonucleotide pool design

Oligonucleotide pool sequences encoding the alanine scanning variants are listed in **Supplementary Table S11**. All positions (including the start codon, excluding the stop codon) were converted to alanine (GCT) (guided by codon usage frequency). Native alanine residues were converted into glycine (GGT). Segment sequences were ordered from Integrated DNA Technologies (IDT) as single-stranded oPools at a 10 pmol/oligo scale, one for each of the sixteen sublibrary segments. In addition, oligonucleotide pools were designed for saturation mutagenesis of positions 293 and 357 (**Supplementary Table S11**). The oligonucleotide length varied from 69 to 119 nucleotides between segments and as such, the pools contain around ∼75% to ∼65% full-length products, respectively, according to the manufacturer’s website.

### Oligonucleotide pool double-stranding

Given the high sequence similarity among oligos within the mutagenesis pools, the pools were only double-stranded instead of amplified to prevent cross-overs leading to undesired products [22]. The lyophilized oligo pools were dissolved in milliQ water to a concentration of 1 µM and double-stranded in two 50 µL reactions using 2X Phire Green Hot Start II Master Mix (Thermo Scientific, F126L) and the primers listed in **Supplementary Table S10**. For the reaction, 2.5 µL of each 10 µM primer stock, and 2.5 µL of the 1 µM oligo pool were used and the reactions were subjected to the following parameters on a thermal cycler: (1) 98°C for 5 s, (2) annealing for 10 s at the appropriate temperature, (3) 72°C for 10 s, (4) repeat 1-2-3 another cycle, (5) 72°C for 30 s, for each pool (**Figure 1E**). For each oligo pool, the two 50 µL reactions were pooled together and DNA was purified using the GenElute PCR Clean-Up Kit, eluting in 30 µL. The primers also add sequences to the oligonucleotides such that each 5’ and 3’ end contains BsaI-restriction sites that create overhangs compatible with scarless and in-frame ligation into the appropriate entry vector.

### Library assembly

To assemble the mutagenesis libraries, a 25 µL Golden Gate reaction was prepared for each sublibrary which contained 20 ng of the appropriate entry vector, 10 µL of the double-stranded oligo pool, 2.5 µL of the T4 ligase buffer, 1 µL of the T7 DNA Ligase (NEB, M0318S), and 1 µL of the BsaI-HF®v2 restriction enzyme (NEB, R3733S) (**Figure 1F**). The reaction mixtures were incubated in a thermal cycler with the following parameters: (1) 42°C for 2 min, (2) 16°C for 5 min, (3) repeat steps 1-2 up to 25 cycles, (4) 60°C for 10 min, (5) 80°C for 10 min. The 25 µL reactions were each transformed in 125 µL of competent *E. coli* DH5α cells, which were plated on selective media to obtain single colonies. Per segment, all white colonies were counted and scraped together and added into 500 µL of LB media supplemented with ampicillin and glycerol. Part of the cell suspension was inoculated into 3 mL LB supplemented with ampicillin and incubated overnight at 37°C/200 rpm, while the remainder was stored at -80°C. From the overnight culture, plasmids were extracted using the GenElute Plasmid Miniprep Kit and eluted in 30 µL. Library completeness was estimated using the GLUE webserver tool (https://guinevere.otago.ac.nz/STATS/glue.php) [60].

### Library transformation into yeast and arraying

250 ng of each sublibrary plasmid pool was transformed into *S. cerevisiae* BY4741. The transformation was plated on selective media to obtain single colonies. Individual colonies were picked and arrayed in 200 µL media (SD media with 2% glucose and 15% glycerol) in 96-well microtiter plates. The first column of each 96-well plate was dedicated to control strains: Row 1, 3 and 5 contained strains producing wild-type K2, row 2 and 4 contained non-killer strains producing GFP (facilitating visual confirmation of plate orientation), and row 6-8 contained previously created strains with K2 variants that result in varying halo sizes (**Figure 1H**). The plates were incubated overnight at 30°C, the cultures were used as sources for the killer toxin assays, and subsequently stored at -80°C.

### Killer toxin activity assays

The 96-well arrayed yeast strains were assayed for secretion of active toxin in a halo assay. The liquid cultures from the 96-well plates were pinned onto solid agar plates (SD media with 1% sucrose, 0.5% galactose, buffered to pH 4.6) in technical duplicates using the ROTOR robotic pinner (**Figure 1I**). After 3 days, halo assay plates were prepared by seeding a sensitive strain (BY4741 Δ*pbs2*), cultured overnight in the same media composition, in fresh agar plates with the same media composition (∼7.5*10^7^ sensitive cells in 30 mL of media - assuming approximately 2*10^7^ cells per OD_600_ unit per mL). The colonies from each solid source plate were then replicated onto these assay plates and the plates were incubated at 25°C for 2 days before imaging zones of inhibition using a FUJI imager (**Figure 1J**).

### Analysis of zones of inhibition

Active toxin production was identified by the formation of a zone of growth inhibition. To quantitatively compare the activity of the different variants, we used image-processing software originally developed for quantification of colony luminescence – CFQuant, created by Dafni et al. [27]. However, the software relies on the color gradient of the luminescent halos to help handle image noise, but the zones of inhibition do not display such a gradient. Therefore, a modified version of the software was used (version 1.7, see Data availability), which includes extra options to handle image noise based on the position of the colonies and the circular shape of the zones of inhibition. The modified software allowed quantification of areas of the zones of inhibition, and the sizes of colony areas were also recorded (**Figure 1K**). To account for plate-to-plate variation, the areas of the zones of inhibition of the three reference wild-type K2 expressing colonies of each plate were averaged, the other areas of zones of inhibition of the respective plate were normalized to the average wild-type K2 halo size on that plate, and then averaged between the two technical duplicates.

We set thresholds for strong effects (i.e. no zone of inhibition) to no effects (i.e. wild-type size) of the mutations on toxin activity. We performed a statistical analysis to set the range of wild-type, which meant taking all the three halos of each plate of the reference wild-type K2 producers into consideration, and establishing what we did not consider to significantly deviate from wild-type (100%) activity. We used this data to estimate the error distribution of wild-type activity, and estimated a 99% confidence interval (CI) on average of 84.5-115.5% of the wild-type toxin halo size for what represented wild-type activity. We then set 80-120% of the average wild-type halo area size as conservative lower and upper thresholds for what is considered wild-type activity. Second, we set a upper threshold of 10% for the variants with the strongest loss-of-function effect, allowing for some biological variation. Two bins were set in between to assess positions with major to moderate effects. The library variants were therefore grouped based on the quantified halo area sizes relative to the wild-type into four bins: Bin 1 (≤10% of the original halo size), bin 2 (10-40%), bin 3 (40-80%) and bin 4 (>80%) (**Figure 1L**). A number of mutants with gain-of-function effects (halo area size >120%) were identified and studied separately.

### Plasmid library extraction from yeast

The genotypes and phenotypes were linked using next-generation sequencing. The K2 ORF of 1089 bases was divided into 5 amplicon regions that could each be sequenced with 250 bp paired-end reads (**Figure 1M**). Amplicon 1 covered sublibrary segments 1-3, amplicon 2 sublibrary segments 4-6, amplicon 3 sublibrary segments 7-9, amplicon 4 sublibrary segments 10-12, and amplicon 5 sublibrary segments 13-16. In this way, these amplicons targeted the site of mutagenesis only, such that these regions could be sequenced to high coverage depth. All yeast colonies belonging to the same bin were pooled together per sublibrary segment into 300 µL of 2% glucose SD media supplemented with 20% glycerol, in such a way that approximately similar amounts of biomass of each colony were added. Of each cell suspension, 100 µL was subsequently inoculated into 3 mL of SD medium with 2% glucose and grown overnight at 30°C, 200 rpm, while the remainder was stored at -80°C. From these overnights, the OD_600_ was determined, and the cultures belonging to the same amplicons and bins were subsequently inoculated together into 20 mL of fresh media as a first multiplexing step, such that the added inoculum was normalized for the number of colonies that the sample represented. This 20 mL culture was again incubated overnight at 30°C, 200 rpm in SD medium containing 2% glucose as the carbon source. The next day, 10 mL of the cultures were centrifuged (2500x *g*, 5 min) and the supernatant was aspirated. For each plasmid extraction, the cell pellet was resuspended in 500 µL of the resuspension buffer of the GenElute Plasmid Miniprep Kit. The samples were kept on ice while the cells were lysed using acid-washed glass beads, with bead beating performed in two cycles at 6 m/s for 20 s (Savant Bio 101 FastPrep FP120 bead mill). 300 µL of the suspension was used for a plasmid extraction following the Plasmid Miniprep Kit manufacturer’s instructions and plasmid DNA was eluted in 30 µL elution buffer.

### Illumina library samples preparation

For primer barcode design, first a set of 407 barcodes of 8 bp was selected (available at http://hannonlab.cshl.edu/nxCode/nxCode/Ready_made_sets.html), and an optimized set of 10 barcode sequences was selected using BARCOSEL (http://ekhidna2.biocenter.helsinki.fi/barcosel/ [61]), with a minimal pairwise Levenshtein sequence distance of 4. PCR-mediated partial Illumina adapter and 8 bp-barcode addition and library amplification were performed using 1 µL of the purified plasmid library as a template in a 50 µL reaction using the 2X Phusion Hot Start II HF PCR Master Mix, with the following parameters for the thermal cycler: (1) 98°C for 30 s, (2) 98°C for 10 s, (3) 56°C for 20 s, (4) 72°C for 1 min, (5) 2-3-4 for 20 cycles, (6) 72°C for 5 min (**Figure 1M**). The samples were purified using the GenElute PCR Clean-Up Kit and eluted in 20 µL elution buffer. Fluorometric quantification of the amplicons was performed with the Quant-iT PicoGreen dsDNA Assay Kit (Invitrogen) in a Synergy Mx plate reader (BioTek). For the fluorometric quantification, 1 µL of amplicons was diluted in 99 µL of 1x TE buffer (10 mM Tris-HCl, 1 mM EDTA, pH 7.5) and mixed with 100 µL of the PicoGreen agent (diluted 1:200 in 1x TE) in a black transparent-bottom 96-well microtiter plate. Standards were prepared following the manufacturer’s instructions. Based on these measurements, the amplicons were further multiplexed into five final sequencing samples. The amounts of amplicons were added proportionally to the plate colony numbers in order to maintain an equimolar representation, with the aim of maintaining equal read distribution for all colonies in the downstream sequencing run as much as feasible.

### Sequencing

The samples were sequenced by Genewiz (Azenta Life Sciences, Leipzig, Germany) using the Amplicon-EZ service (250 bp paired-end reads) on an Illumina Miseq sequencing platform (**Figure 1N**). Independent validation of a set of individual colony sequences was performed by Sanger sequencing (Macrogen).

### Bioinformatics analyses of sequencing reads

Illumina sequencing returned two FASTQ files containing the forward and reverse sequencing reads and quality information. The initial quality profiles of the reads were visualized with FastQC (v0.11.9 [62]). Sequences were preprocessed in several steps (**Figure 1O**), where default parameters were used unless specified otherwise. Paired-end reads were demultiplexed using the 5’ 8 bp-barcodes on forward and reverse reads and assigned to the original segment/bin combinations (allowing one mismatch; –barcode-mm 1) (AdapterRemoval v2.3.3 [63]), and for downstream analysis this barcode was also trimmed at this step. Reads with poor quality were removed and low-quality bases were trimmed from the paired end reads (Trimmomatic [64] using SLIDINGWINDOW:4:20, MINLEN:100). The paired-end reads were subsequently merged into a single read (ea-utils fastq-join, v1.3.1 [65]). The quality of merged reads was assessed with FastQC. The reads were mapped to the reference template of pRP002 using BWA-MEM (v0.7.17 [66]) – of the reads that passed the previous filters, all were mapped - and the alignment files were sorted using SamTools (v1.18 [67]). SamTools indicated good mapping quality (MAPQ alignment score 60). The resulting files were subsequently parsed by a custom python script.

Since the sequencing read covers the entire mutated region, the read from this amplicon is by design representative and sufficient for variant calling and counting. The sequencing results were analyzed to obtain the occurrences of each amino acid conversion. Because of the need to analyze each read as a whole instead of single nucleotide substitutions, we developed a custom script for analysis of the alignment file. The custom script parses through the alignment file and performs stringent quality filtering to retain high-quality reads and assigns counts to the detected variants in the context of the entire read. An alanine mutation is counted when the programmed substitution is present and any additional amino acid variations are absent throughout the entire read. The results are aggregated and merged into excel files for further analysis.

### Functional score calculation

Quantitative data of read counts were normalized for differences in the total read counts of the five different sequencing samples. Overrepresented outliers were determined by applying a mathematical criterion for outliers (the third quartile (Q3) + 1.5 times the interquartile range (IQR)). Subsequently, for each amino acid position in the K2 ORF, the read counts across bin 1-4 were compared. We set the following thresholds for determining the functional score: 1) a minimum threshold for the coverage of 10 variant read counts, together with either 2) a minimum of 80% of all variant reads of that position detected within one bin, or 3) a minimum of 80% of all variant reads of that position was present in two adjacent bins. If criteria 1) and 2) were met, we assigned a functional score that is equal to the bin number (i.e. a variant detected with >80% of the reads in bin 3 was assigned a functional score of 3). If criteria 1) and 3) were met but not criterium 2), we assigned a functional score of the average of the two adjacent bins (i.e. if a variant was detected in bin 3 and 4 is was assigned the score 3.5). If criterium 1) was not met, we excluded the reads from further analysis and the position was assigned a score of 0. As such, the variants obtained a functional score between 1 and 4 with intervals of 0.5 points, or score 0.

### Construction of double mutant L293A + G357A

A construct containing both L293A and G357A mutations was generated by amplification of the respective regions from the two isolated plasmids pRP120 and pRP121 using primers RP01 and RPA25 (**Supplementary Table S9 and S10**). The pRP002 plasmid backbone was digested with EcoRI (Fast Digest, Thermo Scientific, FD0274) and BamHI (Fast Digest, Thermo Scientific, FD0054), and the PCR products were digested with either EcoRI and ApaLI (NEB, R0507) (containing the L293A mutation) or ApaLI and BamHI (containing the G357A mutation). All products were purified from an agarose gel using the GenElute Gel Extraction Kit (Sigma-Aldrich) and ligated in a 20 µL ligation reaction using T4 ligase. After transformation into *E. coli* and subsequent plasmid purification from a monoclonal culture, the sequence was verified by Sanger sequencing.

### Manual halo assay

To assess secretion of active toxin, separate variant colonies were picked and inoculated in 3 mL of SD media (2% sucrose, 1% galactose, pH 4.6) and incubated at 25°C at 200 rpm for 16h. A sensitive strain (*S. cerevisiae* BY4741 Δ*pbs*2 [57]) was also included. The OD_600_ of all overnight cultures was determined and adjusted to 10. Of these cell suspensions, 5 µL (∼10^6^ cells) was spotted onto an agar plate seeded with 125 µL of the OD_600_ 10 culture per 10 mL of solid media (same media composition) of the sensitive strain (∼2.5·10^7^ cells). Plates were incubated at 25°C for 2 days before imaging. ImageJ (v1.52a) was used for quantification of these halo sizes. For enhanced visibility, the brightness and contrast of the images was adjusted equally.

### Spot assay

To assess suicidal phenotypes, separate variant colonies of the strains of interest were picked and inoculated in 3 mL of SD media (2% sucrose, pH 4.6) and incubated at 25°C at 200 rpm for 16h. The OD_600_ of all overnight cultures was determined and adjusted to 10. In a transparent round-bottom 96-wells microtiter plate, a 10-fold dilution series from the OD_600_ 10 culture was prepared in sterile water. From each dilution, 5 µL was spotted onto two target plates: The first one containing 2% glucose as a carbon source (repressing condition), the second buffered to pH 4.6 and containing 2% sucrose as a carbon source and supplemented with 1% galactose (inducing condition). The plates were incubated at 30°C or 25°C, respectively, for 3 days before imaging the growth. For enhanced visibility, the brightness and contrast of the images was adjusted equally.

### Protein structure predictions

Using AlphaFold (v2.3.1) [24,25] and ColabFold (v.1.5.2) [23] (installed locally), several protein structure predictions of monomers or multimers were generated. ColabFold was run with increased recycle counts (up to 50) and seeds (up to 25) (using models alphafold2_ptm or alphafold2_multimer_v3). Models were ranked by pLDDT, pTM and ipTM scores. Top ranked structures were relaxed using the AMBER colaboratory from ColabFold with default settings (relax_amber.ipynb) [23]. UCSF ChimeraX (v1.6.1) and Pymol were used for analysis and visualization of the predicted protein structures.

### Molecular dynamics simulations

All simulations were prepared using GROMACS (v2023.1) with the CHARMM36m all-atom force field [68]. Since the pKa of titratable residues can shift depending on the local environment, pKa values were estimated using PROPKA (v.3.5.1) [44], with default parameters. The protonation state of the proteins was adjusted to reflect conditions at pH 4.0. The proteins were placed into dodecahedral boxes, which a minimal distance of 1.0 nm between the protein and the box boundaries. For the K2 system, the box dimensions were 3.62 x 5.26 x 4.31 nm, and for SMK, they were 4.18 x 3.33 x 3.77 nm. The systems were solvated with TIP3P waters (K2: 11.780 molecules; SMK: 5.699 molecules). To neutralize the system and achieve a 0.15 M NaCl concentration, a corresponding number of water molecules was replaced with Na^+^ and Cl^-^ ions.

All simulations used parameters recommended for the CHARMM force field. The leap-frog integrator was used with a 2 fs time step. Neighbor searching utilized the Verlet cutoff scheme, and van der Waals (Lennard-Jones) interactions were smoothly switched off between 1.0 to 1.2 nm. Coulomb interactions were computed using the Particle-Mesh Ewald (PME) method, with a real-space cutoff of 1.2 nm and a grid spacing of 0.16 nm. The V-rescale thermostat maintained the temperature at 298 K with a time constant of 0.1 ps, while the Parrinello-Rahman barostat maintained pressure at 1 bar using isotropic scaling and a time constant of 2 ps. Bond lengths involving hydrogen atoms were constrained with the LINCS algorithm.

After system construction, the potential energy of the system was minimized using the steepest descent method. This was followed by 100 ps of position-restrained equilibration in the NVT ensemble and 100 ps of equilibration in the NPT ensemble. Once temperature and pressure equilibration were achieved, unrestrained production runs of 500 ns were generated (298 K, 1 bar) with a 2 fs time step, and trajectory data were recorded every 10 ps. Trajectory analysis was conducted using GROMACS built-in tools, with hydrogen bonds defined by a distance cutoff of 3.5 Å and an angular cutoff of 30°.

## Author Contributions

RCP and SB designed the study. RCP conducted the experiments. RCP and SB analyzed the data. TM developed the custom python script for NGS read analysis. ED and IY developed and modified the image analysis software for quantifying the zones of inhibition. RCP wrote the manuscript. SB edited the manuscript.

## Data availability

The data that support the findings of this study are available from the corresponding author, SB, upon reasonable request. The custom variant calling script used in this manuscript is available at https://github.com/Tmarinus/aa-mutation-counter. The software used for quantifying the zones of inhibition (CFQuant version 1.7) is available at https://github.com/eyaldaf/CFQuant.

## Acknowledgements

We thank the Center for Information Technology of the University of Groningen for their support and for providing access to the Hábrók high performance computer cluster. We also thank Michael Chang (European Research Institute for the Biology of Ageing, Groningen, the Netherlands) for providing access to the Singer ROTOR benchtop robot for manipulation of arrayed yeast cultures and colonies.

