## Supplementary information for "Alanine-scanning of the yeast killer toxin K2 reveals key residues for activity, gain-of-function variants, and supports prediction of precursor processing and 3D structure"

This file contains the Supplementary Figures and Tables.

**Supplementary Figures**

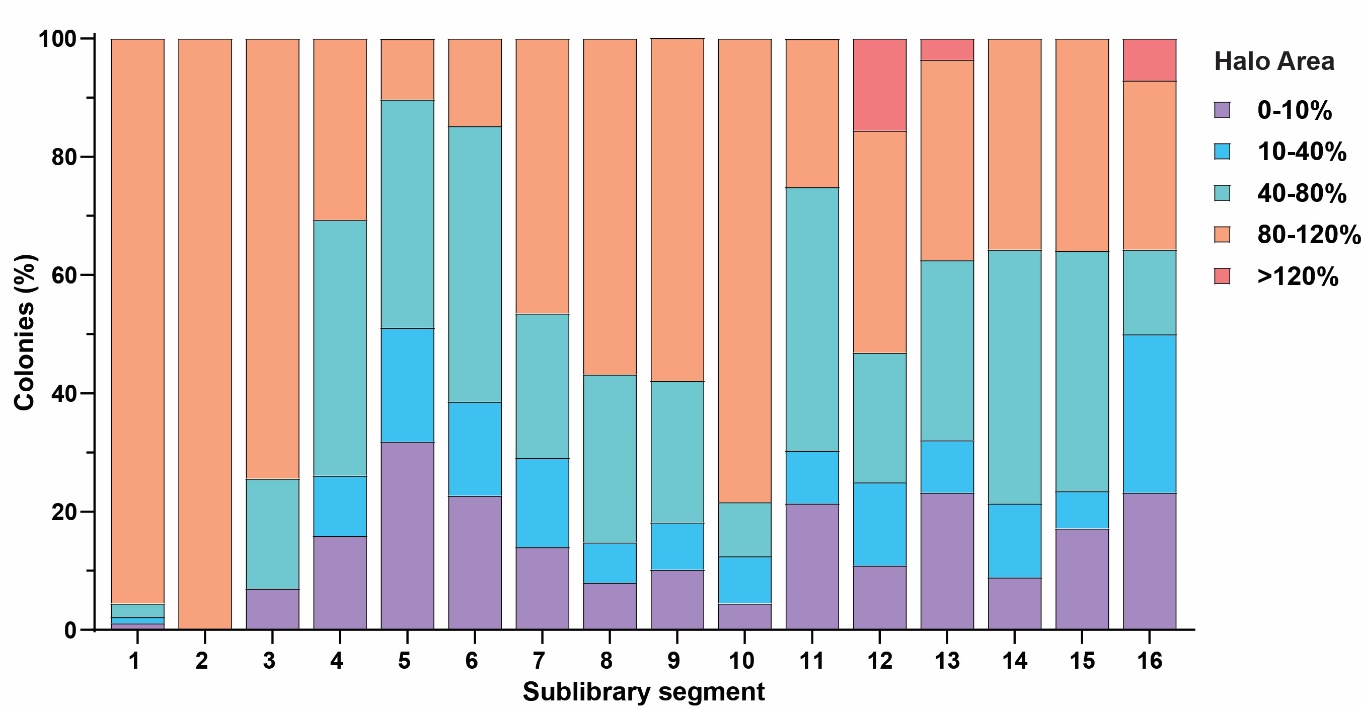

**Supplementary Figure S1. Distribution of the zone of inhibition area sizes of colonies across K2 sublibrary segments.** For each colony, the area of the halo size was quantified relative to the wild-type and averaged across two replicate plates. The colonies were subsequently divided into bins, from 0-10% (bin 1), 10-40% (bin 2), 40-80% (bin 3) to >80% of wild-type activity. The graph shows the percentage of colonies per sublibrary segment in each bin. Here, we also show an additional bin which contains gain-of-function colonies with halo sizes >120% of the wild-type (in red).

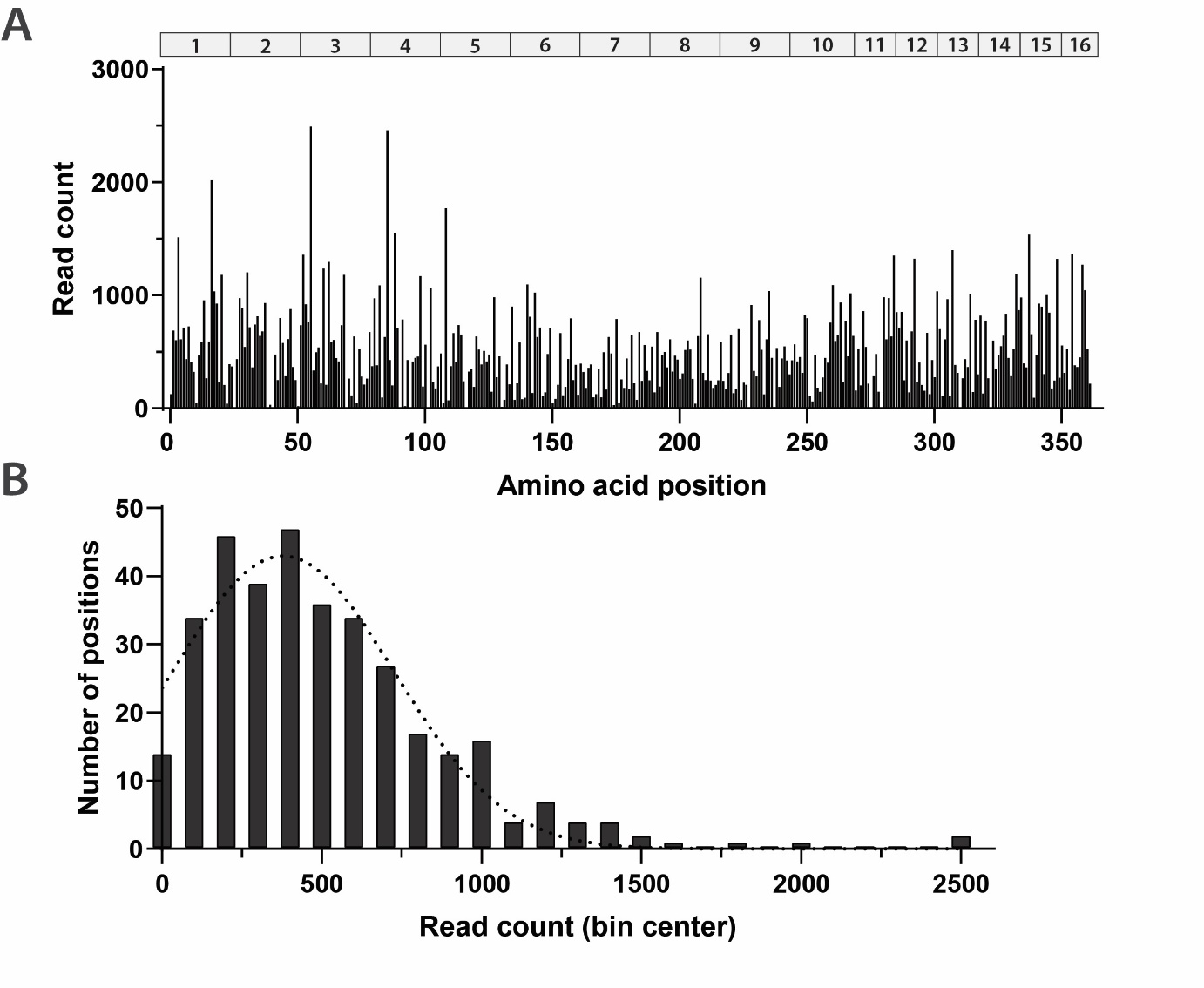

**Supplementary Figure S2. Distribution of NGS read counts.** (**A**) After reads processing, the number of reads for each single, intended mutation per amino acid position was counted. The library segments are displayed on top of the graph for reference. (**B**) The read counts per variant followed a Gaussian distribution (indicated by the dotted curve). The 25% percentile was 243, the median was 443 and the 75% percentile was 684. The mean read count was 515 reads, with a standard deviation of 374 reads.

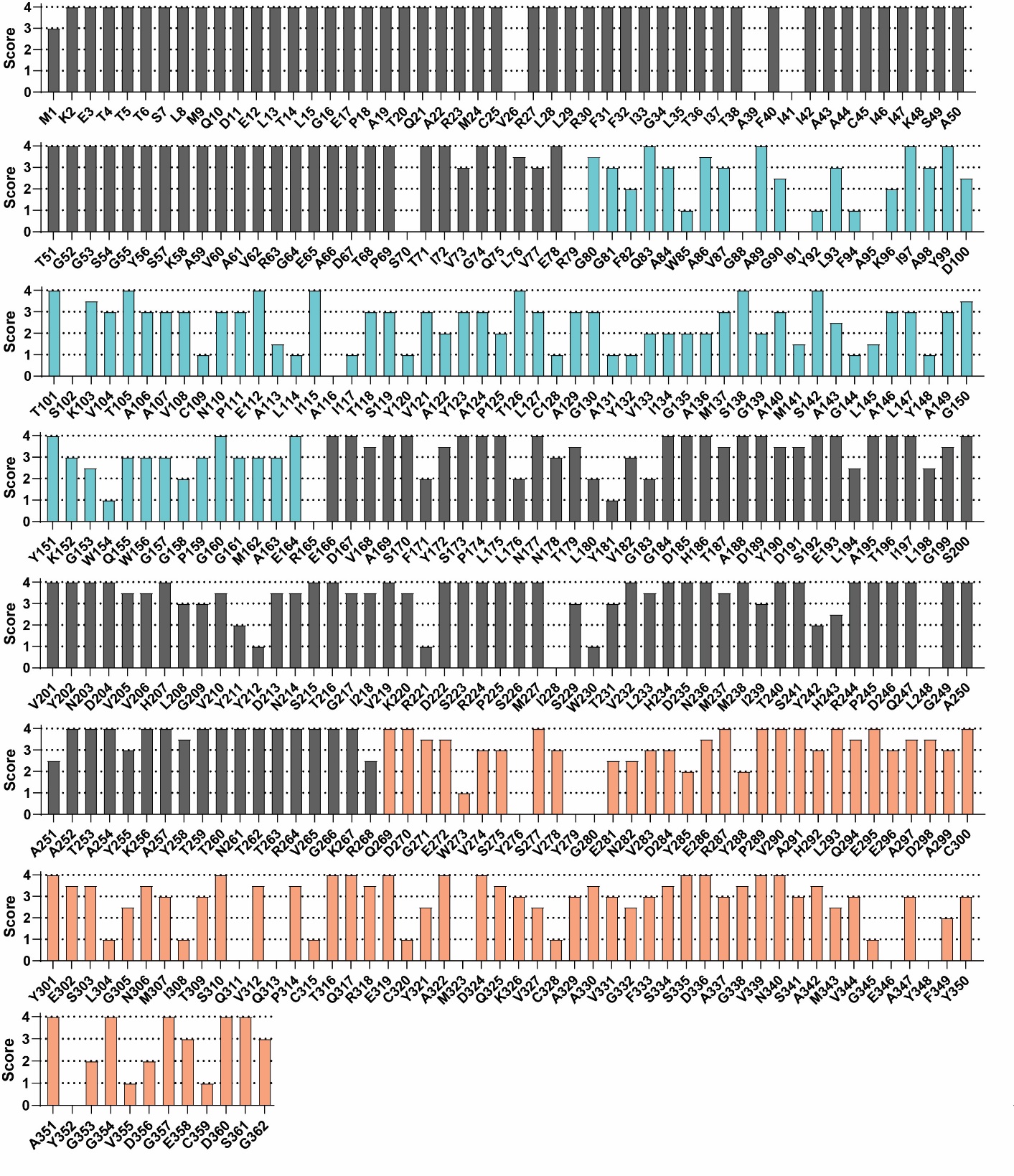

**Supplementary Figure S3. Detailed per-residue sequence-function map.** The functional score - representing the effect of amino acid conversion into alanine (or alanine into glycine) - is displayed for each position in the K2 ORF. The score runs from functional score 1 (loss-of-function) to functional score 4 (wild-type activity). For residues with score 0, the data did not meet the quality standards and the positions were excluded from analysis. The proposed α-domain is indicated in blue, and the β-domain in orange.

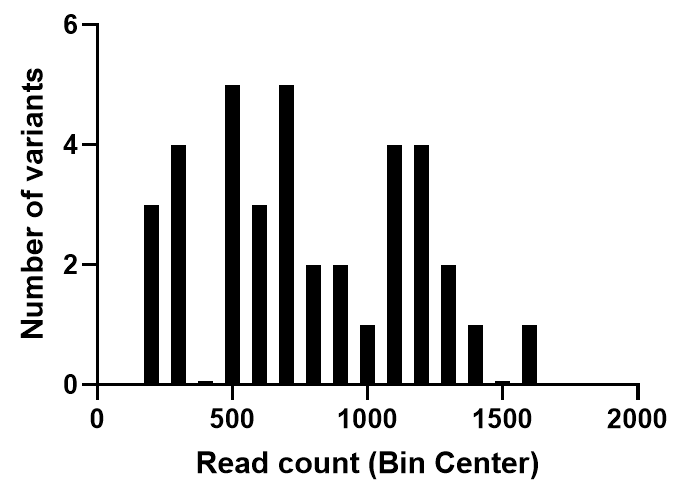

**Supplementary Figure S4. Read distribution for saturation mutagenesis mutants of positions L293 and G357.** After reads processing, the number of reads for each single, intended mutation per amino acid position was counted. The read counts per position varied from 157 to 1636, with an average of 754 and a median of 697 reads.

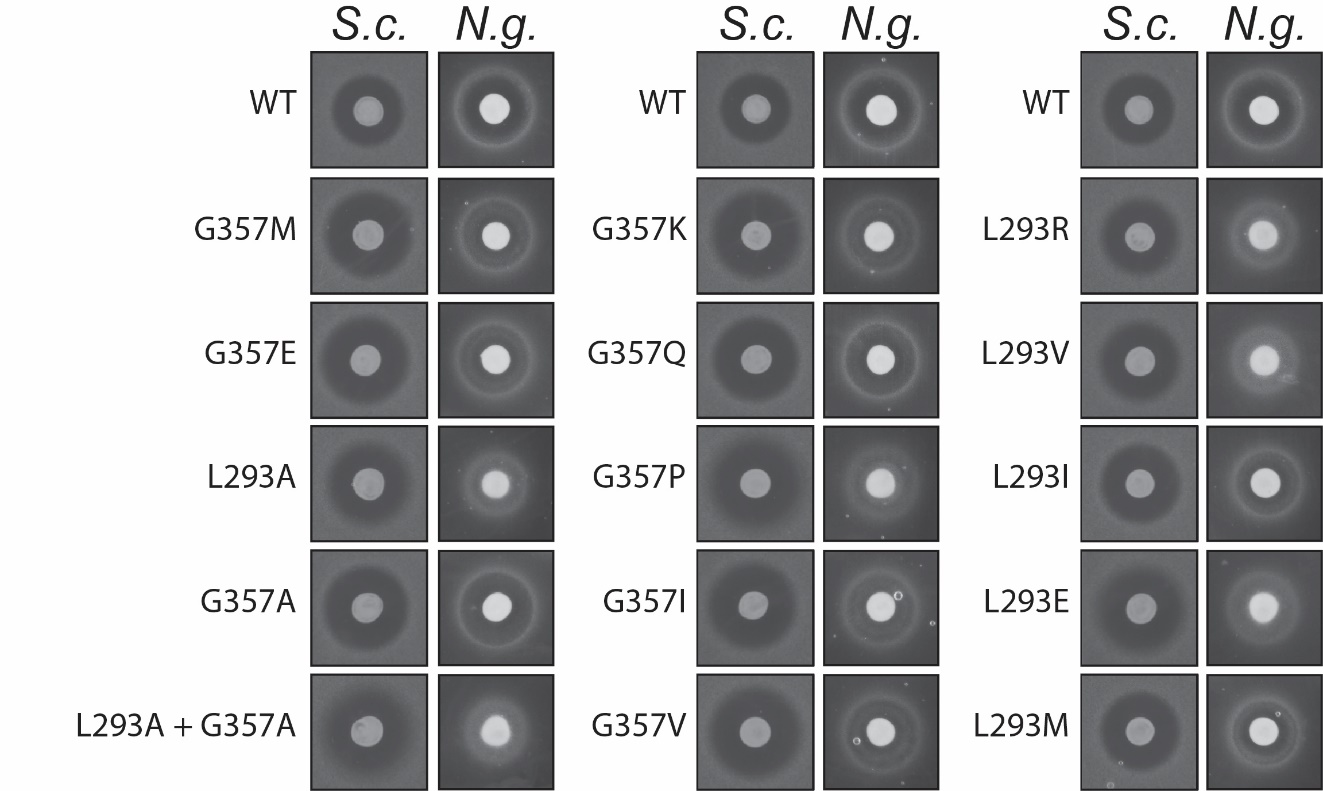

**Supplementary Figure S5. Individual halo assays for saturation mutagenesis variants of positions L293 and G357.** Plasmids from several colonies from the saturation mutagenesis libraries were extracted and reintroduced in yeast. These transformants were spotted on top of a lawn of sensitive background strain, which was either *S.c*. ; *Saccharomyces cerevisiae* (Δ*pbs2*) or *N.g. ; Nakaseomyces glabratus.* The plates were incubated for 2 days before zones of inhibition were imaged. The images are representative of duplicate experiments. Some of the tested variants resulted in zones of inhibition with a more diffuse border against *S. cerevisiae* (i.e. L293A+G357A, L293E, G357P).

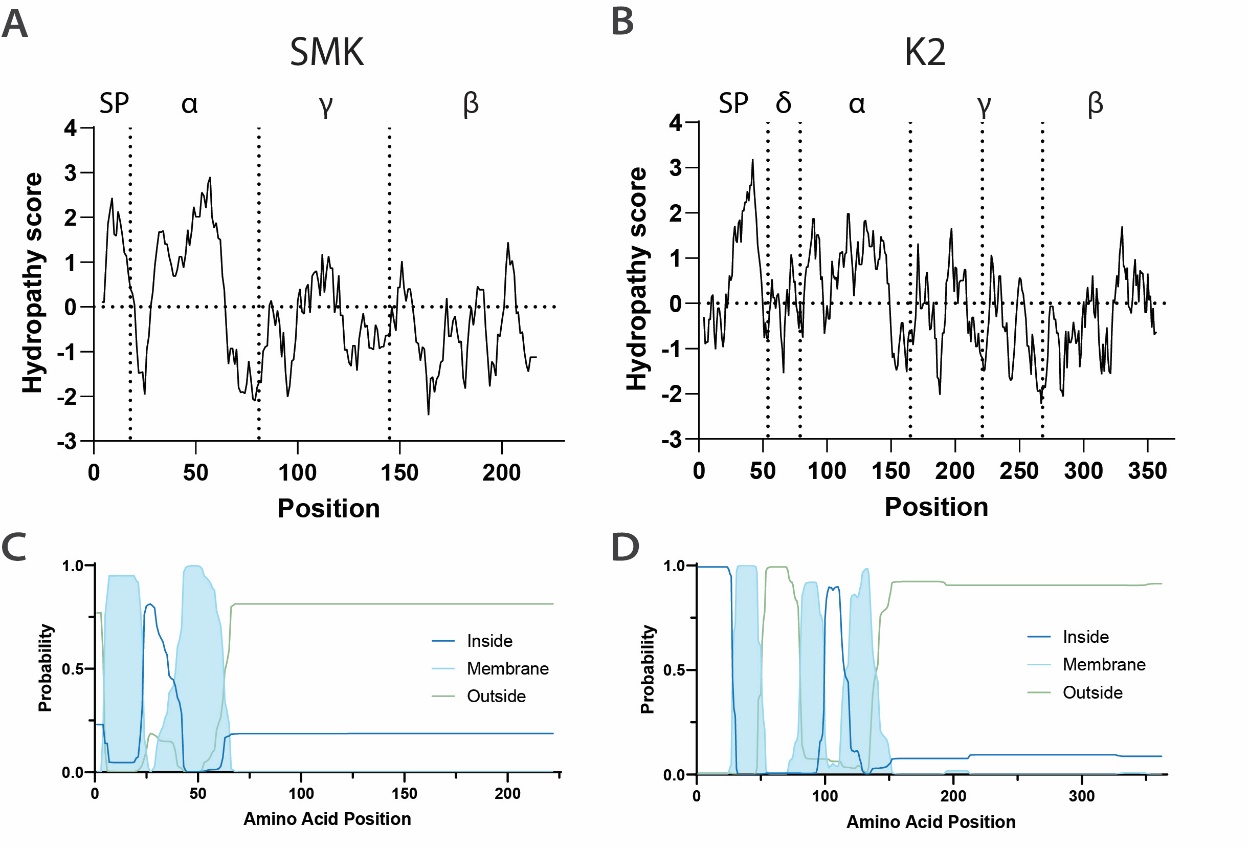

**Supplementary Figure S6. K2 and SMK hydropathy profiles and TM prediction.** Hydropathy profiles for the (**A**) SMK and (**B**) K2 precursor sequences. The proteolytic processing sites are indicated by vertical dotted lines. TMHMM-2.0 (Krogh et al., 2001) transmembrane helix prediction for the (**C**) SMK and (**D**) K2 precursors.

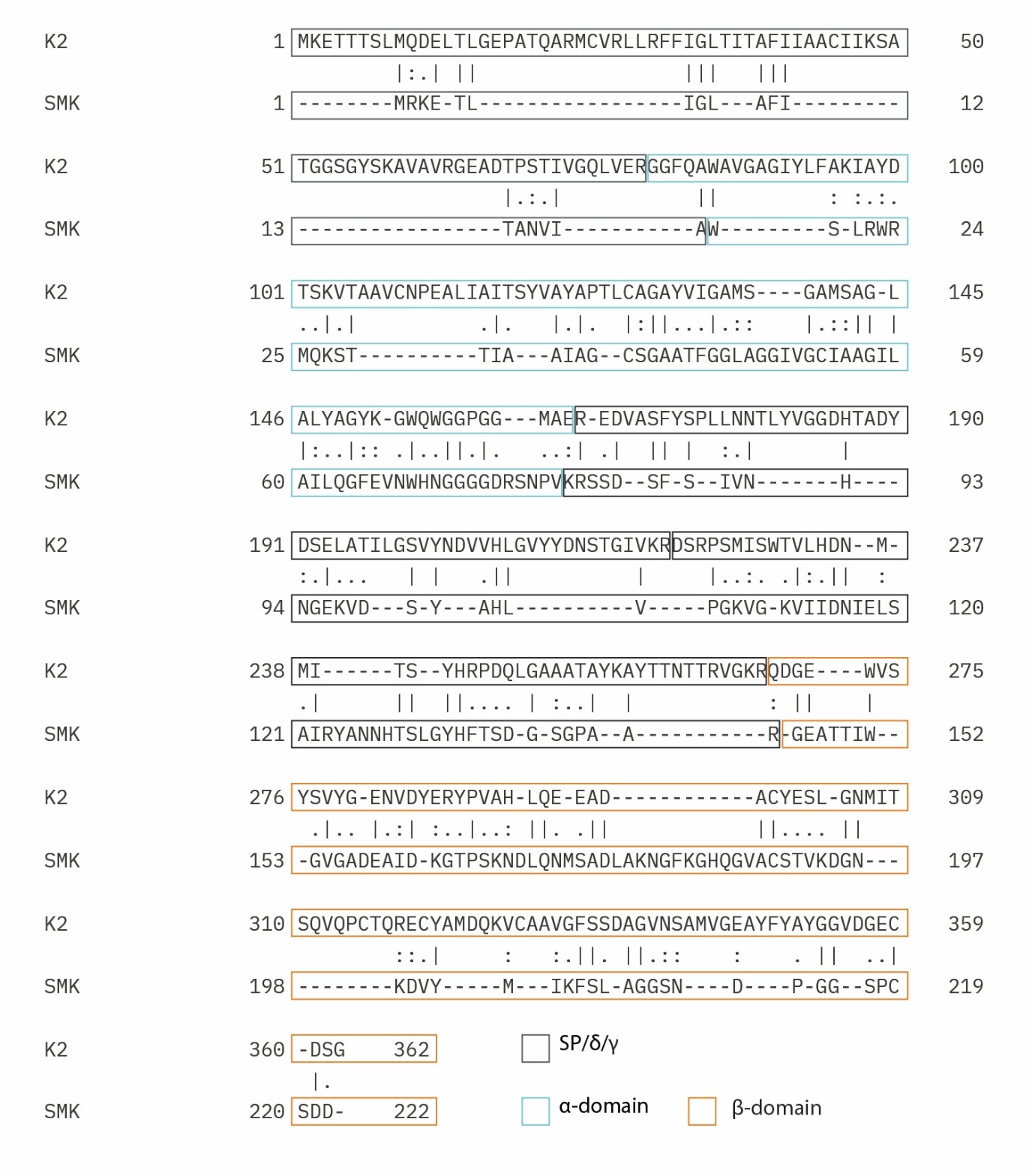

**Supplementary Figure S7. K2 and SMK pairwise amino acid sequence alignment.** A pairwise sequence alignment was created using the EBLOSUM62 matrix (gap_penalty of 5.0 and extend_penalty of 0.1). This resulted in approximately 20% sequence identity and 28% sequence similarity between K2 and SMK sequences. The domains are highlighted with boxes: SP; signal peptide/preregion, δ; proregion.

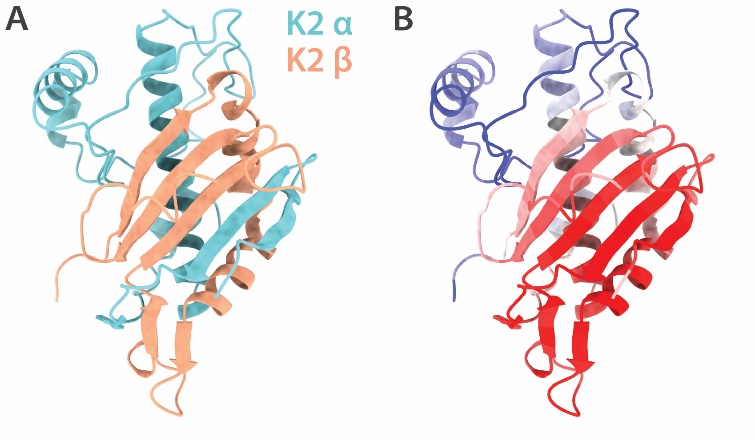

**Supplementary Figure S8. Structure prediction of the mature K2 toxin containing an α-subunit consisting of residues 80-219.** (**A**) Predicted structure of the K2 toxin, using α-subunit residues 80-219 and β-subunit residues 269-362. Average pLDDT score: 71.7. (**B**) pLDDT scores associated with the structure are displayed, scale: blue(low)-white-red(high), indicating the region within the α-subunit with lower pLDDT scores.

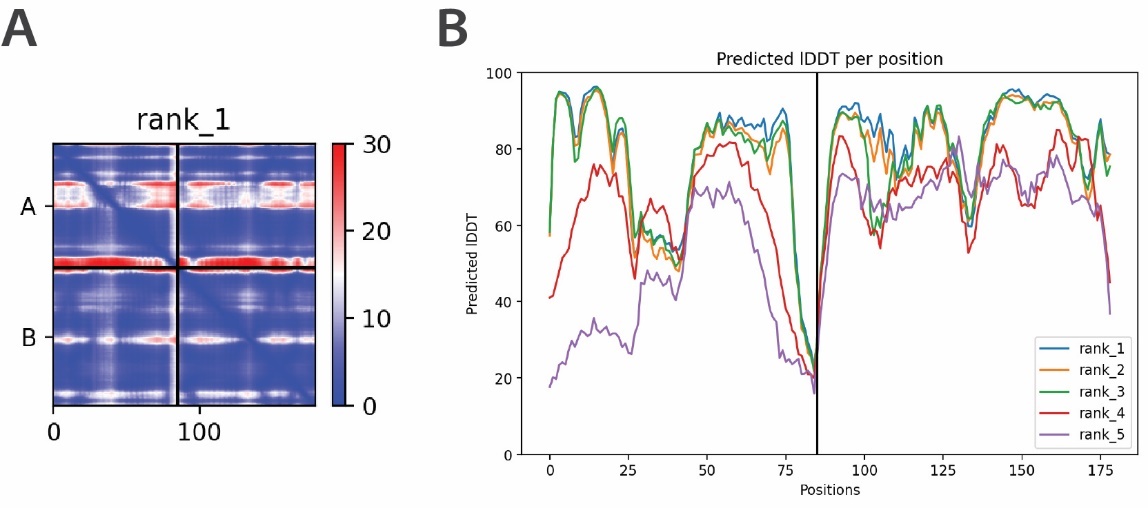

**Supplementary Figure S9. Predicted alignment error (PAE) and predicted local distance difference test (pLDDT) scores associated with the mature K2 toxin structure prediction. (A)** PAE plot of the predicted structure of the K2 mature toxin (A; α-subunit chain, B; β-subunit chain). (**B**) pLDDT scores per position of the K2 mature toxin structure (rank 1, blue line) (average pLDDT 79.8).

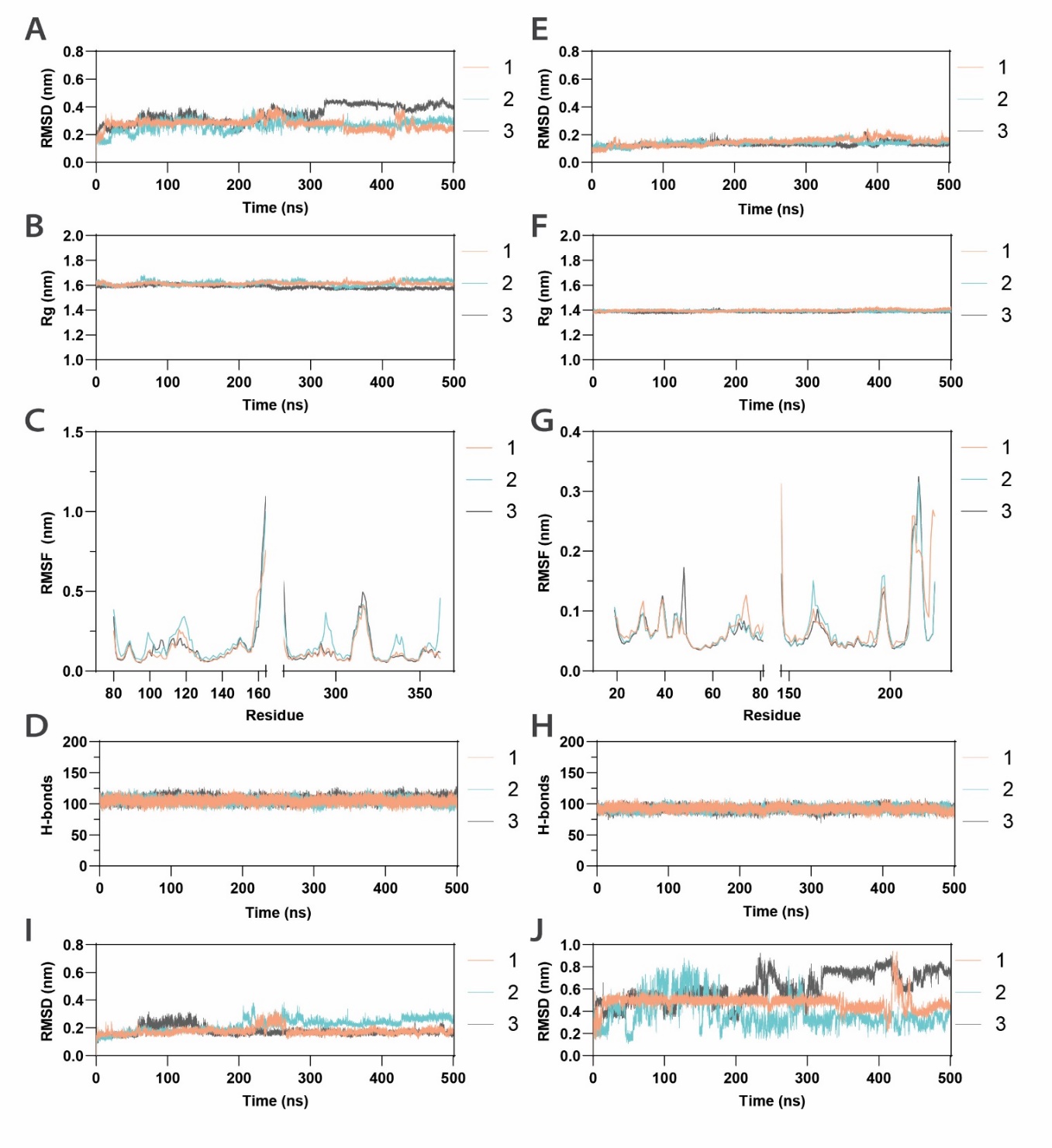

**Supplementary Figure S10. Molecular dynamics simulations of K2 and SMK mature toxins.** Three independent 500 ns simulations were generated of the K2 and SMK mature toxins in a predicted protonation state resembling pH 4.0. Several analyses of the trajectories are displayed in the different graphs. (**A**) K2, C_α_ root mean square deviation (RMSD). (**B**) K2, protein radius of gyration (Rg). (**C**) K2, C_α_ root mean square fluctuation (RMSF) per residue. (**D**) K2 hydrogen bonds within the protein over all timeframes (the average was 106 bonds). (**E**) SMK, C_α_ RMSD. (**F**) SMK, protein Rg. (**G**) SMK, C_α_ RMSF per residue. (**H**) SMK hydrogen bond counts within the protein over time (the average was 92 bonds). (**I**) K2, C_α_ RMSD of all residues except for residues 151-164. (**J**) K2, C_α_ RMSD of residues 151-164 only.
The RMSD of SMK is highly stable over time and only shows minimal deviation from the crystal structure. The RMSD of K2 shows some fluctuation, but within a stable range. The RMSF plot indicates which regions in the protein structure are more dynamic. The stable Rg and hydrogen bond values indicate that the proteins remain overall stable in a compact (folded) form throughout the simulation process. Plot (**C**), (**I**) and (**J**) together show that much of the C_α_ RMSD fluctuation for K2 is due to the C-terminal region of the α-subunit, consisting of residues ~151-164.

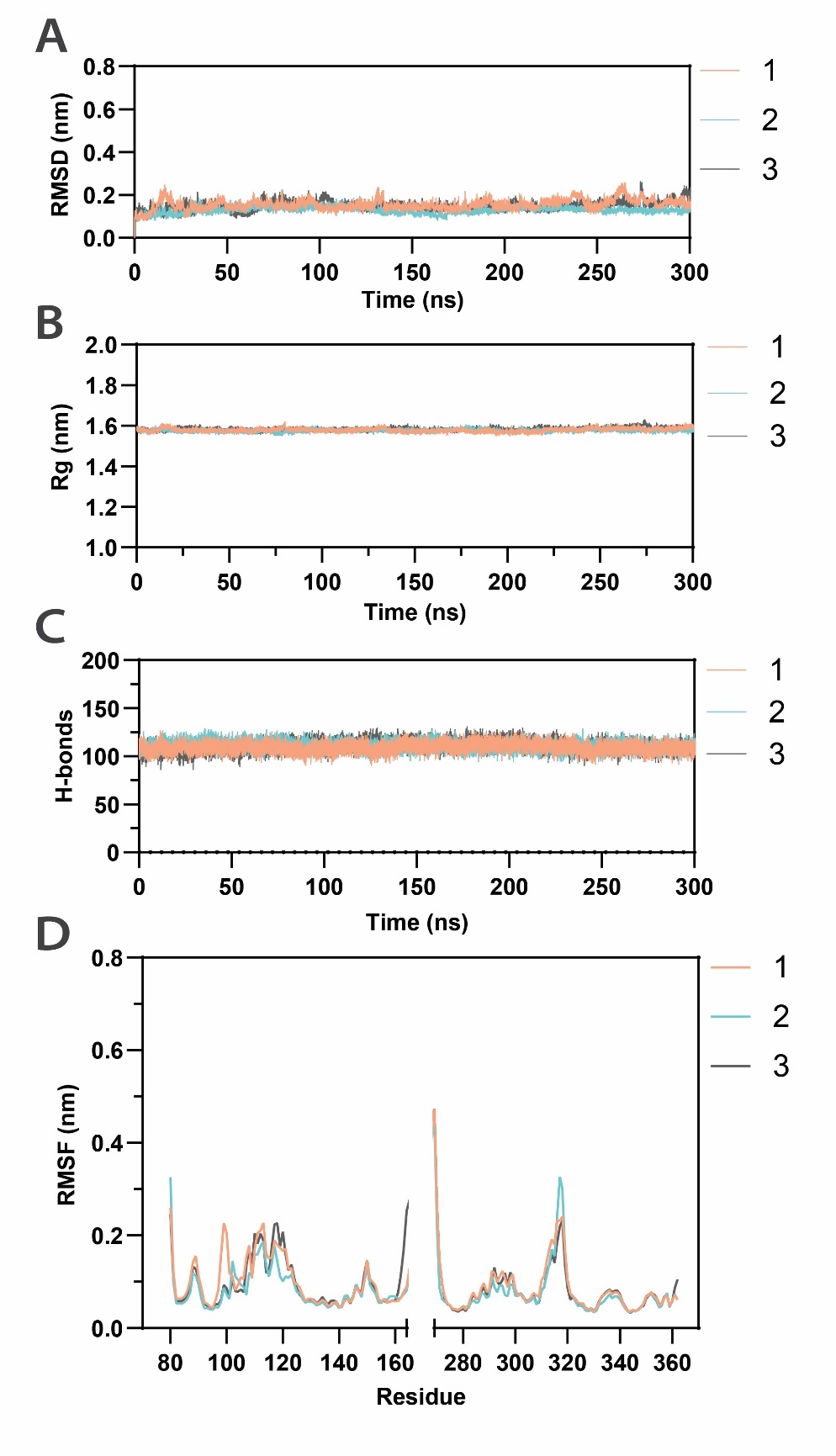

**Supplementary Figure S11. Molecular dynamics simulations of K2 mature toxin with altered protonation state.** Three independent 300 ns simulations were generated, continuing from the previous replicate 3 largest cluster centroid structure with an altered protonation state of D270 and the C-terminus of E164. (**A**) C_α_ root mean square deviation (RMSD). (**B**) Protein radius of gyration (Rg). (**C**) Hydrogen bond number within the protein over time. (**D**) C_α_ root mean square fluctuation (RMSF) per residue.

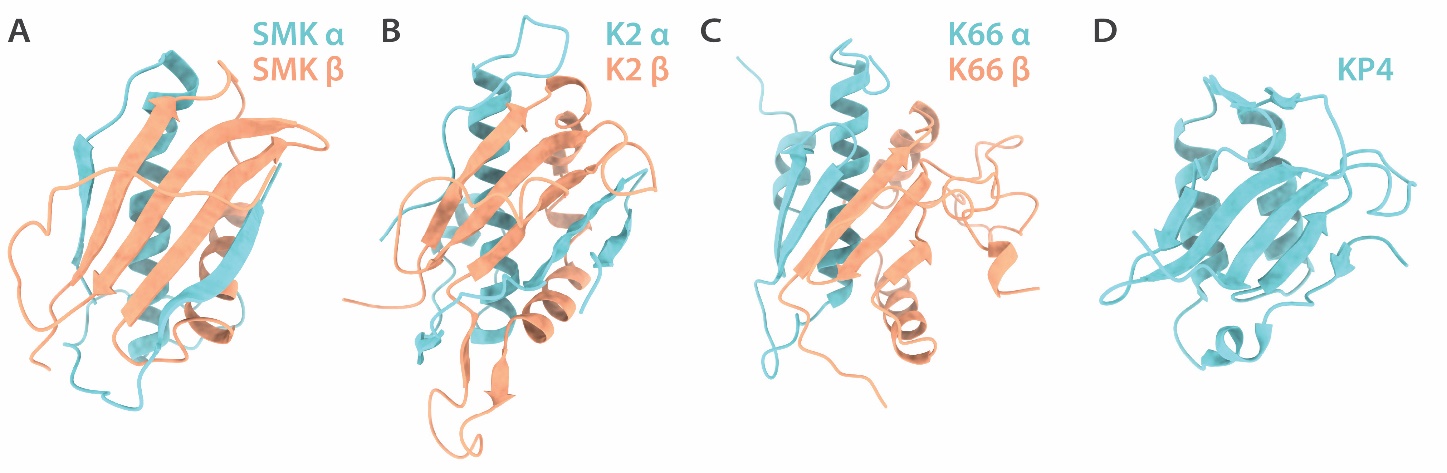

**Supplementary Figure S12. Comparison of mature toxin structures of SMK, K2, K66 and KP4.** (**A**) SMK: PDB ID 1KVE. (**B**) Mature K2 toxin structure resulting from the MD simulations. (**C**) Mature K66 toxin (residues α 60-128 and β 240-346 were used, average pLDDT 74.2) predicted using ColabFold. (**D**) KP4: PDB ID 1KPT.

**Supplementary Tables**

**Supplementary Table S1.** The K2 ORF was divided into 16 segments (Prins & Billerbeck, 2024). The start and end codons of each segment are indicated, as well as the amino acid (aa) length of each segment. The stop codon (363) was excluded from mutagenesis.

| **Segment** | **Start** | **End** | **Length (aa)** |
| --- | --- | --- | --- |
| **1** | 1 | 27 | 27 |
| **2** | 28 | 54 | 27 |
| **3** | 55 | 81 | 27 |
| **4** | 82 | 108 | 27 |
| **5** | 109 | 135 | 27 |
| **6** | 136 | 162 | 27 |
| **7** | 163 | 189 | 27 |
| **8** | 190 | 216 | 27 |
| **9** | 217 | 243 | 27 |
| **10** | 244 | 268 | 25 |
| **11** | 269 | 284 | 16 |
| **12** | 285 | 300 | 16 |
| **13** | 301 | 316 | 16 |
| **14** | 317 | 332 | 16 |
| **15** | 333 | 348 | 16 |
| **16** | 349 | 362 | 14 |

**Supplementary Table S2.** Number of white colonies obtained after library transformation into *E. coli*. The theoretical completeness of the library was calculated.

| **Sublibrary segment** | **White colonies (approx.)** | **Theoretical Completeness** |
| --- | --- | --- |
| **Alanine scanning** | | |
| 1 | 2532 | 1 |
| 2 | 2088 | 1 |
| 3 | 306 | 1 |
| 4 | 1260 | 1 |
| 5 | 800 | 1 |
| 6 | 1332 | 1 |
| 7 | 2400 | 1 |
| 8 | 3480 | 1 |
| 9 | 504 | 1 |
| 10 | 696 | 1 |
| 11 | 1776 | 1 |
| 12 | 510 | 1 |
| 13 | 2460 | 1 |
| 14 | 1296 | 1 |
| 15 | 960 | 1 |
| 16 | 1080 | 1 |
| **Saturation mutagenesis** | | |
| 293 | 800 | 1 |
| 357 | 800 | 1 |

**Supplementary Table S3.** Probability of library completeness in *S. cerevisiae.* In total 1228 colonies were screened (oversampling factor 3.4).

| **Sublibrary segment** | **Screened colonies** | **Expected number of variants** | **Probability** |
| --- | --- | --- | --- |
| **1** | 88 | 27 | 0.96 |
| **2** | 88 | 27 | 0.96 |
| **3** | 86 | 27 | 0.96 |
| **4** | 88 | 27 | 0.96 |
| **5** | 88 | 27 | 0.96 |
| **6** | 88 | 27 | 0.96 |
| **7** | 86 | 27 | 0.96 |
| **8** | 88 | 27 | 0.96 |
| **9** | 88 | 27 | 0.96 |
| **10** | 88 | 25 | 0.97 |
| **11** | 56 | 16 | 0.97 |
| **12** | 64 | 16 | 0.98 |
| **13** | 56 | 16 | 0.97 |
| **14** | 56 | 16 | 0.97 |
| **15** | 64 | 16 | 0.98 |
| **16** | 56 | 14 | 0.98 |
| ***Sum*** | *1228* | *362* |  |

**Supplementary Table S4.** Number of colonies found in each bin per K2 sublibrary segment in *S. cerevisiae*.

| **Sublibrary** | **Bin 1** | **Bin 2** | **Bin 3** | **Bin 4** | ***Sum*** |
| --- | --- | --- | --- | --- | --- |
| **1** | 1 | 1 | 2 | 84 | *88* |
| **2** | 0 | 0 | 0 | 88 | *88* |
| **3** | 6 | 0 | 16 | 64 | *86* |
| **4** | 14 | 9 | 38 | 27 | *88* |
| **5** | 28 | 17 | 34 | 9 | *88* |
| **6** | 20 | 14 | 41 | 13 | *88* |
| **7** | 12 | 13 | 21 | 40 | *86* |
| **8** | 7 | 6 | 25 | 50 | *88* |
| **9** | 9 | 7 | 21 | 51 | *88* |
| **10** | 4 | 7 | 8 | 69 | *88* |
| **11** | 12 | 5 | 25 | 14 | *56* |
| **12** | 7 | 9 | 14 | 34 | *64* |
| **13** | 13 | 5 | 17 | 21 | *56* |
| **14** | 5 | 7 | 24 | 20 | *56* |
| **15** | 11 | 4 | 26 | 23 | *64* |
| **16** | 13 | 15 | 8 | 20 | *56* |
| ***Sum*** | *162* | *119* | *320* | *627* | *1228* |

**Supplementary Table S5.** Barcoded primer combinations used for creation of amplicons for NGS of the alanine scanning mutagenesis. The K2 sequence was divided into 5 sequencing segments (SS).

| **SS** | **Bin 1 (<10%)** | **Bin 2 (10-40%)** | **Bin 3 (40-80%)** | **Bin 4 (80<)** | **Bin rv** |
| --- | --- | --- | --- | --- | --- |
| 1 | RP48-1 | RP48-2 | RP48-3 | RP48-4 | RP49-5 |
| 2 | RP50-1 | RP50-2 | RP50-3 | RP50-4 | RP51-6 |
| 3 | RP52-1 | RP52-2 | RP52-3 | RP52-4 | RP53-7 |
| 4 | RP54-1 | RP54-2 | RP54-3 | RP54-4 | RP55-8 |
| 5 | RP56-1 | RP56-2 | RP56-3 | RP56-4 | RP57-9 |

**Supplementary Table S6.** Distribution of functional scores of variants (between 1 and 4) per amino acid (AA). Total in K2; total number of residues of the respective amino acid present in the K2 amino acid sequence. Missing; number of the amino acid residues that were not present in this library.

| **AA** | **1** | **1,5** | **2** | **2,5** | **3** | **3,5** | **4** | ***SUM*** | ***Total in K2*** | ***Missing*** |
| --- | --- | --- | --- | --- | --- | --- | --- | --- | --- | --- |
| A | 1 | 1 | 2 | 2 | 14 | 4 | 19 | *43* | *46* | *3* |
| C | 6 | 0 | 0 | 0 | 0 | 0 | 3 | *9* | *9* | *0* |
| D | 0 | 0 | 1 | 1 | 1 | 3 | 13 | *19* | *19* | *0* |
| E | 0 | 0 | 0 | 1 | 2 | 3 | 11 | *17* | *18* | *1* |
| F | 1 | 0 | 3 | 0 | 1 | 0 | 3 | *8* | *8* | *0* |
| G | 2 | 0 | 5 | 4 | 6 | 6 | 13 | *36* | *38* | *2* |
| H | 0 | 0 | 0 | 1 | 1 | 0 | 3 | *5* | *5* | *0* |
| I | 2 | 0 | 1 | 0 | 1 | 1 | 9 | *14* | *17* | *3* |
| K | 0 | 0 | 1 | 0 | 2 | 2 | 5 | *10* | *10* | *0* |
| L | 2 | 1 | 2 | 2 | 4 | 2 | 8 | *21* | *22* | *1* |
| M | 0 | 1 | 0 | 1 | 4 | 1 | 4 | *11* | *12* | *1* |
| N | 0 | 0 | 0 | 1 | 2 | 2 | 5 | *10* | *10* | *0* |
| P | 0 | 0 | 1 | 0 | 2 | 1 | 6 | *10* | *10* | *0* |
| Q | 0 | 0 | 0 | 0 | 1 | 2 | 7 | *10* | *12* | *2* |
| R | 1 | 0 | 0 | 1 | 0 | 1 | 8 | *11* | *13* | *2* |
| S | 0 | 0 | 0 | 0 | 4 | 2 | 18 | *24* | *26* | *2* |
| T | 0 | 0 | 0 | 0 | 3 | 2 | 22 | *27* | *27* | *0* |
| V | 1 | 0 | 1 | 1 | 12 | 5 | 8 | *28* | *29* | *1* |
| W | 4 | 0 | 0 | 0 | 1 | 0 | 0 | *5* | *5* | *0* |
| Y | 6 | 0 | 4 | 1 | 3 | 3 | 5 | *22* | *26* | *4* |
| ***SUM*** | *26* | *3* | *21* | *16* | *64* | *40* | *170* | *340* | *362* | *22* |

**Supplementary Table S7.** Number of individual variants assigned to each functional score per K2 sublibrary segment in *S. cerevisiae*. The relative percentage of variants within the sublibrary (calculated across each individual sublibrary segment row) is given in the column to the right of each functional bin.

| **Sublibrary** | **1** | **%** | **1.5** | **%** | **2** | **%** | **2.5** | **%** | **3** | **%** | **3.5** | **%** | **4** | ***%*** | ***Sum*** |
| --- | --- | --- | --- | --- | --- | --- | --- | --- | --- | --- | --- | --- | --- | --- | --- |
| **1** | 0 | 0 | 0 | 0 | 0 | 0 | 0 | 0 | 1 | 4 | 0 | 0 | 25 | 96 | *26* |
| **2** | 0 | 0 | 0 | 0 | 0 | 0 | 0 | 0 | 0 | 0 | 0 | 0 | 25 | 100 | *25* |
| **3** | 0 | 0 | 0 | 0 | 0 | 0 | 0 | 0 | 3 | 12 | 2 | 8 | 20 | 80 | *25* |
| **4** | 3 | 13 | 0 | 0 | 2 | 9 | 2 | 9 | 8 | 35 | 2 | 9 | 6 | 26 | *23* |
| **5** | 7 | 27 | 1 | 4 | 5 | 19 | 0 | 0 | 10 | 38 | 0 | 0 | 3 | 12 | *26* |
| **6** | 3 | 11 | 2 | 7 | 3 | 11 | 2 | 7 | 12 | 44 | 1 | 4 | 4 | 15 | *27* |
| **7** | 1 | 4 | 0 | 0 | 4 | 15 | 0 | 0 | 3 | 12 | 4 | 15 | 14 | 54 | *26* |
| **8** | 1 | 4 | 0 | 0 | 1 | 4 | 2 | 7 | 2 | 7 | 8 | 30 | 13 | 48 | *27* |
| **9** | 2 | 8 | 0 | 0 | 1 | 4 | 1 | 4 | 3 | 12 | 5 | 19 | 14 | 54 | *26* |
| **10** | 0 | 0 | 0 | 0 | 0 | 0 | 2 | 8 | 1 | 4 | 1 | 4 | 20 | 83 | *24* |
| **11** | 1 | 8 | 0 | 0 | 0 | 0 | 2 | 15 | 5 | 38 | 2 | 15 | 3 | 23 | *13* |
| **12** | 0 | 0 | 0 | 0 | 2 | 13 | 0 | 0 | 3 | 19 | 4 | 25 | 7 | 44 | *16* |
| **13** | 3 | 21 | 0 | 0 | 0 | 0 | 1 | 7 | 2 | 14 | 5 | 36 | 3 | 21 | *14* |
| **14** | 2 | 13 | 0 | 0 | 0 | 0 | 3 | 20 | 3 | 20 | 3 | 20 | 4 | 27 | *15* |
| **15** | 1 | 7 | 0 | 0 | 0 | 0 | 1 | 7 | 5 | 36 | 3 | 21 | 4 | 29 | *14* |
| **16** | 2 | 15 | 0 | 0 | 3 | 23 | 0 | 0 | 3 | 23 | 0 | 0 | 5 | 38 | *13* |
| ***Sum*** | *26* |  | *3* |  | *21* |  | *16* |  | *64* |  | *40* |  | *170* |  | *340* |
| ***%*** | *7,65* |  | *0,88* |  | *6,18* |  | *4,71* |  | *18,82* |  | *11,76* |  | *50,00* |  |  |

**Supplementary Table S8.** Barcoded primer combinations used for creation of amplicons for NGS of the saturation mutagenesis of positions 293 and 357.

| **Position** | **Bin 1 (<80%)** | **Bin 2 (80-120%)** | **Bin 3 (120-160%)** | **Bin 4 (160+)** | **Bin rv** |
| --- | --- | --- | --- | --- | --- |
| 293 | RP54-1 | RP54-2 | RP54-3 | RP54-4 | RP55-8 |
| 357 | RP56-1 | RP56-2 | RP56-3 | RP56-4 | RP57-9 |

**Supplementary Table S9.** Plasmids used in this study.

| **Plasmid** | **Name** | **Notes** | **Benchling/Addgene link** | **Reference or source** |
| --- | --- | --- | --- | --- |
| **General plasmids** | | | | |
| pRP002 | pGAL1_K2_tADH1 (WT K2) | pRS423-type Wild-type K2 | https://benchling.com/s/seq-E1w0Ohr4cgXF8OCxrM8J?m=slm-w3UyjPbUjjo4FafJx4IE | (Prins & Billerbeck, 2024) |
| pRP003 | Empty plasmid (EP) | pRS413-type | https://benchling.com/s/seq-ISND4XmJDVAAmC1FDCb8?m=slm-gXRvrrT3nOpveBAZ1BAO |  |
| pYTK047 | Superfolder GFP | GFP-dropout module source | https://www.addgene.org/65154/ | (Lee et al., 2015) |
| pPTK007 | αMFΔ | Signal peptide of the α mating factor | https://www.addgene.org/84966/ | (Obst et al., 2017) |
| pRP091 | pYTK001-K2_1-54_ | K2 N-terminal peptide residues 1-54 type 3a | https://benchling.com/s/seq-e12yFPy0GjYzQjpUQqAW?m=slm-JbEMaKOn5P61BMEmQrnR | (Prins & Billerbeck, 2024) |
| pRP246 | Empty plasmid (EP) | Empty plasmid MoClo-type with spacer | https://benchling.com/s/seq-YVDbOQ9EsdtGjsCRnsVw?m=slm-TN18jAHVbFlcPStNPcbt | (Prins & Billerbeck, 2024) |
| pRP223 | pGAL1_K2_tENO1 (WT K2) | K2 wild-type MoClo-type | https://benchling.com/s/seq-Q1DQYXKBI0DzQ1IMfeky?m=slm-6iJC4fykPLVHl1eiP3hP | (Prins & Billerbeck, 2024) |
| **SbfI-site segment replaced constructs** | | | | |
| pRP035 | Segment 1 SbfI | For Gibson Assembly of GFP-dropout construct | https://benchling.com/s/seq-8m2Ga1TjqeIleGi5k9Hg?m=slm-KeE19MgFNVug4WTt8HDR | (Prins & Billerbeck, 2024) |
| pRP036 | Segment 2 SbfI | For Gibson Assembly of GFP-dropout construct | https://benchling.com/s/seq-WGMrfroM2MamJhfVYSJA?m=slm-SKAJTSn9FG8kesV0yhkI | (Prins & Billerbeck, 2024) |
| pRP037 | Segment 3 SbfI | For Gibson Assembly of GFP-dropout construct | https://benchling.com/s/seq-JNpLGpHYRsAQFuPuKV1x?m=slm-vziyRKFDi0WVtva0AD4Y | (Prins & Billerbeck, 2024) |
| pRP038 | Segment 4 SbfI | For Gibson Assembly of GFP-dropout construct | https://benchling.com/s/seq-CBgJqPIh8x0So8dAyuFc?m=slm-PJyfFdFpNxav2G3sKQRB | (Prins & Billerbeck, 2024) |
| pRP039 | Segment 5 SbfI | For Gibson Assembly of GFP-dropout construct | https://benchling.com/s/seq-DjzlnR8fTVUUWjA4UG5E?m=slm-pqK77LFHQ8s8G1lO6lMd | (Prins & Billerbeck, 2024) |
| pRP040 | Segment 6 SbfI | For Gibson Assembly of GFP-dropout construct | https://benchling.com/s/seq-7CK2HsFX2ASKYLiuplla?m=slm-S2Cn5TIq6CTHw81CUauJ | (Prins & Billerbeck, 2024) |
| pRP041 | Segment 7 SbfI | For Gibson Assembly of GFP-dropout construct | https://benchling.com/s/seq-q7TuN62wSyqcFGY5qTpd?m=slm-FVeoz0NDSpAD2bAoO417 | (Prins & Billerbeck, 2024) |
| pRP042 | Segment 8 SbfI | For Gibson Assembly of GFP-dropout construct | https://benchling.com/s/seq-LPbRKTTrCQGW43QzFqKs?m=slm-wtAxiB5FFqpxlee3y7tz | (Prins & Billerbeck, 2024) |
| pRP043 | Segment 9 SbfI | For Gibson Assembly of GFP-dropout construct | https://benchling.com/s/seq-hrngD3Sm9EQ3Ow280x0X?m=slm-lrS2eDd7Ws1ht67FH18q | (Prins & Billerbeck, 2024) |
| pRP044 | Segment 10 SbfI | For Gibson Assembly of GFP-dropout construct | https://benchling.com/s/seq-AMIhFm24lpNyzqpDnDlb?m=slm-iNEElWPONyKflIqHDcuv | (Prins & Billerbeck, 2024) |
| pRP045 | Segment 11 SbfI | For Gibson Assembly of GFP-dropout construct | https://benchling.com/s/seq-Xhta7Usp79RbkbxnyRXF?m=slm-I8T6DVkUc3oM5T9FYCeG | (Prins & Billerbeck, 2024) |
| pRP046 | Segment 12 SbfI | For Gibson Assembly of GFP-dropout construct | https://benchling.com/s/seq-6Il4rddwoj7DwW0xAhsa?m=slm-QUX5uy0OcbknqmLqNF1l | (Prins & Billerbeck, 2024) |
| pRP047 | Segment 13 SbfI | For Gibson Assembly of GFP-dropout construct | https://benchling.com/s/seq-PJ80MxUSikW41K8gguQH?m=slm-WapChdqoAOFgIfA64PtJ | (Prins & Billerbeck, 2024) |
| pRP048 | Segment 14 SbfI | For Gibson Assembly of GFP-dropout construct | https://benchling.com/s/seq-ryEqugH7eHI3wLCVzGJF?m=slm-K9zUGvLX0PpMo7qq0YdO | (Prins & Billerbeck, 2024) |
| pRP049 | Segment 15 SbfI | For Gibson Assembly of GFP-dropout construct | https://benchling.com/s/seq-BKmeJUOXJyZ9694e4tbn?m=slm-AfjRkeHZwjsOHHwWhGff | (Prins & Billerbeck, 2024) |
| pRP050 | Segment 16 SbfI | For Gibson Assembly of GFP-dropout construct | https://benchling.com/s/seq-xKiGMDJpLJETvGnpKvzm?m=slm-w1zaKdvuOZuxRltMDrL0 | (Prins & Billerbeck, 2024) |
| **GFP-dropout destination vectors** | | | | |
| pRP139 | Segment 1 GFP-dropout | GFP-dropout for library assembly with oligo pool | https://benchling.com/s/seq-GkFqGKUfA4Wev89E8hhD?m=slm-jwsuhchy44z18etDxuXk | This study |
| pRP140 | Segment 2 GFP-dropout | GFP-dropout for library assembly with oligo pool | https://benchling.com/s/seq-tlFIjPdEB50Q6MYzYPy4?m=slm-PHaQ2bKwhwQZt5feaenK | This study |
| pRP141 | Segment 3 GFP-dropout | GFP-dropout for library assembly with oligo pool | https://benchling.com/s/seq-vkDbK0EiqJiJWHzow2Vn?m=slm-gAjFqOHKYFM8f6gYKsMS | This study |
| pRP142 | Segment 4 GFP-dropout | GFP-dropout for library assembly with oligo pool | https://benchling.com/s/seq-2DMsKAbxbnQHoGTo51ES?m=slm-xFeTzzk9Xo0B1djUIbWV | This study |
| pRP143 | Segment 5 GFP-dropout | GFP-dropout for library assembly with oligo pool | https://benchling.com/s/seq-XTlYesqsqyqhrnL9LK4N?m=slm-Sfg2dj0TcNNtZbzoJu0j | This study |
| pRP144 | Segment 6 GFP-dropout | GFP-dropout for library assembly with oligo pool | https://benchling.com/s/seq-rqTAgyK8gPmZxJlDzZh2?m=slm-wlX8ezwC1bjDhNpHaIXn | This study |
| pRP145 | Segment 7 GFP-dropout | GFP-dropout for library assembly with oligo pool | https://benchling.com/s/seq-1XeB7UzzRHS8nh6Uyjvm?m=slm-I4yxAt1zgP1VCbueieIk | This study |
| pRP146 | Segment 8 GFP-dropout | GFP-dropout for library assembly with oligo pool | https://benchling.com/s/seq-6MjRBUgriEESg3zcwCQV?m=slm-QTNYIMBVytd6iLqlVVgy | This study |
| pRP147 | Segment 9 GFP-dropout | GFP-dropout for library assembly with oligo pool | https://benchling.com/s/seq-ymq9oGkFEdxA2NNsGxxa?m=slm-vkgwKWzGtBHOMhvPef4V | This study |
| pRP148 | Segment 10 GFP-dropout | GFP-dropout for library assembly with oligo pool | https://benchling.com/s/seq-AguuBgyfNZaZOtIaoQ4H?m=slm-yLhe7dfNaPkHMxEvrp6I | This study |
| pRP149 | Segment 11 GFP-dropout | GFP-dropout for library assembly with oligo pool | https://benchling.com/s/seq-3yu8mlCZL9QkzIbpcw9B?m=slm-JLUF5xLJb2h4lEShMspt | This study |
| pRP150 | Segment 12 GFP-dropout | GFP-dropout for library assembly with oligo pool | https://benchling.com/s/seq-i8rErVyRlGFAvEvyrPhZ?m=slm-nhVFqIcmEWnhZ1vxdPMd | This study |
| pRP151 | Segment 13 GFP-dropout | GFP-dropout for library assembly with oligo pool | https://benchling.com/s/seq-KN1G424MCytiEyqJgevQ?m=slm-GY8h2uqGXZxOZT6ieARt | This study |
| pRP152 | Segment 14 GFP-dropout | GFP-dropout for library assembly with oligo pool | https://benchling.com/s/seq-v8AyJv8YR8SGh4agGRRB?m=slm-5PphmDMKePujvYgq5vdC | This study |
| pRP153 | Segment 15 GFP-dropout | GFP-dropout for library assembly with oligo pool | https://benchling.com/s/seq-ebMHJnZxNChqH8G6vS0Q?m=slm-N7x1YF49ZqQ9WOt9iJPR | This study |
| pRP154 | Segment 16 GFP-dropout | GFP-dropout for library assembly with oligo pool | https://benchling.com/s/seq-gHlICSDfahNRivvVBc05?m=slm-NGuNjLnhtt4xV5LvXbh5 | This study |
| **K2 variant constructs** | | | | |
| pRP116 | L293A+G357A | K2 with double mutation | https://benchling.com/s/seq-JVtsuIot6yHtFyyMEUz3?m=slm-4pg9ofcD5ktzJ3lC6wpQ | This study |
| pRP420 | R79A | K2 with R79A mutation | https://benchling.com/s/seq-ZCMcbXcxLzcRSNQ4r34w?m=slm-701sapNlsnnzLp55GCRE | This study |
| pRP421 | E78K | K2 with E78K mutation | https://benchling.com/s/seq-2huOda0yaH4hhnw2tyC2?m=slm-ej0Ih0gNNRTq8tetblHs | This study |
| pRP422 | R165A | K2 with R165A mutation | https://benchling.com/s/seq-XeyXD4m9WrNE1Rf3s25H?m=slm-WusgEu0rpmM5XCPMjDWV | This study |
| pRP423 | E164K | K2 with E164K mutation | https://benchling.com/s/seq-VKwOYKEAzjc9dI14h6sc?m=slm-L1gdeBtnVRrJJApl0zVK | This study |
| pRP426 | ALFA-269 | ALFA-tag inserted at position 269 | https://benchling.com/s/seq-gaWNVyWHzNfOfL9ThLCj?m=slm-cDIigHANb1TVoqSfp7VV | This study |
| pRP427 | αMF_SP_:α_80-164_ | αMF signal peptide fused to K2 α-subunit 80-164 | https://benchling.com/s/seq-RwTa2ZHmawW0VRc8ogBG?m=slm-HUNM7LN45DqZZSyUDobi | This study |
| pRP358 | αMF_SP_-K2_54-219_ | αMF signal peptide fused to K2 54-219, previously shown to be suicidal | https://benchling.com/s/seq-tZvZy0bxIuXU2KQ0tjRN?m=slm-lOdfBWS2QNpPXjq4fL3o | (Prins & Billerbeck, 2024) |
| **Isolated K2 variants** | | | | |
| pRP120 | L293A | K2 L293A, isolated from the library | https://benchling.com/s/seq-6OLe0GhogMWwChx3xKA8?m=slm-qlv3OzGmca4pbO24txy4 | This study |
| pRP373 | L293R | K2 L293R, isolated from the library | https://benchling.com/s/seq-S1RmaBEDvE8wUrwdRii7?m=slm-rho2zDCrs8DgL8r9oFOq | This study |
| pRP374 | L293V | K2 L293V, isolated from the library | https://benchling.com/s/seq-vA0fPNFfOtaJnpAWJE3h?m=slm-PGuTDFHmk59YlGm869q4 | This study |
| pRP375 | L293I | K2 L293I, isolated from the library | https://benchling.com/s/seq-ESGT0gs5UvkVJBASvCuG?m=slm-NIvpN7fv5ctheJVN4tTU | This study |
| pRP376 | L293E | K2 L293E, isolated from the library | https://benchling.com/s/seq-LZ3eHaiRp8Xrxl7Gop8s?m=slm-chz6cb4jExA5evH0Ds6Y | This study |
| pRP377 | L293M | K2 L293M, isolated from the library | https://benchling.com/s/seq-rlVdDL0X1T6cI7UPCsde?m=slm-nKvEWbXT2VDh2jcpXOZg | This study |
| pRP121 | G357A | K2 G357A, isolated from the library | https://benchling.com/s/seq-hCOGNbEcpzNIxieUwBCB?m=slm-bGdjpI0xMbsY6hWPSOxB | This study |
| pRP383 | G357M | K2 G357M, isolated from the library | https://benchling.com/s/seq-74VwMlP2U8khQ5bgWtOw?m=slm-hB2ZnKoYiVslJcjUmKZK | This study |
| pRP384 | G357E | K2 G357E, isolated from the library | https://benchling.com/s/seq-6RdaIpjw3QOn3z5Yl6ek?m=slm-AzpP4eUJauxMcNsJ8nim | This study |
| pRP378 | G357K | K2 G357K, isolated from the library | https://benchling.com/s/seq-K8HfAB651v0rRM3PXEXv?m=slm-s1EAan6LVvskhu7n4s9Y | This study |
| pRP379 | G357Q | K2 G357Q, isolated from the library | https://benchling.com/s/seq-qJHXnI0zjmBQaQWWGwH2?m=slm-vxCCO0KoCR5MBW6OEbuX | This study |
| pRP380 | G357P | K2 G357P, isolated from the library | https://benchling.com/s/seq-2hzH8meD97ZKoLofib1h?m=slm-FKHLMLo3ENSzKycy0TT4 | This study |
| pRP381 | G357I | K2 G357I, isolated from the library | https://benchling.com/s/seq-OyujDCmrJSmYRrC4s891?m=slm-Kcp8MCzOyufNuLkVzR0V | This study |
| pRP382 | G357V | K2 G357V, isolated from the library | https://benchling.com/s/seq-vAQPxfUm4KLqyLfR6gQe?m=slm-Oz26pmbVjjbsvae9PpbD | This study |
| **N-terminal variants** | | | | |
| pRP395 | αMF_SP_-K2_65-219_ | αMF signal peptide fused to K2 65-219 | https://benchling.com/s/seq-OZwHu1eKoS62pRK6KOQA?m=slm-UH2Om9JwQFvXPdqlpJbg | This study |
| pRP396 | αMF_SP_-K2_75-219_ | αMF signal peptide fused to K2 75-219 | https://benchling.com/s/seq-YgTZqbZyd34FsMqhmxKP?m=slm-swRz4s4P5AY4mwzCzV4J | This study |
| pRP397 | αMF_SP_-K2_80-219_ | αMF signal peptide fused to K2 80-219 | https://benchling.com/s/seq-yj7RqVFYWMIAnn3k6BFd?m=slm-IBzsp3wZCRTWQfUPCa6W | This study |
| pRP398 | αMF_SP_-K2_90-219_ | αMF signal peptide fused to K2 90-219 | https://benchling.com/s/seq-fKVKnW7YLZWGwFUIVJjq?m=slm-aPzzGquJODwpxpf0PWX7 | This study |
| pRP399 | αMF_SP_-K2_100-219_ | αMF signal peptide fused to K2 100-219 | https://benchling.com/s/seq-r8iINYdxEdGNNJPrdf6z?m=slm-uiEWxuBmoV1cIFF2sL2X | This study |
| pRP400 | K2_SP_-K2_65-219_ | K2 signal peptide fused to K2 65-219 | https://benchling.com/s/seq-6NhEddbnYmm8fWlr1kr7?m=slm-cWFO7Wm5oJwVktWv25KH | This study |
| pRP401 | K2_SP_-K2_75-219_ | K2 signal peptide fused to K2 75-219 | https://benchling.com/s/seq-rHaKALKZFkUDi443WRAS?m=slm-T4dpYfu13MmUt2Z6Nuu2 | This study |
| pRP402 | K2_SP_-K2_80-219_ | K2 signal peptide fused to K2 80-219 | https://benchling.com/s/seq-i06DILPORmaxLLG52ogC?m=slm-pG2sn7pTutI503MaR8z5 | This study |
| pRP403 | K2_SP_-K2_90-219_ | K2 signal peptide fused to K2 90-219 | https://benchling.com/s/seq-8O9LXbqccAzaKfLxNq9W?m=slm-Gj1gLs0yX7wVjYNxcKmZ | This study |
| pRP404 | K2_SP_-K2_100-219_ | K2 signal peptide fused to K2 100-219 | https://benchling.com/s/seq-bWC1FS4BaJj02eSukwp8?m=slm-ZQrtW3STXhoYIYkmF07V | This study |
| pRP390 | K2_65-362_ | K2 residues 65-362, after initiator methionine | https://benchling.com/s/seq-VXZV6MezjB4xIxjx9usZ?m=slm-yqtEf1KQJDmaKog7Ojvh | This study |
| pRP391 | K2_75-362_ | K2 residues 75-362, after initiator methionine | https://benchling.com/s/seq-rKja2OK0iVK54OYOSGXg?m=slm-hCegb5O8xSB06Ve9klkA | This study |
| pRP392 | K2_80-362_ | K2 residues 80-362, after initiator methionine | https://benchling.com/s/seq-905Q70sax8WEe6fUu6WO?m=slm-7HNVaJQaoO0FPYk7fydt | This study |
| pRP393 | K2_90-362_ | K2 residues 90-362, after initiator methionine | https://benchling.com/s/seq-Iuw9SqSg1oXlVhObJIHt?m=slm-BupfhfabjIl3615xBBIt | This study |
| pRP394 | K2_100-362_ | K2 residues 100-362, after initiator methionine | https://benchling.com/s/seq-wVPG3rSmUIpoRKaj8fz5?m=slm-cjvphxvsUrLbFo36e6Y9 | This study |

**Supplementary Table S10.** Primers used in this study.

| **Primer** | **Note** | **Sequence** |
| --- | --- | --- |
| **Alanine scanning oligonucleotide pool primers** | | |
| RP139 | Segment 1 fw | CATCGGTCTCA AACC GAATTCCCAAAAGAA |
| RP140 | Segment 1 rv | CTCAGGTCTCA CAAT AAAAAAGCGCAGCAG |
| RP143 | Segment 2 fw | CATCGGTCTCA GCAT GTGCGTGCGC |
| RP144 | Segment 2 rv | CTCAGGTCTCA ACCG CTTTGCTATAGCC |
| RP147 | Segment 3 fw | CATCGGTCTCA ACCG GCGGCAGC |
| RP148 | Segment 3 rv | CTCAGGTCTCA GCCC ACGCCTGAAA |
| RP151 | Segment 4 fw | CATCGGTCTCA TGGA ACGCGGCGGC |
| RP152 | Segment 4 rv | CTCAGGTCTCA CGCT TCCGGGTTGCA |
| RP155 | Segment 5 fw | CATCGGTCTCA GACC GCGGCGGTG |
| RP156 | Segment 5 rv | CTCAGGTCTCA CGCC GCTCATCGC |
| RP159 | Segment 6 fw | CATCGGTCTCA CGCG TATGTGATTGGC |
| RP160 | Segment 6 rv | CTCAGGTCTCA CATC TTCGCGTTCCGC |
| RP163 | Segment 7 fw | CATCGGTCTCA CCCG GGCGGCATG |
| RP164 | Segment 7 rv | CTCAGGTCTCA CGCC AGTTCGCTATCATA |
| RP167 | Segment 8 fw | CATCGGTCTCA GCGA TCATACCGCGGAT |
| RP168 | Segment 8 rv | CTCAGGTCTCA CTAT CGCGTTTCACAATGCC |
| RP171 | Segment 9 fw | CATCGGTCTCA GTGT ATTATGATAACAGCACC |
| RP172 | Segment 9 rv | CTCAGGTCTCA AGCT GATCCGGGCG |
| RP175 | Segment 10 fw | CATCGGTCTCA CATG ATGATTACCAGCTATCAT |
| RP176 | Segment 10 rv | CTCAGGTCTCA CCCA TTCGCCATCCTG |
| RP179 | Segment 11 fw | CATCGGTCTCA GGGC AAACGC |
| RP180 | Segment 11 rv | CTCAGGTCTCA GGAT AGCGTTCATA |
| RP183 | Segment 12 fw | CATCGGTCTCA GCGA AAACGTGGAT |
| RP184 | Segment 12 rv | CTCAGGTCTCA CCAG GCTTTCATA |
| RP187 | Segment 13 fw | CATCGGTCTCA CGGA TGCGTGC |
| RP188 | Segment 13 rv | CTCAGGTCTCA ATTC GCGCTG |
| RP191 | Segment 14 fw | CATCGGTCTCA TGCA GCCGTGCACC |
| RP192 | Segment 14 rv | CTCAGGTCTCA CCGC ATCGCTGCTAAA |
| RP195 | Segment 15 fw | CATCGGTCTCA GCGG CGGTGGGC |
| RP196 | Segment 15 rv | CTCAGGTCTCA CGCC GCCATACGCATAAAA |
| RP199 | Segment 16 fw | CATCGGTCTCA GTGG GCGAAGCGTAT |
| RP200 | Segment 16 rv | CTCAGGTCTCA CAAA AGCAGAGACAGATATCTTA |
| **Saturation mutagenesis pool primers** | | |
| RP211 | Segment 12 rv | CTCAGGTCTCACCAG |
| RP212 | Segment 16 rv | CTCAGGTCTCACAAAAG |
| **GFP-dropout module primers** | | |
| RP137 | Segment 1 fw | TCTATACTTTAACGTCAAGGAGAAAA AACC TGAGACCGAAAGTGAAAC |
| RP138 | Segment 1 rv | CAATAATAAACGCGGTAATGGTCAGGCCAAT TGAGACCTATAAACGCAG |
| RP141 | Segment 2 fw | CCTGGGCGAACCGGCGACCCAGGCGC GCAT TGAGACCGAAAGTGAAAC |
| RP142 | Segment 2 rv | GGGTATCCGCTTCGCCGCGCACCGCC ACCG TGAGACCTATAAACGCAG |
| RP145 | Segment 3 fw | TTGCGGCGTGCATTATTAAAAGCGCG ACCG TGAGACCGAAAGTGAAAC |
| RP146 | Segment 3 rv | CAAACAGATAAATGCCCGCGCCCACC GCCC TGAGACCTATAAACGCAG |
| RP149 | Segment 4 fw | CCCGAGCACCATTGTGGGCCAGCTGG TGGA TGAGACCGAAAGTGAAAC |
| RP150 | Segment 4 rv | GCCACATAGCTGGTAATCGCAATCAG CGCT TGAGACCTATAAACGCAG |
| RP153 | Segment 5 fw | GAAAATTGCGTATGATACCAGCAAAGT GACC TGAGACCGAAAGTGAAAC |
| RP154 | Segment 5 rv | ATACAGCGCCAGGCCCGCGCTCATCG CGCC TGAGACCTATAAACGCAG |
| RP157 | Segment 6 fw | GTGGCGTATGCGCCGACCCTGTGCGCGGG CGCG TGAGACCGAAAGTGAAAC |
| RP158 | Segment 6 rv | CAGCAGCGGGCTATAAAAGCTCGCCA CATC TGAGACCTATAAACGCAG |
| RP161 | Segment 7 fw | GGCTATAAAGGCTGGCAGTGGGGCGG CCCG TGAGACCGAAAGTGAAAC |
| RP162 | Segment 7 rv | TCGTTATACACGCTGCCCAGAATGGT CGCC TGAGACCTATAAACGCAG |
| RP165 | Segment 8 fw | GCTGAACAACACCCTGTATGTGGGCG GCGA TGAGACCGAAAGTGAAAC |
| RP166 | Segment 8 rv | CACGGTCCAGCTAATCATGCTCGGGCGG CTAT TGAGACCTATAAACGCAG |
| RP169 | Segment 9 fw | GTGTATAACGATGTGGTGCATCTGGGC GTGT TGAGACCGAAAGTGAAAC |
| RP170 | Segment 9 rv | CTTTATACGCGGTCGCCGCCGCGCCC AGCT TGAGACCTATAAACGCAG |
| RP173 | Segment 10 fw | GATTAGCTGGACCGTGCTGCATGATAA CATG TGAGACCGAAAGTGAAAC |
| RP174 | Segment 10 rv | GTTTTCGCCATACACGCTATAGCTCA CCCA TGAGACCTATAAACGCAG |
| RP177 | Segment 11 fw | GCGTATACCACCAACACCACCCGCGT GGGC TGAGACCGAAAGTGAAAC |
| RP178 | Segment 11 rv | CATCCGCTTCTTCCTGCAGATGCGCCACC GGAT TGAGACCTATAAACGCAG |
| RP181 | Segment 12 fw | CGAATGGGTGAGCTATAGCGTGTATG GCGA TGAGACCGAAAGTGAAAC |
| RP182 | Segment 12 rv | CTGCACCTGGCTGGTAATCATGTTGC CCAG TGAGACCTATAAACGCAG |
| RP185 | Segment 13 fw | TCCGGTGGCGCATCTGCAGGAAGAAG CGGA TGAGACCGAAAGTGAAAC |
| RP186 | Segment 13 rv | CGCACACTTTCTGATCCATCGCATAGC ATTC TGAGACCTATAAACGCAG |
| RP189 | Segment 14 fw | AGCCTGGGCAACATGATTACCAGCCAGG TGCA TGAGACCGAAAGTGAAAC |
| RP190 | Segment 14 rv | CTTCGCCCACCATCGCGCTGTTCACGC CCGC TGAGACCTATAAACGCAG |
| RP193 | Segment 15 fw | GCTATGCGATGGATCAGAAAGTGTGC GCGG TGAGACCGAAAGTGAAAC |
| RP194 | Segment 15 rv | TAGCCGCTATCGCATTCGCCATCCAC GCC TGAGACCTATAAACGCAG |
| RP197 | Segment 16 fw | CAGCGATGCGGGCGTGAACAGCGCGATG GTGG TGAGACCGAAAGTGAAAC |
| RP198 | Segment 16 rv | AGAGGTACATACATAAACATACGCGCA CAAA TGAGACCTATAAACGCAG |
| **Next Generation Sequencing Primers** | | |
| RP48_1 | Amplicon 1 fw / barcode 1 | ACA CTC TTT CCC TAC ACG ACG CTC TTC CGA TCT AAACGATC CTATACTTTAACGTCAAGGAG |
| RP48_2 | Amplicon 1 fw / barcode 2 | ACA CTC TTT CCC TAC ACG ACG CTC TTC CGA TCT CCAGACAG CTATACTTTAACGTCAAGGAG |
| RP48_3 | Amplicon 1 fw / barcode 3 | ACA CTC TTT CCC TAC ACG ACG CTC TTC CGA TCT GGTAGCCA CTATACTTTAACGTCAAGGAG |
| RP48_4 | Amplicon 1 fw / barcode 4 | ACA CTC TTT CCC TAC ACG ACG CTC TTC CGA TCT TCTATTCC CTATACTTTAACGTCAAGGAG |
| RP49_5 | Amplicon 1 rv / barcode 5 | GAC TGG AGT TCA GAC GTG TGC TCT TCC GAT CT AAGGATGA CACTTTGCTGGTATCATACG |
| RP50_1 | Amplicon 2 fw / barcode 1 | ACA CTC TTT CCC TAC ACG ACG CTC TTC CGA TCT AAACGATC CGAGCACCATTGTGG |
| RP50_2 | Amplicon 2 fw / barcode 2 | ACA CTC TTT CCC TAC ACG ACG CTC TTC CGA TCT CCAGACAG CGAGCACCATTGTGG |
| RP50_3 | Amplicon 2 fw / barcode 3 | ACA CTC TTT CCC TAC ACG ACG CTC TTC CGA TCT GGTAGCCA CGAGCACCATTGTGG |
| RP50_4 | Amplicon 2 fw / barcode 4 | ACA CTC TTT CCC TAC ACG ACG CTC TTC CGA TCT TCTATTCC CGAGCACCATTGTGG |
| RP51_6 | Amplicon 2 rv / barcode 6 | GAC TGG AGT TCA GAC GTG TGC TCT TCC GAT CT CGACATTT CATAATCCGCGGTATGATC |
| RP52_1 | Amplicon 3 fw / barcode 1 | ACA CTC TTT CCC TAC ACG ACG CTC TTC CGA TCT AAACGATC GCTATAAAGGCTGGCAG |
| RP52_2 | Amplicon 3 fw / barcode 2 | ACA CTC TTT CCC TAC ACG ACG CTC TTC CGA TCT CCAGACAG GCTATAAAGGCTGGCAG |
| RP52_3 | Amplicon 3 fw / barcode 3 | ACA CTC TTT CCC TAC ACG ACG CTC TTC CGA TCT GGTAGCCA GCTATAAAGGCTGGCAG |
| RP52_4 | Amplicon 3 fw / barcode 4 | ACA CTC TTT CCC TAC ACG ACG CTC TTC CGA TCT TCTATTCC GCTATAAAGGCTGGCAG |
| RP53_7 | Amplicon 3 rv / barcode 7 | GAC TGG AGT TCA GAC GTG TGC TCT TCC GAT CT GTCTGGAA CATCCTGGCGTTTGC |
| RP54_1 | Amplicon 4 fw / barcode 1 | ACA CTC TTT CCC TAC ACG ACG CTC TTC CGA TCT AAACGATC CCGGCATTGTGAAACG |
| RP54_2 | Amplicon 4 fw / barcode 2 | ACA CTC TTT CCC TAC ACG ACG CTC TTC CGA TCT CCAGACAG CCGGCATTGTGAAACG |
| RP54_3 | Amplicon 4 fw / barcode 3 | ACA CTC TTT CCC TAC ACG ACG CTC TTC CGA TCT GGTAGCCA CCGGCATTGTGAAACG |
| RP54_4 | Amplicon 4 fw / barcode 4 | ACA CTC TTT CCC TAC ACG ACG CTC TTC CGA TCT TCTATTCC CCGGCATTGTGAAACG |
| RP55_8 | Amplicon 4 rv / barcode 8 | GAC TGG AGT TCA GAC GTG TGC TCT TCC GAT CT TTCTCGCG CACACTTTCTGATCCATC |
| RP56_1 | Amplicon 5 fw / barcode 1 | ACA CTC TTT CCC TAC ACG ACG CTC TTC CGA TCT AAACGATC GGTGAGCTATAGCGTG |
| RP56_2 | Amplicon 5 fw / barcode 2 | ACA CTC TTT CCC TAC ACG ACG CTC TTC CGA TCT CCAGACAG GGTGAGCTATAGCGTG |
| RP56_3 | Amplicon 5 fw / barcode 3 | ACA CTC TTT CCC TAC ACG ACG CTC TTC CGA TCT GGTAGCCA GGTGAGCTATAGCGTG |
| RP56_4 | Amplicon 5 fw / barcode 4 | ACA CTC TTT CCC TAC ACG ACG CTC TTC CGA TCT TCTATTCC GGTGAGCTATAGCGTG |
| RP57_9 | Amplicon 5 rv / barcode 9 | GAC TGG AGT TCA GAC GTG TGC TCT TCC GAT CT ACGGCAGT TATTGAGAGGGTGGTTTAA |
| **Sanger sequencing primers** | | |
| RP01 | pGAL1 fw | CTTTCAACATTTTCGGTTTG |
| RP02 | tADH1 rv | GCTTAAACACGTCTTTTCC |
| **Primers for cloning individual variants** | | |
| RP353 | R79A | ACCGGCGGCAGCGGCTATAGCAAAGCGGTGGCGGTGCGCGGCGAAGCGGATACCCCGAGCACCATTGTGGGCCAGCTGGTGGAAGCTGGCGGCTTTCAGGCGTGGGC |
| RP354 | E78K | ACCGGCGGCAGCGGCTATAGCAAAGCGGTGGCGGTGCGCGGCGAAGCGGATACCCCGAGCACCATTGTGGGCCAGCTGGTGAAACGCGGCGGCTTTCAGGCGTGGGC |
| RP355 | R165A | CCCGGGCGGCATGGCGGAAGCTGAAGATGTGGCGAGCTTTTATAGCCCGCTGCTGAACAACACCCTGTATGTGGGCGGCGATCATACCGCGGATTATGATAGCGAACTGGCG |
| RP356 | E164K | CCCGGGCGGCATGGCGAAACGCGAAGATGTGGCGAGCTTTTATAGCCCGCTGCTGAACAACACCCTGTATGTGGGCGGCGATCATACCGCGGATTATGATAGCGAACTGGCG |
| RP359 | ALFA-tag inserted at position 269 | CGTCTCTCATCGGTCTCAGGGC AAACGC CCA TCT AGA TTG GAA GAA GAA TTG AGA AGA AGA TTG ACT GAA CCA CAG GAT GGC GAA TGG GTG AGC TAT AGC GTG TAT GGC GAA AAC GTG GAT TATGAACGCT ATCCTGAGACCTGAGACGGCA |
| RP358 | Position 164 rv (type 3) | CTCAGGTCTCAGGATTTATTCCGCCATGCCG |
| RP126 | Position 55 fw (type 3b) | GCATCGTCTCATCGGTCTCATTCTAGCGGCTATAGCAAAG |
| RP134 | Position 219 rv (type 3) | ATGCCGTCTCAGGTCTCAGGATTTA CACAATGCCGGTGC |
| SB161 | Position 362 rv (type 3) | ATGCCGTCTCAGGTCTCAGGATTTAGCCGCTATCGC |
| RP330 | Position 65 fw (type 3) | CATCGGTCTCATATG GAAGCGGATACCCC |
| RP331 | Position 75 fw (type 3) | CATCGGTCTCATATG CAGCTGGTGGAACG |
| RP332 | Position 80 fw (type 3) | CATCGGTCTCATATG GGCGGCTTTCAGG |
| RP333 | Position 90 fw (type 3) | CATCGGTCTCATATG GGCATTTATCTGTTTGC |
| RP334 | Position 100 fw (type 3) | CATCGGTCTCATATG GATACCAGCAAAGTGAC |
| RP335 | Position 65 fw (type 3b) | CATCGGTCTCATTCT GAAGCGGATACCCC |
| RP336 | Position 75 fw (type 3b) | CATCGGTCTCATTCT CAGCTGGTGGAACG |
| RP337 | Position 80 fw (type 3b) | CATCGGTCTCATTCT GGCGGCTTTCAGG |
| RP338 | Position 90 fw (type 3b) | CATCGGTCTCATTCT GGCATTTATCTGTTTGC |
| RP339 | Position 100 fw (type 3b) | CATCGGTCTCATTCT GATACCAGCAAAGTGAC |
| RPA01 | L293A+G357A fw | ATACCTCTATACTTTAACGTCAAGGAG |
| RPA25 | L293A+G357A rv | GAACAAAAGCTGGAGCTCCACC |
| **Primers for verifying deletion mutants** | | |
| RP351 | To check ΔPBS2 locus | AAGGATCTTTCTAACGTGTGTTGTC |
| RP352 | To check ΔPBS2 locus | ACATCCCCTCAATACTCTGTCATAA |

**Supplementary Table S11.** Oligonucleotide pool sequences encoding the variants.

| **Position** | **Pool** | **Sequence** |
| --- | --- | --- |
| **Alanine scanning variants** | | |
| 1 | 1 | CCGAATTCCCAAAAGAA GCT AAA GAA ACC ACC ACC AGC CTG ATG CAG GAT GAA CTG ACC CTG GGC GAA CCG GCG ACC CAG GCG CGC ATG TGC GTG CGC CTGCTGCGCTTTTTTATTG |
| 2 | 1 | CCGAATTCCCAAAAGAA ATG GCT GAA ACC ACC ACC AGC CTG ATG CAG GAT GAA CTG ACC CTG GGC GAA CCG GCG ACC CAG GCG CGC ATG TGC GTG CGC CTGCTGCGCTTTTTTATTG |
| 3 | 1 | CCGAATTCCCAAAAGAA ATG AAA GCT ACC ACC ACC AGC CTG ATG CAG GAT GAA CTG ACC CTG GGC GAA CCG GCG ACC CAG GCG CGC ATG TGC GTG CGC CTGCTGCGCTTTTTTATTG |
| 4 | 1 | CCGAATTCCCAAAAGAA ATG AAA GAA GCT ACC ACC AGC CTG ATG CAG GAT GAA CTG ACC CTG GGC GAA CCG GCG ACC CAG GCG CGC ATG TGC GTG CGC CTGCTGCGCTTTTTTATTG |
| 5 | 1 | CCGAATTCCCAAAAGAA ATG AAA GAA ACC GCT ACC AGC CTG ATG CAG GAT GAA CTG ACC CTG GGC GAA CCG GCG ACC CAG GCG CGC ATG TGC GTG CGC CTGCTGCGCTTTTTTATTG |
| 6 | 1 | CCGAATTCCCAAAAGAA ATG AAA GAA ACC ACC GCT AGC CTG ATG CAG GAT GAA CTG ACC CTG GGC GAA CCG GCG ACC CAG GCG CGC ATG TGC GTG CGC CTGCTGCGCTTTTTTATTG |
| 7 | 1 | CCGAATTCCCAAAAGAA ATG AAA GAA ACC ACC ACC GCT CTG ATG CAG GAT GAA CTG ACC CTG GGC GAA CCG GCG ACC CAG GCG CGC ATG TGC GTG CGC CTGCTGCGCTTTTTTATTG |
| 8 | 1 | CCGAATTCCCAAAAGAA ATG AAA GAA ACC ACC ACC AGC GCT ATG CAG GAT GAA CTG ACC CTG GGC GAA CCG GCG ACC CAG GCG CGC ATG TGC GTG CGC CTGCTGCGCTTTTTTATTG |
| 9 | 1 | CCGAATTCCCAAAAGAA ATG AAA GAA ACC ACC ACC AGC CTG GCT CAG GAT GAA CTG ACC CTG GGC GAA CCG GCG ACC CAG GCG CGC ATG TGC GTG CGC CTGCTGCGCTTTTTTATTG |
| 10 | 1 | CCGAATTCCCAAAAGAA ATG AAA GAA ACC ACC ACC AGC CTG ATG GCT GAT GAA CTG ACC CTG GGC GAA CCG GCG ACC CAG GCG CGC ATG TGC GTG CGC CTGCTGCGCTTTTTTATTG |
| 11 | 1 | CCGAATTCCCAAAAGAA ATG AAA GAA ACC ACC ACC AGC CTG ATG CAG GCT GAA CTG ACC CTG GGC GAA CCG GCG ACC CAG GCG CGC ATG TGC GTG CGC CTGCTGCGCTTTTTTATTG |
| 12 | 1 | CCGAATTCCCAAAAGAA ATG AAA GAA ACC ACC ACC AGC CTG ATG CAG GAT GCT CTG ACC CTG GGC GAA CCG GCG ACC CAG GCG CGC ATG TGC GTG CGC CTGCTGCGCTTTTTTATTG |
| 13 | 1 | CCGAATTCCCAAAAGAA ATG AAA GAA ACC ACC ACC AGC CTG ATG CAG GAT GAA GCT ACC CTG GGC GAA CCG GCG ACC CAG GCG CGC ATG TGC GTG CGC CTGCTGCGCTTTTTTATTG |
| 14 | 1 | CCGAATTCCCAAAAGAA ATG AAA GAA ACC ACC ACC AGC CTG ATG CAG GAT GAA CTG GCT CTG GGC GAA CCG GCG ACC CAG GCG CGC ATG TGC GTG CGC CTGCTGCGCTTTTTTATTG |
| 15 | 1 | CCGAATTCCCAAAAGAA ATG AAA GAA ACC ACC ACC AGC CTG ATG CAG GAT GAA CTG ACC GCT GGC GAA CCG GCG ACC CAG GCG CGC ATG TGC GTG CGC CTGCTGCGCTTTTTTATTG |
| 16 | 1 | CCGAATTCCCAAAAGAA ATG AAA GAA ACC ACC ACC AGC CTG ATG CAG GAT GAA CTG ACC CTG GCT GAA CCG GCG ACC CAG GCG CGC ATG TGC GTG CGC CTGCTGCGCTTTTTTATTG |
| 17 | 1 | CCGAATTCCCAAAAGAA ATG AAA GAA ACC ACC ACC AGC CTG ATG CAG GAT GAA CTG ACC CTG GGC GCT CCG GCG ACC CAG GCG CGC ATG TGC GTG CGC CTGCTGCGCTTTTTTATTG |
| 18 | 1 | CCGAATTCCCAAAAGAA ATG AAA GAA ACC ACC ACC AGC CTG ATG CAG GAT GAA CTG ACC CTG GGC GAA GCT GCG ACC CAG GCG CGC ATG TGC GTG CGC CTGCTGCGCTTTTTTATTG |
| 19 | 1 | CCGAATTCCCAAAAGAA ATG AAA GAA ACC ACC ACC AGC CTG ATG CAG GAT GAA CTG ACC CTG GGC GAA CCG GGT ACC CAG GCG CGC ATG TGC GTG CGC CTGCTGCGCTTTTTTATTG |
| 20 | 1 | CCGAATTCCCAAAAGAA ATG AAA GAA ACC ACC ACC AGC CTG ATG CAG GAT GAA CTG ACC CTG GGC GAA CCG GCG GCT CAG GCG CGC ATG TGC GTG CGC CTGCTGCGCTTTTTTATTG |
| 21 | 1 | CCGAATTCCCAAAAGAA ATG AAA GAA ACC ACC ACC AGC CTG ATG CAG GAT GAA CTG ACC CTG GGC GAA CCG GCG ACC GCT GCG CGC ATG TGC GTG CGC CTGCTGCGCTTTTTTATTG |
| 22 | 1 | CCGAATTCCCAAAAGAA ATG AAA GAA ACC ACC ACC AGC CTG ATG CAG GAT GAA CTG ACC CTG GGC GAA CCG GCG ACC CAG GGT CGC ATG TGC GTG CGC CTGCTGCGCTTTTTTATTG |
| 23 | 1 | CCGAATTCCCAAAAGAA ATG AAA GAA ACC ACC ACC AGC CTG ATG CAG GAT GAA CTG ACC CTG GGC GAA CCG GCG ACC CAG GCG GCT ATG TGC GTG CGC CTGCTGCGCTTTTTTATTG |
| 24 | 1 | CCGAATTCCCAAAAGAA ATG AAA GAA ACC ACC ACC AGC CTG ATG CAG GAT GAA CTG ACC CTG GGC GAA CCG GCG ACC CAG GCG CGC GCT TGC GTG CGC CTGCTGCGCTTTTTTATTG |
| 25 | 1 | CCGAATTCCCAAAAGAA ATG AAA GAA ACC ACC ACC AGC CTG ATG CAG GAT GAA CTG ACC CTG GGC GAA CCG GCG ACC CAG GCG CGC ATG GCT GTG CGC CTGCTGCGCTTTTTTATTG |
| 26 | 1 | CCGAATTCCCAAAAGAA ATG AAA GAA ACC ACC ACC AGC CTG ATG CAG GAT GAA CTG ACC CTG GGC GAA CCG GCG ACC CAG GCG CGC ATG TGC GCT CGC CTGCTGCGCTTTTTTATTG |
| 27 | 1 | CCGAATTCCCAAAAGAA ATG AAA GAA ACC ACC ACC AGC CTG ATG CAG GAT GAA CTG ACC CTG GGC GAA CCG GCG ACC CAG GCG CGC ATG TGC GTG GCT CTGCTGCGCTTTTTTATTG |
| 28 | 2 | GCATGTGCGTGCGC GCT CTG CGC TTT TTT ATT GGC CTG ACC ATT ACC GCG TTT ATT ATT GCG GCG TGC ATT ATT AAA AGC GCG ACC GGC GGC AGC GGCTATAGCAAAGCGGT |
| 29 | 2 | GCATGTGCGTGCGC CTG GCT CGC TTT TTT ATT GGC CTG ACC ATT ACC GCG TTT ATT ATT GCG GCG TGC ATT ATT AAA AGC GCG ACC GGC GGC AGC GGCTATAGCAAAGCGGT |
| 30 | 2 | GCATGTGCGTGCGC CTG CTG GCT TTT TTT ATT GGC CTG ACC ATT ACC GCG TTT ATT ATT GCG GCG TGC ATT ATT AAA AGC GCG ACC GGC GGC AGC GGCTATAGCAAAGCGGT |
| 31 | 2 | GCATGTGCGTGCGC CTG CTG CGC GCT TTT ATT GGC CTG ACC ATT ACC GCG TTT ATT ATT GCG GCG TGC ATT ATT AAA AGC GCG ACC GGC GGC AGC GGCTATAGCAAAGCGGT |
| 32 | 2 | GCATGTGCGTGCGC CTG CTG CGC TTT GCT ATT GGC CTG ACC ATT ACC GCG TTT ATT ATT GCG GCG TGC ATT ATT AAA AGC GCG ACC GGC GGC AGC GGCTATAGCAAAGCGGT |
| 33 | 2 | GCATGTGCGTGCGC CTG CTG CGC TTT TTT GCT GGC CTG ACC ATT ACC GCG TTT ATT ATT GCG GCG TGC ATT ATT AAA AGC GCG ACC GGC GGC AGC GGCTATAGCAAAGCGGT |
| 34 | 2 | GCATGTGCGTGCGC CTG CTG CGC TTT TTT ATT GCT CTG ACC ATT ACC GCG TTT ATT ATT GCG GCG TGC ATT ATT AAA AGC GCG ACC GGC GGC AGC GGCTATAGCAAAGCGGT |
| 35 | 2 | GCATGTGCGTGCGC CTG CTG CGC TTT TTT ATT GGC GCT ACC ATT ACC GCG TTT ATT ATT GCG GCG TGC ATT ATT AAA AGC GCG ACC GGC GGC AGC GGCTATAGCAAAGCGGT |
| 36 | 2 | GCATGTGCGTGCGC CTG CTG CGC TTT TTT ATT GGC CTG GCT ATT ACC GCG TTT ATT ATT GCG GCG TGC ATT ATT AAA AGC GCG ACC GGC GGC AGC GGCTATAGCAAAGCGGT |
| 37 | 2 | GCATGTGCGTGCGC CTG CTG CGC TTT TTT ATT GGC CTG ACC GCT ACC GCG TTT ATT ATT GCG GCG TGC ATT ATT AAA AGC GCG ACC GGC GGC AGC GGCTATAGCAAAGCGGT |
| 38 | 2 | GCATGTGCGTGCGC CTG CTG CGC TTT TTT ATT GGC CTG ACC ATT GCT GCG TTT ATT ATT GCG GCG TGC ATT ATT AAA AGC GCG ACC GGC GGC AGC GGCTATAGCAAAGCGGT |
| 39 | 2 | GCATGTGCGTGCGC CTG CTG CGC TTT TTT ATT GGC CTG ACC ATT ACC GGT TTT ATT ATT GCG GCG TGC ATT ATT AAA AGC GCG ACC GGC GGC AGC GGCTATAGCAAAGCGGT |
| 40 | 2 | GCATGTGCGTGCGC CTG CTG CGC TTT TTT ATT GGC CTG ACC ATT ACC GCG GCT ATT ATT GCG GCG TGC ATT ATT AAA AGC GCG ACC GGC GGC AGC GGCTATAGCAAAGCGGT |
| 41 | 2 | GCATGTGCGTGCGC CTG CTG CGC TTT TTT ATT GGC CTG ACC ATT ACC GCG TTT GCT ATT GCG GCG TGC ATT ATT AAA AGC GCG ACC GGC GGC AGC GGCTATAGCAAAGCGGT |
| 42 | 2 | GCATGTGCGTGCGC CTG CTG CGC TTT TTT ATT GGC CTG ACC ATT ACC GCG TTT ATT GCT GCG GCG TGC ATT ATT AAA AGC GCG ACC GGC GGC AGC GGCTATAGCAAAGCGGT |
| 43 | 2 | GCATGTGCGTGCGC CTG CTG CGC TTT TTT ATT GGC CTG ACC ATT ACC GCG TTT ATT ATT GGT GCG TGC ATT ATT AAA AGC GCG ACC GGC GGC AGC GGCTATAGCAAAGCGGT |
| 44 | 2 | GCATGTGCGTGCGC CTG CTG CGC TTT TTT ATT GGC CTG ACC ATT ACC GCG TTT ATT ATT GCG GGT TGC ATT ATT AAA AGC GCG ACC GGC GGC AGC GGCTATAGCAAAGCGGT |
| 45 | 2 | GCATGTGCGTGCGC CTG CTG CGC TTT TTT ATT GGC CTG ACC ATT ACC GCG TTT ATT ATT GCG GCG GCT ATT ATT AAA AGC GCG ACC GGC GGC AGC GGCTATAGCAAAGCGGT |
| 46 | 2 | GCATGTGCGTGCGC CTG CTG CGC TTT TTT ATT GGC CTG ACC ATT ACC GCG TTT ATT ATT GCG GCG TGC GCT ATT AAA AGC GCG ACC GGC GGC AGC GGCTATAGCAAAGCGGT |
| 47 | 2 | GCATGTGCGTGCGC CTG CTG CGC TTT TTT ATT GGC CTG ACC ATT ACC GCG TTT ATT ATT GCG GCG TGC ATT GCT AAA AGC GCG ACC GGC GGC AGC GGCTATAGCAAAGCGGT |
| 48 | 2 | GCATGTGCGTGCGC CTG CTG CGC TTT TTT ATT GGC CTG ACC ATT ACC GCG TTT ATT ATT GCG GCG TGC ATT ATT GCT AGC GCG ACC GGC GGC AGC GGCTATAGCAAAGCGGT |
| 49 | 2 | GCATGTGCGTGCGC CTG CTG CGC TTT TTT ATT GGC CTG ACC ATT ACC GCG TTT ATT ATT GCG GCG TGC ATT ATT AAA GCT GCG ACC GGC GGC AGC GGCTATAGCAAAGCGGT |
| 50 | 2 | GCATGTGCGTGCGC CTG CTG CGC TTT TTT ATT GGC CTG ACC ATT ACC GCG TTT ATT ATT GCG GCG TGC ATT ATT AAA AGC GGT ACC GGC GGC AGC GGCTATAGCAAAGCGGT |
| 51 | 2 | GCATGTGCGTGCGC CTG CTG CGC TTT TTT ATT GGC CTG ACC ATT ACC GCG TTT ATT ATT GCG GCG TGC ATT ATT AAA AGC GCG GCT GGC GGC AGC GGCTATAGCAAAGCGGT |
| 52 | 2 | GCATGTGCGTGCGC CTG CTG CGC TTT TTT ATT GGC CTG ACC ATT ACC GCG TTT ATT ATT GCG GCG TGC ATT ATT AAA AGC GCG ACC GCT GGC AGC GGCTATAGCAAAGCGGT |
| 53 | 2 | GCATGTGCGTGCGC CTG CTG CGC TTT TTT ATT GGC CTG ACC ATT ACC GCG TTT ATT ATT GCG GCG TGC ATT ATT AAA AGC GCG ACC GGC GCT AGC GGCTATAGCAAAGCGGT |
| 54 | 2 | GCATGTGCGTGCGC CTG CTG CGC TTT TTT ATT GGC CTG ACC ATT ACC GCG TTT ATT ATT GCG GCG TGC ATT ATT AAA AGC GCG ACC GGC GGC GCT GGCTATAGCAAAGCGGT |
| 55 | 3 | ACCGGCGGCAGC GCT TAT AGC AAA GCG GTG GCG GTG CGC GGC GAA GCG GAT ACC CCG AGC ACC ATT GTG GGC CAG CTG GTG GAA CGC GGC GGC TTTCAGGCGTGGGC |
| 56 | 3 | ACCGGCGGCAGC GGC GCT AGC AAA GCG GTG GCG GTG CGC GGC GAA GCG GAT ACC CCG AGC ACC ATT GTG GGC CAG CTG GTG GAA CGC GGC GGC TTTCAGGCGTGGGC |
| 57 | 3 | ACCGGCGGCAGC GGC TAT GCT AAA GCG GTG GCG GTG CGC GGC GAA GCG GAT ACC CCG AGC ACC ATT GTG GGC CAG CTG GTG GAA CGC GGC GGC TTTCAGGCGTGGGC |
| 58 | 3 | ACCGGCGGCAGC GGC TAT AGC GCT GCG GTG GCG GTG CGC GGC GAA GCG GAT ACC CCG AGC ACC ATT GTG GGC CAG CTG GTG GAA CGC GGC GGC TTTCAGGCGTGGGC |
| 59 | 3 | ACCGGCGGCAGC GGC TAT AGC AAA GGT GTG GCG GTG CGC GGC GAA GCG GAT ACC CCG AGC ACC ATT GTG GGC CAG CTG GTG GAA CGC GGC GGC TTTCAGGCGTGGGC |
| 60 | 3 | ACCGGCGGCAGC GGC TAT AGC AAA GCG GCT GCG GTG CGC GGC GAA GCG GAT ACC CCG AGC ACC ATT GTG GGC CAG CTG GTG GAA CGC GGC GGC TTTCAGGCGTGGGC |
| 61 | 3 | ACCGGCGGCAGC GGC TAT AGC AAA GCG GTG GGT GTG CGC GGC GAA GCG GAT ACC CCG AGC ACC ATT GTG GGC CAG CTG GTG GAA CGC GGC GGC TTTCAGGCGTGGGC |
| 62 | 3 | ACCGGCGGCAGC GGC TAT AGC AAA GCG GTG GCG GCT CGC GGC GAA GCG GAT ACC CCG AGC ACC ATT GTG GGC CAG CTG GTG GAA CGC GGC GGC TTTCAGGCGTGGGC |
| 63 | 3 | ACCGGCGGCAGC GGC TAT AGC AAA GCG GTG GCG GTG GCT GGC GAA GCG GAT ACC CCG AGC ACC ATT GTG GGC CAG CTG GTG GAA CGC GGC GGC TTTCAGGCGTGGGC |
| 64 | 3 | ACCGGCGGCAGC GGC TAT AGC AAA GCG GTG GCG GTG CGC GCT GAA GCG GAT ACC CCG AGC ACC ATT GTG GGC CAG CTG GTG GAA CGC GGC GGC TTTCAGGCGTGGGC |
| 65 | 3 | ACCGGCGGCAGC GGC TAT AGC AAA GCG GTG GCG GTG CGC GGC GCT GCG GAT ACC CCG AGC ACC ATT GTG GGC CAG CTG GTG GAA CGC GGC GGC TTTCAGGCGTGGGC |
| 66 | 3 | ACCGGCGGCAGC GGC TAT AGC AAA GCG GTG GCG GTG CGC GGC GAA GGT GAT ACC CCG AGC ACC ATT GTG GGC CAG CTG GTG GAA CGC GGC GGC TTTCAGGCGTGGGC |
| 67 | 3 | ACCGGCGGCAGC GGC TAT AGC AAA GCG GTG GCG GTG CGC GGC GAA GCG GCT ACC CCG AGC ACC ATT GTG GGC CAG CTG GTG GAA CGC GGC GGC TTTCAGGCGTGGGC |
| 68 | 3 | ACCGGCGGCAGC GGC TAT AGC AAA GCG GTG GCG GTG CGC GGC GAA GCG GAT GCT CCG AGC ACC ATT GTG GGC CAG CTG GTG GAA CGC GGC GGC TTTCAGGCGTGGGC |
| 69 | 3 | ACCGGCGGCAGC GGC TAT AGC AAA GCG GTG GCG GTG CGC GGC GAA GCG GAT ACC GCT AGC ACC ATT GTG GGC CAG CTG GTG GAA CGC GGC GGC TTTCAGGCGTGGGC |
| 70 | 3 | ACCGGCGGCAGC GGC TAT AGC AAA GCG GTG GCG GTG CGC GGC GAA GCG GAT ACC CCG GCT ACC ATT GTG GGC CAG CTG GTG GAA CGC GGC GGC TTTCAGGCGTGGGC |
| 71 | 3 | ACCGGCGGCAGC GGC TAT AGC AAA GCG GTG GCG GTG CGC GGC GAA GCG GAT ACC CCG AGC GCT ATT GTG GGC CAG CTG GTG GAA CGC GGC GGC TTTCAGGCGTGGGC |
| 72 | 3 | ACCGGCGGCAGC GGC TAT AGC AAA GCG GTG GCG GTG CGC GGC GAA GCG GAT ACC CCG AGC ACC GCT GTG GGC CAG CTG GTG GAA CGC GGC GGC TTTCAGGCGTGGGC |
| 73 | 3 | ACCGGCGGCAGC GGC TAT AGC AAA GCG GTG GCG GTG CGC GGC GAA GCG GAT ACC CCG AGC ACC ATT GCT GGC CAG CTG GTG GAA CGC GGC GGC TTTCAGGCGTGGGC |
| 74 | 3 | ACCGGCGGCAGC GGC TAT AGC AAA GCG GTG GCG GTG CGC GGC GAA GCG GAT ACC CCG AGC ACC ATT GTG GCT CAG CTG GTG GAA CGC GGC GGC TTTCAGGCGTGGGC |
| 75 | 3 | ACCGGCGGCAGC GGC TAT AGC AAA GCG GTG GCG GTG CGC GGC GAA GCG GAT ACC CCG AGC ACC ATT GTG GGC GCT CTG GTG GAA CGC GGC GGC TTTCAGGCGTGGGC |
| 76 | 3 | ACCGGCGGCAGC GGC TAT AGC AAA GCG GTG GCG GTG CGC GGC GAA GCG GAT ACC CCG AGC ACC ATT GTG GGC CAG GCT GTG GAA CGC GGC GGC TTTCAGGCGTGGGC |
| 77 | 3 | ACCGGCGGCAGC GGC TAT AGC AAA GCG GTG GCG GTG CGC GGC GAA GCG GAT ACC CCG AGC ACC ATT GTG GGC CAG CTG GCT GAA CGC GGC GGC TTTCAGGCGTGGGC |
| 78 | 3 | ACCGGCGGCAGC GGC TAT AGC AAA GCG GTG GCG GTG CGC GGC GAA GCG GAT ACC CCG AGC ACC ATT GTG GGC CAG CTG GTG GCT CGC GGC GGC TTTCAGGCGTGGGC |
| 79 | 3 | ACCGGCGGCAGC GGC TAT AGC AAA GCG GTG GCG GTG CGC GGC GAA GCG GAT ACC CCG AGC ACC ATT GTG GGC CAG CTG GTG GAA GCT GGC GGC TTTCAGGCGTGGGC |
| 80 | 3 | ACCGGCGGCAGC GGC TAT AGC AAA GCG GTG GCG GTG CGC GGC GAA GCG GAT ACC CCG AGC ACC ATT GTG GGC CAG CTG GTG GAA CGC GCT GGC TTTCAGGCGTGGGC |
| 81 | 3 | ACCGGCGGCAGC GGC TAT AGC AAA GCG GTG GCG GTG CGC GGC GAA GCG GAT ACC CCG AGC ACC ATT GTG GGC CAG CTG GTG GAA CGC GGC GCT TTTCAGGCGTGGGC |
| 82 | 4 | TGGAACGCGGCGGC GCT CAG GCG TGG GCG GTG GGC GCG GGC ATT TAT CTG TTT GCG AAA ATT GCG TAT GAT ACC AGC AAA GTG ACC GCG GCG GTG TGCAACCCGGAAGCG |
| 83 | 4 | TGGAACGCGGCGGC TTT GCT GCG TGG GCG GTG GGC GCG GGC ATT TAT CTG TTT GCG AAA ATT GCG TAT GAT ACC AGC AAA GTG ACC GCG GCG GTG TGCAACCCGGAAGCG |
| 84 | 4 | TGGAACGCGGCGGC TTT CAG GGT TGG GCG GTG GGC GCG GGC ATT TAT CTG TTT GCG AAA ATT GCG TAT GAT ACC AGC AAA GTG ACC GCG GCG GTG TGCAACCCGGAAGCG |
| 85 | 4 | TGGAACGCGGCGGC TTT CAG GCG GCT GCG GTG GGC GCG GGC ATT TAT CTG TTT GCG AAA ATT GCG TAT GAT ACC AGC AAA GTG ACC GCG GCG GTG TGCAACCCGGAAGCG |
| 86 | 4 | TGGAACGCGGCGGC TTT CAG GCG TGG GGT GTG GGC GCG GGC ATT TAT CTG TTT GCG AAA ATT GCG TAT GAT ACC AGC AAA GTG ACC GCG GCG GTG TGCAACCCGGAAGCG |
| 87 | 4 | TGGAACGCGGCGGC TTT CAG GCG TGG GCG GCT GGC GCG GGC ATT TAT CTG TTT GCG AAA ATT GCG TAT GAT ACC AGC AAA GTG ACC GCG GCG GTG TGCAACCCGGAAGCG |
| 88 | 4 | TGGAACGCGGCGGC TTT CAG GCG TGG GCG GTG GCT GCG GGC ATT TAT CTG TTT GCG AAA ATT GCG TAT GAT ACC AGC AAA GTG ACC GCG GCG GTG TGCAACCCGGAAGCG |
| 89 | 4 | TGGAACGCGGCGGC TTT CAG GCG TGG GCG GTG GGC GGT GGC ATT TAT CTG TTT GCG AAA ATT GCG TAT GAT ACC AGC AAA GTG ACC GCG GCG GTG TGCAACCCGGAAGCG |
| 90 | 4 | TGGAACGCGGCGGC TTT CAG GCG TGG GCG GTG GGC GCG GCT ATT TAT CTG TTT GCG AAA ATT GCG TAT GAT ACC AGC AAA GTG ACC GCG GCG GTG TGCAACCCGGAAGCG |
| 91 | 4 | TGGAACGCGGCGGC TTT CAG GCG TGG GCG GTG GGC GCG GGC GCT TAT CTG TTT GCG AAA ATT GCG TAT GAT ACC AGC AAA GTG ACC GCG GCG GTG TGCAACCCGGAAGCG |
| 92 | 4 | TGGAACGCGGCGGC TTT CAG GCG TGG GCG GTG GGC GCG GGC ATT GCT CTG TTT GCG AAA ATT GCG TAT GAT ACC AGC AAA GTG ACC GCG GCG GTG TGCAACCCGGAAGCG |
| 93 | 4 | TGGAACGCGGCGGC TTT CAG GCG TGG GCG GTG GGC GCG GGC ATT TAT GCT TTT GCG AAA ATT GCG TAT GAT ACC AGC AAA GTG ACC GCG GCG GTG TGCAACCCGGAAGCG |
| 94 | 4 | TGGAACGCGGCGGC TTT CAG GCG TGG GCG GTG GGC GCG GGC ATT TAT CTG GCT GCG AAA ATT GCG TAT GAT ACC AGC AAA GTG ACC GCG GCG GTG TGCAACCCGGAAGCG |
| 95 | 4 | TGGAACGCGGCGGC TTT CAG GCG TGG GCG GTG GGC GCG GGC ATT TAT CTG TTT GGT AAA ATT GCG TAT GAT ACC AGC AAA GTG ACC GCG GCG GTG TGCAACCCGGAAGCG |
| 96 | 4 | TGGAACGCGGCGGC TTT CAG GCG TGG GCG GTG GGC GCG GGC ATT TAT CTG TTT GCG GCT ATT GCG TAT GAT ACC AGC AAA GTG ACC GCG GCG GTG TGCAACCCGGAAGCG |
| 97 | 4 | TGGAACGCGGCGGC TTT CAG GCG TGG GCG GTG GGC GCG GGC ATT TAT CTG TTT GCG AAA GCT GCG TAT GAT ACC AGC AAA GTG ACC GCG GCG GTG TGCAACCCGGAAGCG |
| 98 | 4 | TGGAACGCGGCGGC TTT CAG GCG TGG GCG GTG GGC GCG GGC ATT TAT CTG TTT GCG AAA ATT GGT TAT GAT ACC AGC AAA GTG ACC GCG GCG GTG TGCAACCCGGAAGCG |
| 99 | 4 | TGGAACGCGGCGGC TTT CAG GCG TGG GCG GTG GGC GCG GGC ATT TAT CTG TTT GCG AAA ATT GCG GCT GAT ACC AGC AAA GTG ACC GCG GCG GTG TGCAACCCGGAAGCG |
| 100 | 4 | TGGAACGCGGCGGC TTT CAG GCG TGG GCG GTG GGC GCG GGC ATT TAT CTG TTT GCG AAA ATT GCG TAT GCT ACC AGC AAA GTG ACC GCG GCG GTG TGCAACCCGGAAGCG |
| 101 | 4 | TGGAACGCGGCGGC TTT CAG GCG TGG GCG GTG GGC GCG GGC ATT TAT CTG TTT GCG AAA ATT GCG TAT GAT GCT AGC AAA GTG ACC GCG GCG GTG TGCAACCCGGAAGCG |
| 102 | 4 | TGGAACGCGGCGGC TTT CAG GCG TGG GCG GTG GGC GCG GGC ATT TAT CTG TTT GCG AAA ATT GCG TAT GAT ACC GCT AAA GTG ACC GCG GCG GTG TGCAACCCGGAAGCG |
| 103 | 4 | TGGAACGCGGCGGC TTT CAG GCG TGG GCG GTG GGC GCG GGC ATT TAT CTG TTT GCG AAA ATT GCG TAT GAT ACC AGC GCT GTG ACC GCG GCG GTG TGCAACCCGGAAGCG |
| 104 | 4 | TGGAACGCGGCGGC TTT CAG GCG TGG GCG GTG GGC GCG GGC ATT TAT CTG TTT GCG AAA ATT GCG TAT GAT ACC AGC AAA GCT ACC GCG GCG GTG TGCAACCCGGAAGCG |
| 105 | 4 | TGGAACGCGGCGGC TTT CAG GCG TGG GCG GTG GGC GCG GGC ATT TAT CTG TTT GCG AAA ATT GCG TAT GAT ACC AGC AAA GTG GCT GCG GCG GTG TGCAACCCGGAAGCG |
| 106 | 4 | TGGAACGCGGCGGC TTT CAG GCG TGG GCG GTG GGC GCG GGC ATT TAT CTG TTT GCG AAA ATT GCG TAT GAT ACC AGC AAA GTG ACC GGT GCG GTG TGCAACCCGGAAGCG |
| 107 | 4 | TGGAACGCGGCGGC TTT CAG GCG TGG GCG GTG GGC GCG GGC ATT TAT CTG TTT GCG AAA ATT GCG TAT GAT ACC AGC AAA GTG ACC GCG GGT GTG TGCAACCCGGAAGCG |
| 108 | 4 | TGGAACGCGGCGGC TTT CAG GCG TGG GCG GTG GGC GCG GGC ATT TAT CTG TTT GCG AAA ATT GCG TAT GAT ACC AGC AAA GTG ACC GCG GCG GCT TGCAACCCGGAAGCG |
| 109 | 5 | GACCGCGGCGGTG GCT AAC CCG GAA GCG CTG ATT GCG ATT ACC AGC TAT GTG GCG TAT GCG CCG ACC CTG TGC GCG GGC GCG TAT GTG ATT GGC GCGATGAGCGGCG |
| 110 | 5 | GACCGCGGCGGTG TGC GCT CCG GAA GCG CTG ATT GCG ATT ACC AGC TAT GTG GCG TAT GCG CCG ACC CTG TGC GCG GGC GCG TAT GTG ATT GGC GCGATGAGCGGCG |
| 111 | 5 | GACCGCGGCGGTG TGC AAC GCT GAA GCG CTG ATT GCG ATT ACC AGC TAT GTG GCG TAT GCG CCG ACC CTG TGC GCG GGC GCG TAT GTG ATT GGC GCGATGAGCGGCG |
| 112 | 5 | GACCGCGGCGGTG TGC AAC CCG GCT GCG CTG ATT GCG ATT ACC AGC TAT GTG GCG TAT GCG CCG ACC CTG TGC GCG GGC GCG TAT GTG ATT GGC GCGATGAGCGGCG |
| 113 | 5 | GACCGCGGCGGTG TGC AAC CCG GAA GGT CTG ATT GCG ATT ACC AGC TAT GTG GCG TAT GCG CCG ACC CTG TGC GCG GGC GCG TAT GTG ATT GGC GCGATGAGCGGCG |
| 114 | 5 | GACCGCGGCGGTG TGC AAC CCG GAA GCG GCT ATT GCG ATT ACC AGC TAT GTG GCG TAT GCG CCG ACC CTG TGC GCG GGC GCG TAT GTG ATT GGC GCGATGAGCGGCG |
| 115 | 5 | GACCGCGGCGGTG TGC AAC CCG GAA GCG CTG GCT GCG ATT ACC AGC TAT GTG GCG TAT GCG CCG ACC CTG TGC GCG GGC GCG TAT GTG ATT GGC GCGATGAGCGGCG |
| 116 | 5 | GACCGCGGCGGTG TGC AAC CCG GAA GCG CTG ATT GGT ATT ACC AGC TAT GTG GCG TAT GCG CCG ACC CTG TGC GCG GGC GCG TAT GTG ATT GGC GCGATGAGCGGCG |
| 117 | 5 | GACCGCGGCGGTG TGC AAC CCG GAA GCG CTG ATT GCG GCT ACC AGC TAT GTG GCG TAT GCG CCG ACC CTG TGC GCG GGC GCG TAT GTG ATT GGC GCGATGAGCGGCG |
| 118 | 5 | GACCGCGGCGGTG TGC AAC CCG GAA GCG CTG ATT GCG ATT GCT AGC TAT GTG GCG TAT GCG CCG ACC CTG TGC GCG GGC GCG TAT GTG ATT GGC GCGATGAGCGGCG |
| 119 | 5 | GACCGCGGCGGTG TGC AAC CCG GAA GCG CTG ATT GCG ATT ACC GCT TAT GTG GCG TAT GCG CCG ACC CTG TGC GCG GGC GCG TAT GTG ATT GGC GCGATGAGCGGCG |
| 120 | 5 | GACCGCGGCGGTG TGC AAC CCG GAA GCG CTG ATT GCG ATT ACC AGC GCT GTG GCG TAT GCG CCG ACC CTG TGC GCG GGC GCG TAT GTG ATT GGC GCGATGAGCGGCG |
| 121 | 5 | GACCGCGGCGGTG TGC AAC CCG GAA GCG CTG ATT GCG ATT ACC AGC TAT GCT GCG TAT GCG CCG ACC CTG TGC GCG GGC GCG TAT GTG ATT GGC GCGATGAGCGGCG |
| 122 | 5 | GACCGCGGCGGTG TGC AAC CCG GAA GCG CTG ATT GCG ATT ACC AGC TAT GTG GGT TAT GCG CCG ACC CTG TGC GCG GGC GCG TAT GTG ATT GGC GCGATGAGCGGCG |
| 123 | 5 | GACCGCGGCGGTG TGC AAC CCG GAA GCG CTG ATT GCG ATT ACC AGC TAT GTG GCG GCT GCG CCG ACC CTG TGC GCG GGC GCG TAT GTG ATT GGC GCGATGAGCGGCG |
| 124 | 5 | GACCGCGGCGGTG TGC AAC CCG GAA GCG CTG ATT GCG ATT ACC AGC TAT GTG GCG TAT GGT CCG ACC CTG TGC GCG GGC GCG TAT GTG ATT GGC GCGATGAGCGGCG |
| 125 | 5 | GACCGCGGCGGTG TGC AAC CCG GAA GCG CTG ATT GCG ATT ACC AGC TAT GTG GCG TAT GCG GCT ACC CTG TGC GCG GGC GCG TAT GTG ATT GGC GCGATGAGCGGCG |
| 126 | 5 | GACCGCGGCGGTG TGC AAC CCG GAA GCG CTG ATT GCG ATT ACC AGC TAT GTG GCG TAT GCG CCG GCT CTG TGC GCG GGC GCG TAT GTG ATT GGC GCGATGAGCGGCG |
| 127 | 5 | GACCGCGGCGGTG TGC AAC CCG GAA GCG CTG ATT GCG ATT ACC AGC TAT GTG GCG TAT GCG CCG ACC GCT TGC GCG GGC GCG TAT GTG ATT GGC GCGATGAGCGGCG |
| 128 | 5 | GACCGCGGCGGTG TGC AAC CCG GAA GCG CTG ATT GCG ATT ACC AGC TAT GTG GCG TAT GCG CCG ACC CTG GCT GCG GGC GCG TAT GTG ATT GGC GCGATGAGCGGCG |
| 129 | 5 | GACCGCGGCGGTG TGC AAC CCG GAA GCG CTG ATT GCG ATT ACC AGC TAT GTG GCG TAT GCG CCG ACC CTG TGC GGT GGC GCG TAT GTG ATT GGC GCGATGAGCGGCG |
| 130 | 5 | GACCGCGGCGGTG TGC AAC CCG GAA GCG CTG ATT GCG ATT ACC AGC TAT GTG GCG TAT GCG CCG ACC CTG TGC GCG GCT GCG TAT GTG ATT GGC GCGATGAGCGGCG |
| 131 | 5 | GACCGCGGCGGTG TGC AAC CCG GAA GCG CTG ATT GCG ATT ACC AGC TAT GTG GCG TAT GCG CCG ACC CTG TGC GCG GGC GGT TAT GTG ATT GGC GCGATGAGCGGCG |
| 132 | 5 | GACCGCGGCGGTG TGC AAC CCG GAA GCG CTG ATT GCG ATT ACC AGC TAT GTG GCG TAT GCG CCG ACC CTG TGC GCG GGC GCG GCT GTG ATT GGC GCGATGAGCGGCG |
| 133 | 5 | GACCGCGGCGGTG TGC AAC CCG GAA GCG CTG ATT GCG ATT ACC AGC TAT GTG GCG TAT GCG CCG ACC CTG TGC GCG GGC GCG TAT GCT ATT GGC GCGATGAGCGGCG |
| 134 | 5 | GACCGCGGCGGTG TGC AAC CCG GAA GCG CTG ATT GCG ATT ACC AGC TAT GTG GCG TAT GCG CCG ACC CTG TGC GCG GGC GCG TAT GTG GCT GGC GCGATGAGCGGCG |
| 135 | 5 | GACCGCGGCGGTG TGC AAC CCG GAA GCG CTG ATT GCG ATT ACC AGC TAT GTG GCG TAT GCG CCG ACC CTG TGC GCG GGC GCG TAT GTG ATT GCT GCGATGAGCGGCG |
| 136 | 6 | CGCGTATGTGATTGGC GGT ATG AGC GGC GCG ATG AGC GCG GGC CTG GCG CTG TAT GCG GGC TAT AAA GGC TGG CAG TGG GGC GGC CCG GGC GGC ATG GCGGAACGCGAAGATG |
| 137 | 6 | CGCGTATGTGATTGGC GCG GCT AGC GGC GCG ATG AGC GCG GGC CTG GCG CTG TAT GCG GGC TAT AAA GGC TGG CAG TGG GGC GGC CCG GGC GGC ATG GCGGAACGCGAAGATG |
| 138 | 6 | CGCGTATGTGATTGGC GCG ATG GCT GGC GCG ATG AGC GCG GGC CTG GCG CTG TAT GCG GGC TAT AAA GGC TGG CAG TGG GGC GGC CCG GGC GGC ATG GCGGAACGCGAAGATG |
| 139 | 6 | CGCGTATGTGATTGGC GCG ATG AGC GCT GCG ATG AGC GCG GGC CTG GCG CTG TAT GCG GGC TAT AAA GGC TGG CAG TGG GGC GGC CCG GGC GGC ATG GCGGAACGCGAAGATG |
| 140 | 6 | CGCGTATGTGATTGGC GCG ATG AGC GGC GGT ATG AGC GCG GGC CTG GCG CTG TAT GCG GGC TAT AAA GGC TGG CAG TGG GGC GGC CCG GGC GGC ATG GCGGAACGCGAAGATG |
| 141 | 6 | CGCGTATGTGATTGGC GCG ATG AGC GGC GCG GCT AGC GCG GGC CTG GCG CTG TAT GCG GGC TAT AAA GGC TGG CAG TGG GGC GGC CCG GGC GGC ATG GCGGAACGCGAAGATG |
| 142 | 6 | CGCGTATGTGATTGGC GCG ATG AGC GGC GCG ATG GCT GCG GGC CTG GCG CTG TAT GCG GGC TAT AAA GGC TGG CAG TGG GGC GGC CCG GGC GGC ATG GCGGAACGCGAAGATG |
| 143 | 6 | CGCGTATGTGATTGGC GCG ATG AGC GGC GCG ATG AGC GGT GGC CTG GCG CTG TAT GCG GGC TAT AAA GGC TGG CAG TGG GGC GGC CCG GGC GGC ATG GCGGAACGCGAAGATG |
| 144 | 6 | CGCGTATGTGATTGGC GCG ATG AGC GGC GCG ATG AGC GCG GCT CTG GCG CTG TAT GCG GGC TAT AAA GGC TGG CAG TGG GGC GGC CCG GGC GGC ATG GCGGAACGCGAAGATG |
| 145 | 6 | CGCGTATGTGATTGGC GCG ATG AGC GGC GCG ATG AGC GCG GGC GCT GCG CTG TAT GCG GGC TAT AAA GGC TGG CAG TGG GGC GGC CCG GGC GGC ATG GCGGAACGCGAAGATG |
| 146 | 6 | CGCGTATGTGATTGGC GCG ATG AGC GGC GCG ATG AGC GCG GGC CTG GGT CTG TAT GCG GGC TAT AAA GGC TGG CAG TGG GGC GGC CCG GGC GGC ATG GCGGAACGCGAAGATG |
| 147 | 6 | CGCGTATGTGATTGGC GCG ATG AGC GGC GCG ATG AGC GCG GGC CTG GCG GCT TAT GCG GGC TAT AAA GGC TGG CAG TGG GGC GGC CCG GGC GGC ATG GCGGAACGCGAAGATG |
| 148 | 6 | CGCGTATGTGATTGGC GCG ATG AGC GGC GCG ATG AGC GCG GGC CTG GCG CTG GCT GCG GGC TAT AAA GGC TGG CAG TGG GGC GGC CCG GGC GGC ATG GCGGAACGCGAAGATG |
| 149 | 6 | CGCGTATGTGATTGGC GCG ATG AGC GGC GCG ATG AGC GCG GGC CTG GCG CTG TAT GGT GGC TAT AAA GGC TGG CAG TGG GGC GGC CCG GGC GGC ATG GCGGAACGCGAAGATG |
| 150 | 6 | CGCGTATGTGATTGGC GCG ATG AGC GGC GCG ATG AGC GCG GGC CTG GCG CTG TAT GCG GCT TAT AAA GGC TGG CAG TGG GGC GGC CCG GGC GGC ATG GCGGAACGCGAAGATG |
| 151 | 6 | CGCGTATGTGATTGGC GCG ATG AGC GGC GCG ATG AGC GCG GGC CTG GCG CTG TAT GCG GGC GCT AAA GGC TGG CAG TGG GGC GGC CCG GGC GGC ATG GCGGAACGCGAAGATG |
| 152 | 6 | CGCGTATGTGATTGGC GCG ATG AGC GGC GCG ATG AGC GCG GGC CTG GCG CTG TAT GCG GGC TAT GCT GGC TGG CAG TGG GGC GGC CCG GGC GGC ATG GCGGAACGCGAAGATG |
| 153 | 6 | CGCGTATGTGATTGGC GCG ATG AGC GGC GCG ATG AGC GCG GGC CTG GCG CTG TAT GCG GGC TAT AAA GCT TGG CAG TGG GGC GGC CCG GGC GGC ATG GCGGAACGCGAAGATG |
| 154 | 6 | CGCGTATGTGATTGGC GCG ATG AGC GGC GCG ATG AGC GCG GGC CTG GCG CTG TAT GCG GGC TAT AAA GGC GCT CAG TGG GGC GGC CCG GGC GGC ATG GCGGAACGCGAAGATG |
| 155 | 6 | CGCGTATGTGATTGGC GCG ATG AGC GGC GCG ATG AGC GCG GGC CTG GCG CTG TAT GCG GGC TAT AAA GGC TGG GCT TGG GGC GGC CCG GGC GGC ATG GCGGAACGCGAAGATG |
| 156 | 6 | CGCGTATGTGATTGGC GCG ATG AGC GGC GCG ATG AGC GCG GGC CTG GCG CTG TAT GCG GGC TAT AAA GGC TGG CAG GCT GGC GGC CCG GGC GGC ATG GCGGAACGCGAAGATG |
| 157 | 6 | CGCGTATGTGATTGGC GCG ATG AGC GGC GCG ATG AGC GCG GGC CTG GCG CTG TAT GCG GGC TAT AAA GGC TGG CAG TGG GCT GGC CCG GGC GGC ATG GCGGAACGCGAAGATG |
| 158 | 6 | CGCGTATGTGATTGGC GCG ATG AGC GGC GCG ATG AGC GCG GGC CTG GCG CTG TAT GCG GGC TAT AAA GGC TGG CAG TGG GGC GCT CCG GGC GGC ATG GCGGAACGCGAAGATG |
| 159 | 6 | CGCGTATGTGATTGGC GCG ATG AGC GGC GCG ATG AGC GCG GGC CTG GCG CTG TAT GCG GGC TAT AAA GGC TGG CAG TGG GGC GGC GCT GGC GGC ATG GCGGAACGCGAAGATG |
| 160 | 6 | CGCGTATGTGATTGGC GCG ATG AGC GGC GCG ATG AGC GCG GGC CTG GCG CTG TAT GCG GGC TAT AAA GGC TGG CAG TGG GGC GGC CCG GCT GGC ATG GCGGAACGCGAAGATG |
| 161 | 6 | CGCGTATGTGATTGGC GCG ATG AGC GGC GCG ATG AGC GCG GGC CTG GCG CTG TAT GCG GGC TAT AAA GGC TGG CAG TGG GGC GGC CCG GGC GCT ATG GCGGAACGCGAAGATG |
| 162 | 6 | CGCGTATGTGATTGGC GCG ATG AGC GGC GCG ATG AGC GCG GGC CTG GCG CTG TAT GCG GGC TAT AAA GGC TGG CAG TGG GGC GGC CCG GGC GGC GCT GCGGAACGCGAAGATG |
| 163 | 7 | CCCGGGCGGCATG GGT GAA CGC GAA GAT GTG GCG AGC TTT TAT AGC CCG CTG CTG AAC AAC ACC CTG TAT GTG GGC GGC GAT CAT ACC GCG GAT TATGATAGCGAACTGGCG |
| 164 | 7 | CCCGGGCGGCATG GCG GCT CGC GAA GAT GTG GCG AGC TTT TAT AGC CCG CTG CTG AAC AAC ACC CTG TAT GTG GGC GGC GAT CAT ACC GCG GAT TATGATAGCGAACTGGCG |
| 165 | 7 | CCCGGGCGGCATG GCG GAA GCT GAA GAT GTG GCG AGC TTT TAT AGC CCG CTG CTG AAC AAC ACC CTG TAT GTG GGC GGC GAT CAT ACC GCG GAT TATGATAGCGAACTGGCG |
| 166 | 7 | CCCGGGCGGCATG GCG GAA CGC GCT GAT GTG GCG AGC TTT TAT AGC CCG CTG CTG AAC AAC ACC CTG TAT GTG GGC GGC GAT CAT ACC GCG GAT TATGATAGCGAACTGGCG |
| 167 | 7 | CCCGGGCGGCATG GCG GAA CGC GAA GCT GTG GCG AGC TTT TAT AGC CCG CTG CTG AAC AAC ACC CTG TAT GTG GGC GGC GAT CAT ACC GCG GAT TATGATAGCGAACTGGCG |
| 168 | 7 | CCCGGGCGGCATG GCG GAA CGC GAA GAT GCT GCG AGC TTT TAT AGC CCG CTG CTG AAC AAC ACC CTG TAT GTG GGC GGC GAT CAT ACC GCG GAT TATGATAGCGAACTGGCG |
| 169 | 7 | CCCGGGCGGCATG GCG GAA CGC GAA GAT GTG GGT AGC TTT TAT AGC CCG CTG CTG AAC AAC ACC CTG TAT GTG GGC GGC GAT CAT ACC GCG GAT TATGATAGCGAACTGGCG |
| 170 | 7 | CCCGGGCGGCATG GCG GAA CGC GAA GAT GTG GCG GCT TTT TAT AGC CCG CTG CTG AAC AAC ACC CTG TAT GTG GGC GGC GAT CAT ACC GCG GAT TATGATAGCGAACTGGCG |
| 171 | 7 | CCCGGGCGGCATG GCG GAA CGC GAA GAT GTG GCG AGC GCT TAT AGC CCG CTG CTG AAC AAC ACC CTG TAT GTG GGC GGC GAT CAT ACC GCG GAT TATGATAGCGAACTGGCG |
| 172 | 7 | CCCGGGCGGCATG GCG GAA CGC GAA GAT GTG GCG AGC TTT GCT AGC CCG CTG CTG AAC AAC ACC CTG TAT GTG GGC GGC GAT CAT ACC GCG GAT TATGATAGCGAACTGGCG |
| 173 | 7 | CCCGGGCGGCATG GCG GAA CGC GAA GAT GTG GCG AGC TTT TAT GCT CCG CTG CTG AAC AAC ACC CTG TAT GTG GGC GGC GAT CAT ACC GCG GAT TATGATAGCGAACTGGCG |
| 174 | 7 | CCCGGGCGGCATG GCG GAA CGC GAA GAT GTG GCG AGC TTT TAT AGC GCT CTG CTG AAC AAC ACC CTG TAT GTG GGC GGC GAT CAT ACC GCG GAT TATGATAGCGAACTGGCG |
| 175 | 7 | CCCGGGCGGCATG GCG GAA CGC GAA GAT GTG GCG AGC TTT TAT AGC CCG GCT CTG AAC AAC ACC CTG TAT GTG GGC GGC GAT CAT ACC GCG GAT TATGATAGCGAACTGGCG |
| 176 | 7 | CCCGGGCGGCATG GCG GAA CGC GAA GAT GTG GCG AGC TTT TAT AGC CCG CTG GCT AAC AAC ACC CTG TAT GTG GGC GGC GAT CAT ACC GCG GAT TATGATAGCGAACTGGCG |
| 177 | 7 | CCCGGGCGGCATG GCG GAA CGC GAA GAT GTG GCG AGC TTT TAT AGC CCG CTG CTG GCT AAC ACC CTG TAT GTG GGC GGC GAT CAT ACC GCG GAT TATGATAGCGAACTGGCG |
| 178 | 7 | CCCGGGCGGCATG GCG GAA CGC GAA GAT GTG GCG AGC TTT TAT AGC CCG CTG CTG AAC GCT ACC CTG TAT GTG GGC GGC GAT CAT ACC GCG GAT TATGATAGCGAACTGGCG |
| 179 | 7 | CCCGGGCGGCATG GCG GAA CGC GAA GAT GTG GCG AGC TTT TAT AGC CCG CTG CTG AAC AAC GCT CTG TAT GTG GGC GGC GAT CAT ACC GCG GAT TATGATAGCGAACTGGCG |
| 180 | 7 | CCCGGGCGGCATG GCG GAA CGC GAA GAT GTG GCG AGC TTT TAT AGC CCG CTG CTG AAC AAC ACC GCT TAT GTG GGC GGC GAT CAT ACC GCG GAT TATGATAGCGAACTGGCG |
| 181 | 7 | CCCGGGCGGCATG GCG GAA CGC GAA GAT GTG GCG AGC TTT TAT AGC CCG CTG CTG AAC AAC ACC CTG GCT GTG GGC GGC GAT CAT ACC GCG GAT TATGATAGCGAACTGGCG |
| 182 | 7 | CCCGGGCGGCATG GCG GAA CGC GAA GAT GTG GCG AGC TTT TAT AGC CCG CTG CTG AAC AAC ACC CTG TAT GCT GGC GGC GAT CAT ACC GCG GAT TATGATAGCGAACTGGCG |
| 183 | 7 | CCCGGGCGGCATG GCG GAA CGC GAA GAT GTG GCG AGC TTT TAT AGC CCG CTG CTG AAC AAC ACC CTG TAT GTG GCT GGC GAT CAT ACC GCG GAT TATGATAGCGAACTGGCG |
| 184 | 7 | CCCGGGCGGCATG GCG GAA CGC GAA GAT GTG GCG AGC TTT TAT AGC CCG CTG CTG AAC AAC ACC CTG TAT GTG GGC GCT GAT CAT ACC GCG GAT TATGATAGCGAACTGGCG |
| 185 | 7 | CCCGGGCGGCATG GCG GAA CGC GAA GAT GTG GCG AGC TTT TAT AGC CCG CTG CTG AAC AAC ACC CTG TAT GTG GGC GGC GCT CAT ACC GCG GAT TATGATAGCGAACTGGCG |
| 186 | 7 | CCCGGGCGGCATG GCG GAA CGC GAA GAT GTG GCG AGC TTT TAT AGC CCG CTG CTG AAC AAC ACC CTG TAT GTG GGC GGC GAT GCT ACC GCG GAT TATGATAGCGAACTGGCG |
| 187 | 7 | CCCGGGCGGCATG GCG GAA CGC GAA GAT GTG GCG AGC TTT TAT AGC CCG CTG CTG AAC AAC ACC CTG TAT GTG GGC GGC GAT CAT GCT GCG GAT TATGATAGCGAACTGGCG |
| 188 | 7 | CCCGGGCGGCATG GCG GAA CGC GAA GAT GTG GCG AGC TTT TAT AGC CCG CTG CTG AAC AAC ACC CTG TAT GTG GGC GGC GAT CAT ACC GGT GAT TATGATAGCGAACTGGCG |
| 189 | 7 | CCCGGGCGGCATG GCG GAA CGC GAA GAT GTG GCG AGC TTT TAT AGC CCG CTG CTG AAC AAC ACC CTG TAT GTG GGC GGC GAT CAT ACC GCG GCT TATGATAGCGAACTGGCG |
| 190 | 8 | GCGATCATACCGCGGAT GCT GAT AGC GAA CTG GCG ACC ATT CTG GGC AGC GTG TAT AAC GAT GTG GTG CAT CTG GGC GTG TAT TAT GAT AAC AGC ACC GGCATTGTGAAACGCGATAG |
| 191 | 8 | GCGATCATACCGCGGAT TAT GCT AGC GAA CTG GCG ACC ATT CTG GGC AGC GTG TAT AAC GAT GTG GTG CAT CTG GGC GTG TAT TAT GAT AAC AGC ACC GGCATTGTGAAACGCGATAG |
| 192 | 8 | GCGATCATACCGCGGAT TAT GAT GCT GAA CTG GCG ACC ATT CTG GGC AGC GTG TAT AAC GAT GTG GTG CAT CTG GGC GTG TAT TAT GAT AAC AGC ACC GGCATTGTGAAACGCGATAG |
| 193 | 8 | GCGATCATACCGCGGAT TAT GAT AGC GCT CTG GCG ACC ATT CTG GGC AGC GTG TAT AAC GAT GTG GTG CAT CTG GGC GTG TAT TAT GAT AAC AGC ACC GGCATTGTGAAACGCGATAG |
| 194 | 8 | GCGATCATACCGCGGAT TAT GAT AGC GAA GCT GCG ACC ATT CTG GGC AGC GTG TAT AAC GAT GTG GTG CAT CTG GGC GTG TAT TAT GAT AAC AGC ACC GGCATTGTGAAACGCGATAG |
| 195 | 8 | GCGATCATACCGCGGAT TAT GAT AGC GAA CTG GGT ACC ATT CTG GGC AGC GTG TAT AAC GAT GTG GTG CAT CTG GGC GTG TAT TAT GAT AAC AGC ACC GGCATTGTGAAACGCGATAG |
| 196 | 8 | GCGATCATACCGCGGAT TAT GAT AGC GAA CTG GCG GCT ATT CTG GGC AGC GTG TAT AAC GAT GTG GTG CAT CTG GGC GTG TAT TAT GAT AAC AGC ACC GGCATTGTGAAACGCGATAG |
| 197 | 8 | GCGATCATACCGCGGAT TAT GAT AGC GAA CTG GCG ACC GCT CTG GGC AGC GTG TAT AAC GAT GTG GTG CAT CTG GGC GTG TAT TAT GAT AAC AGC ACC GGCATTGTGAAACGCGATAG |
| 198 | 8 | GCGATCATACCGCGGAT TAT GAT AGC GAA CTG GCG ACC ATT GCT GGC AGC GTG TAT AAC GAT GTG GTG CAT CTG GGC GTG TAT TAT GAT AAC AGC ACC GGCATTGTGAAACGCGATAG |
| 199 | 8 | GCGATCATACCGCGGAT TAT GAT AGC GAA CTG GCG ACC ATT CTG GCT AGC GTG TAT AAC GAT GTG GTG CAT CTG GGC GTG TAT TAT GAT AAC AGC ACC GGCATTGTGAAACGCGATAG |
| 200 | 8 | GCGATCATACCGCGGAT TAT GAT AGC GAA CTG GCG ACC ATT CTG GGC GCT GTG TAT AAC GAT GTG GTG CAT CTG GGC GTG TAT TAT GAT AAC AGC ACC GGCATTGTGAAACGCGATAG |
| 201 | 8 | GCGATCATACCGCGGAT TAT GAT AGC GAA CTG GCG ACC ATT CTG GGC AGC GCT TAT AAC GAT GTG GTG CAT CTG GGC GTG TAT TAT GAT AAC AGC ACC GGCATTGTGAAACGCGATAG |
| 202 | 8 | GCGATCATACCGCGGAT TAT GAT AGC GAA CTG GCG ACC ATT CTG GGC AGC GTG GCT AAC GAT GTG GTG CAT CTG GGC GTG TAT TAT GAT AAC AGC ACC GGCATTGTGAAACGCGATAG |
| 203 | 8 | GCGATCATACCGCGGAT TAT GAT AGC GAA CTG GCG ACC ATT CTG GGC AGC GTG TAT GCT GAT GTG GTG CAT CTG GGC GTG TAT TAT GAT AAC AGC ACC GGCATTGTGAAACGCGATAG |
| 204 | 8 | GCGATCATACCGCGGAT TAT GAT AGC GAA CTG GCG ACC ATT CTG GGC AGC GTG TAT AAC GCT GTG GTG CAT CTG GGC GTG TAT TAT GAT AAC AGC ACC GGCATTGTGAAACGCGATAG |
| 205 | 8 | GCGATCATACCGCGGAT TAT GAT AGC GAA CTG GCG ACC ATT CTG GGC AGC GTG TAT AAC GAT GCT GTG CAT CTG GGC GTG TAT TAT GAT AAC AGC ACC GGCATTGTGAAACGCGATAG |
| 206 | 8 | GCGATCATACCGCGGAT TAT GAT AGC GAA CTG GCG ACC ATT CTG GGC AGC GTG TAT AAC GAT GTG GCT CAT CTG GGC GTG TAT TAT GAT AAC AGC ACC GGCATTGTGAAACGCGATAG |
| 207 | 8 | GCGATCATACCGCGGAT TAT GAT AGC GAA CTG GCG ACC ATT CTG GGC AGC GTG TAT AAC GAT GTG GTG GCT CTG GGC GTG TAT TAT GAT AAC AGC ACC GGCATTGTGAAACGCGATAG |
| 208 | 8 | GCGATCATACCGCGGAT TAT GAT AGC GAA CTG GCG ACC ATT CTG GGC AGC GTG TAT AAC GAT GTG GTG CAT GCT GGC GTG TAT TAT GAT AAC AGC ACC GGCATTGTGAAACGCGATAG |
| 209 | 8 | GCGATCATACCGCGGAT TAT GAT AGC GAA CTG GCG ACC ATT CTG GGC AGC GTG TAT AAC GAT GTG GTG CAT CTG GCT GTG TAT TAT GAT AAC AGC ACC GGCATTGTGAAACGCGATAG |
| 210 | 8 | GCGATCATACCGCGGAT TAT GAT AGC GAA CTG GCG ACC ATT CTG GGC AGC GTG TAT AAC GAT GTG GTG CAT CTG GGC GCT TAT TAT GAT AAC AGC ACC GGCATTGTGAAACGCGATAG |
| 211 | 8 | GCGATCATACCGCGGAT TAT GAT AGC GAA CTG GCG ACC ATT CTG GGC AGC GTG TAT AAC GAT GTG GTG CAT CTG GGC GTG GCT TAT GAT AAC AGC ACC GGCATTGTGAAACGCGATAG |
| 212 | 8 | GCGATCATACCGCGGAT TAT GAT AGC GAA CTG GCG ACC ATT CTG GGC AGC GTG TAT AAC GAT GTG GTG CAT CTG GGC GTG TAT GCT GAT AAC AGC ACC GGCATTGTGAAACGCGATAG |
| 213 | 8 | GCGATCATACCGCGGAT TAT GAT AGC GAA CTG GCG ACC ATT CTG GGC AGC GTG TAT AAC GAT GTG GTG CAT CTG GGC GTG TAT TAT GCT AAC AGC ACC GGCATTGTGAAACGCGATAG |
| 214 | 8 | GCGATCATACCGCGGAT TAT GAT AGC GAA CTG GCG ACC ATT CTG GGC AGC GTG TAT AAC GAT GTG GTG CAT CTG GGC GTG TAT TAT GAT GCT AGC ACC GGCATTGTGAAACGCGATAG |
| 215 | 8 | GCGATCATACCGCGGAT TAT GAT AGC GAA CTG GCG ACC ATT CTG GGC AGC GTG TAT AAC GAT GTG GTG CAT CTG GGC GTG TAT TAT GAT AAC GCT ACC GGCATTGTGAAACGCGATAG |
| 216 | 8 | GCGATCATACCGCGGAT TAT GAT AGC GAA CTG GCG ACC ATT CTG GGC AGC GTG TAT AAC GAT GTG GTG CAT CTG GGC GTG TAT TAT GAT AAC AGC GCT GGCATTGTGAAACGCGATAG |
| 217 | 9 | GTGTATTATGATAACAGCACC GCT ATT GTG AAA CGC GAT AGC CGC CCG AGC ATG ATT AGC TGG ACC GTG CTG CAT GAT AAC ATG ATG ATT ACC AGC TAT CAT CGCCCGGATCAGCT |
| 218 | 9 | GTGTATTATGATAACAGCACC GGC GCT GTG AAA CGC GAT AGC CGC CCG AGC ATG ATT AGC TGG ACC GTG CTG CAT GAT AAC ATG ATG ATT ACC AGC TAT CAT CGCCCGGATCAGCT |
| 219 | 9 | GTGTATTATGATAACAGCACC GGC ATT GCT AAA CGC GAT AGC CGC CCG AGC ATG ATT AGC TGG ACC GTG CTG CAT GAT AAC ATG ATG ATT ACC AGC TAT CAT CGCCCGGATCAGCT |
| 220 | 9 | GTGTATTATGATAACAGCACC GGC ATT GTG GCT CGC GAT AGC CGC CCG AGC ATG ATT AGC TGG ACC GTG CTG CAT GAT AAC ATG ATG ATT ACC AGC TAT CAT CGCCCGGATCAGCT |
| 221 | 9 | GTGTATTATGATAACAGCACC GGC ATT GTG AAA GCT GAT AGC CGC CCG AGC ATG ATT AGC TGG ACC GTG CTG CAT GAT AAC ATG ATG ATT ACC AGC TAT CAT CGCCCGGATCAGCT |
| 222 | 9 | GTGTATTATGATAACAGCACC GGC ATT GTG AAA CGC GCT AGC CGC CCG AGC ATG ATT AGC TGG ACC GTG CTG CAT GAT AAC ATG ATG ATT ACC AGC TAT CAT CGCCCGGATCAGCT |
| 223 | 9 | GTGTATTATGATAACAGCACC GGC ATT GTG AAA CGC GAT GCT CGC CCG AGC ATG ATT AGC TGG ACC GTG CTG CAT GAT AAC ATG ATG ATT ACC AGC TAT CAT CGCCCGGATCAGCT |
| 224 | 9 | GTGTATTATGATAACAGCACC GGC ATT GTG AAA CGC GAT AGC GCT CCG AGC ATG ATT AGC TGG ACC GTG CTG CAT GAT AAC ATG ATG ATT ACC AGC TAT CAT CGCCCGGATCAGCT |
| 225 | 9 | GTGTATTATGATAACAGCACC GGC ATT GTG AAA CGC GAT AGC CGC GCT AGC ATG ATT AGC TGG ACC GTG CTG CAT GAT AAC ATG ATG ATT ACC AGC TAT CAT CGCCCGGATCAGCT |
| 226 | 9 | GTGTATTATGATAACAGCACC GGC ATT GTG AAA CGC GAT AGC CGC CCG GCT ATG ATT AGC TGG ACC GTG CTG CAT GAT AAC ATG ATG ATT ACC AGC TAT CAT CGCCCGGATCAGCT |
| 227 | 9 | GTGTATTATGATAACAGCACC GGC ATT GTG AAA CGC GAT AGC CGC CCG AGC GCT ATT AGC TGG ACC GTG CTG CAT GAT AAC ATG ATG ATT ACC AGC TAT CAT CGCCCGGATCAGCT |
| 228 | 9 | GTGTATTATGATAACAGCACC GGC ATT GTG AAA CGC GAT AGC CGC CCG AGC ATG GCT AGC TGG ACC GTG CTG CAT GAT AAC ATG ATG ATT ACC AGC TAT CAT CGCCCGGATCAGCT |
| 229 | 9 | GTGTATTATGATAACAGCACC GGC ATT GTG AAA CGC GAT AGC CGC CCG AGC ATG ATT GCT TGG ACC GTG CTG CAT GAT AAC ATG ATG ATT ACC AGC TAT CAT CGCCCGGATCAGCT |
| 230 | 9 | GTGTATTATGATAACAGCACC GGC ATT GTG AAA CGC GAT AGC CGC CCG AGC ATG ATT AGC GCT ACC GTG CTG CAT GAT AAC ATG ATG ATT ACC AGC TAT CAT CGCCCGGATCAGCT |
| 231 | 9 | GTGTATTATGATAACAGCACC GGC ATT GTG AAA CGC GAT AGC CGC CCG AGC ATG ATT AGC TGG GCT GTG CTG CAT GAT AAC ATG ATG ATT ACC AGC TAT CAT CGCCCGGATCAGCT |
| 232 | 9 | GTGTATTATGATAACAGCACC GGC ATT GTG AAA CGC GAT AGC CGC CCG AGC ATG ATT AGC TGG ACC GCT CTG CAT GAT AAC ATG ATG ATT ACC AGC TAT CAT CGCCCGGATCAGCT |
| 233 | 9 | GTGTATTATGATAACAGCACC GGC ATT GTG AAA CGC GAT AGC CGC CCG AGC ATG ATT AGC TGG ACC GTG GCT CAT GAT AAC ATG ATG ATT ACC AGC TAT CAT CGCCCGGATCAGCT |
| 234 | 9 | GTGTATTATGATAACAGCACC GGC ATT GTG AAA CGC GAT AGC CGC CCG AGC ATG ATT AGC TGG ACC GTG CTG GCT GAT AAC ATG ATG ATT ACC AGC TAT CAT CGCCCGGATCAGCT |
| 235 | 9 | GTGTATTATGATAACAGCACC GGC ATT GTG AAA CGC GAT AGC CGC CCG AGC ATG ATT AGC TGG ACC GTG CTG CAT GCT AAC ATG ATG ATT ACC AGC TAT CAT CGCCCGGATCAGCT |
| 236 | 9 | GTGTATTATGATAACAGCACC GGC ATT GTG AAA CGC GAT AGC CGC CCG AGC ATG ATT AGC TGG ACC GTG CTG CAT GAT GCT ATG ATG ATT ACC AGC TAT CAT CGCCCGGATCAGCT |
| 237 | 9 | GTGTATTATGATAACAGCACC GGC ATT GTG AAA CGC GAT AGC CGC CCG AGC ATG ATT AGC TGG ACC GTG CTG CAT GAT AAC GCT ATG ATT ACC AGC TAT CAT CGCCCGGATCAGCT |
| 238 | 9 | GTGTATTATGATAACAGCACC GGC ATT GTG AAA CGC GAT AGC CGC CCG AGC ATG ATT AGC TGG ACC GTG CTG CAT GAT AAC ATG GCT ATT ACC AGC TAT CAT CGCCCGGATCAGCT |
| 239 | 9 | GTGTATTATGATAACAGCACC GGC ATT GTG AAA CGC GAT AGC CGC CCG AGC ATG ATT AGC TGG ACC GTG CTG CAT GAT AAC ATG ATG GCT ACC AGC TAT CAT CGCCCGGATCAGCT |
| 240 | 9 | GTGTATTATGATAACAGCACC GGC ATT GTG AAA CGC GAT AGC CGC CCG AGC ATG ATT AGC TGG ACC GTG CTG CAT GAT AAC ATG ATG ATT GCT AGC TAT CAT CGCCCGGATCAGCT |
| 241 | 9 | GTGTATTATGATAACAGCACC GGC ATT GTG AAA CGC GAT AGC CGC CCG AGC ATG ATT AGC TGG ACC GTG CTG CAT GAT AAC ATG ATG ATT ACC GCT TAT CAT CGCCCGGATCAGCT |
| 242 | 9 | GTGTATTATGATAACAGCACC GGC ATT GTG AAA CGC GAT AGC CGC CCG AGC ATG ATT AGC TGG ACC GTG CTG CAT GAT AAC ATG ATG ATT ACC AGC GCT CAT CGCCCGGATCAGCT |
| 243 | 9 | GTGTATTATGATAACAGCACC GGC ATT GTG AAA CGC GAT AGC CGC CCG AGC ATG ATT AGC TGG ACC GTG CTG CAT GAT AAC ATG ATG ATT ACC AGC TAT GCT CGCCCGGATCAGCT |
| 244 | 10 | CATGATGATTACCAGCTATCAT GCT CCG GAT CAG CTG GGC GCG GCG GCG ACC GCG TAT AAA GCG TAT ACC ACC AAC ACC ACC CGC GTG GGC AAA CGC CAGGATGGCGAATGGG |
| 245 | 10 | CATGATGATTACCAGCTATCAT CGC GCT GAT CAG CTG GGC GCG GCG GCG ACC GCG TAT AAA GCG TAT ACC ACC AAC ACC ACC CGC GTG GGC AAA CGC CAGGATGGCGAATGGG |
| 246 | 10 | CATGATGATTACCAGCTATCAT CGC CCG GCT CAG CTG GGC GCG GCG GCG ACC GCG TAT AAA GCG TAT ACC ACC AAC ACC ACC CGC GTG GGC AAA CGC CAGGATGGCGAATGGG |
| 247 | 10 | CATGATGATTACCAGCTATCAT CGC CCG GAT GCT CTG GGC GCG GCG GCG ACC GCG TAT AAA GCG TAT ACC ACC AAC ACC ACC CGC GTG GGC AAA CGC CAGGATGGCGAATGGG |
| 248 | 10 | CATGATGATTACCAGCTATCAT CGC CCG GAT CAG GCT GGC GCG GCG GCG ACC GCG TAT AAA GCG TAT ACC ACC AAC ACC ACC CGC GTG GGC AAA CGC CAGGATGGCGAATGGG |
| 249 | 10 | CATGATGATTACCAGCTATCAT CGC CCG GAT CAG CTG GCT GCG GCG GCG ACC GCG TAT AAA GCG TAT ACC ACC AAC ACC ACC CGC GTG GGC AAA CGC CAGGATGGCGAATGGG |
| 250 | 10 | CATGATGATTACCAGCTATCAT CGC CCG GAT CAG CTG GGC GGT GCG GCG ACC GCG TAT AAA GCG TAT ACC ACC AAC ACC ACC CGC GTG GGC AAA CGC CAGGATGGCGAATGGG |
| 251 | 10 | CATGATGATTACCAGCTATCAT CGC CCG GAT CAG CTG GGC GCG GGT GCG ACC GCG TAT AAA GCG TAT ACC ACC AAC ACC ACC CGC GTG GGC AAA CGC CAGGATGGCGAATGGG |
| 252 | 10 | CATGATGATTACCAGCTATCAT CGC CCG GAT CAG CTG GGC GCG GCG GGT ACC GCG TAT AAA GCG TAT ACC ACC AAC ACC ACC CGC GTG GGC AAA CGC CAGGATGGCGAATGGG |
| 253 | 10 | CATGATGATTACCAGCTATCAT CGC CCG GAT CAG CTG GGC GCG GCG GCG GCT GCG TAT AAA GCG TAT ACC ACC AAC ACC ACC CGC GTG GGC AAA CGC CAGGATGGCGAATGGG |
| 254 | 10 | CATGATGATTACCAGCTATCAT CGC CCG GAT CAG CTG GGC GCG GCG GCG ACC GGT TAT AAA GCG TAT ACC ACC AAC ACC ACC CGC GTG GGC AAA CGC CAGGATGGCGAATGGG |
| 255 | 10 | CATGATGATTACCAGCTATCAT CGC CCG GAT CAG CTG GGC GCG GCG GCG ACC GCG GCT AAA GCG TAT ACC ACC AAC ACC ACC CGC GTG GGC AAA CGC CAGGATGGCGAATGGG |
| 256 | 10 | CATGATGATTACCAGCTATCAT CGC CCG GAT CAG CTG GGC GCG GCG GCG ACC GCG TAT GCT GCG TAT ACC ACC AAC ACC ACC CGC GTG GGC AAA CGC CAGGATGGCGAATGGG |
| 257 | 10 | CATGATGATTACCAGCTATCAT CGC CCG GAT CAG CTG GGC GCG GCG GCG ACC GCG TAT AAA GGT TAT ACC ACC AAC ACC ACC CGC GTG GGC AAA CGC CAGGATGGCGAATGGG |
| 258 | 10 | CATGATGATTACCAGCTATCAT CGC CCG GAT CAG CTG GGC GCG GCG GCG ACC GCG TAT AAA GCG GCT ACC ACC AAC ACC ACC CGC GTG GGC AAA CGC CAGGATGGCGAATGGG |
| 259 | 10 | CATGATGATTACCAGCTATCAT CGC CCG GAT CAG CTG GGC GCG GCG GCG ACC GCG TAT AAA GCG TAT GCT ACC AAC ACC ACC CGC GTG GGC AAA CGC CAGGATGGCGAATGGG |
| 260 | 10 | CATGATGATTACCAGCTATCAT CGC CCG GAT CAG CTG GGC GCG GCG GCG ACC GCG TAT AAA GCG TAT ACC GCT AAC ACC ACC CGC GTG GGC AAA CGC CAGGATGGCGAATGGG |
| 261 | 10 | CATGATGATTACCAGCTATCAT CGC CCG GAT CAG CTG GGC GCG GCG GCG ACC GCG TAT AAA GCG TAT ACC ACC GCT ACC ACC CGC GTG GGC AAA CGC CAGGATGGCGAATGGG |
| 262 | 10 | CATGATGATTACCAGCTATCAT CGC CCG GAT CAG CTG GGC GCG GCG GCG ACC GCG TAT AAA GCG TAT ACC ACC AAC GCT ACC CGC GTG GGC AAA CGC CAGGATGGCGAATGGG |
| 263 | 10 | CATGATGATTACCAGCTATCAT CGC CCG GAT CAG CTG GGC GCG GCG GCG ACC GCG TAT AAA GCG TAT ACC ACC AAC ACC GCT CGC GTG GGC AAA CGC CAGGATGGCGAATGGG |
| 264 | 10 | CATGATGATTACCAGCTATCAT CGC CCG GAT CAG CTG GGC GCG GCG GCG ACC GCG TAT AAA GCG TAT ACC ACC AAC ACC ACC GCT GTG GGC AAA CGC CAGGATGGCGAATGGG |
| 265 | 10 | CATGATGATTACCAGCTATCAT CGC CCG GAT CAG CTG GGC GCG GCG GCG ACC GCG TAT AAA GCG TAT ACC ACC AAC ACC ACC CGC GCT GGC AAA CGC CAGGATGGCGAATGGG |
| 266 | 10 | CATGATGATTACCAGCTATCAT CGC CCG GAT CAG CTG GGC GCG GCG GCG ACC GCG TAT AAA GCG TAT ACC ACC AAC ACC ACC CGC GTG GCT AAA CGC CAGGATGGCGAATGGG |
| 267 | 10 | CATGATGATTACCAGCTATCAT CGC CCG GAT CAG CTG GGC GCG GCG GCG ACC GCG TAT AAA GCG TAT ACC ACC AAC ACC ACC CGC GTG GGC GCT CGC CAGGATGGCGAATGGG |
| 268 | 10 | CATGATGATTACCAGCTATCAT CGC CCG GAT CAG CTG GGC GCG GCG GCG ACC GCG TAT AAA GCG TAT ACC ACC AAC ACC ACC CGC GTG GGC AAA GCT CAGGATGGCGAATGGG |
| 269 | 11 | GGGCAAACGC GCT GAT GGC GAA TGG GTG AGC TAT AGC GTG TAT GGC GAA AAC GTG GAT TATGAACGCTATCC |
| 270 | 11 | GGGCAAACGC CAG GCT GGC GAA TGG GTG AGC TAT AGC GTG TAT GGC GAA AAC GTG GAT TATGAACGCTATCC |
| 271 | 11 | GGGCAAACGC CAG GAT GCT GAA TGG GTG AGC TAT AGC GTG TAT GGC GAA AAC GTG GAT TATGAACGCTATCC |
| 272 | 11 | GGGCAAACGC CAG GAT GGC GCT TGG GTG AGC TAT AGC GTG TAT GGC GAA AAC GTG GAT TATGAACGCTATCC |
| 273 | 11 | GGGCAAACGC CAG GAT GGC GAA GCT GTG AGC TAT AGC GTG TAT GGC GAA AAC GTG GAT TATGAACGCTATCC |
| 274 | 11 | GGGCAAACGC CAG GAT GGC GAA TGG GCT AGC TAT AGC GTG TAT GGC GAA AAC GTG GAT TATGAACGCTATCC |
| 275 | 11 | GGGCAAACGC CAG GAT GGC GAA TGG GTG GCT TAT AGC GTG TAT GGC GAA AAC GTG GAT TATGAACGCTATCC |
| 276 | 11 | GGGCAAACGC CAG GAT GGC GAA TGG GTG AGC GCT AGC GTG TAT GGC GAA AAC GTG GAT TATGAACGCTATCC |
| 277 | 11 | GGGCAAACGC CAG GAT GGC GAA TGG GTG AGC TAT GCT GTG TAT GGC GAA AAC GTG GAT TATGAACGCTATCC |
| 278 | 11 | GGGCAAACGC CAG GAT GGC GAA TGG GTG AGC TAT AGC GCT TAT GGC GAA AAC GTG GAT TATGAACGCTATCC |
| 279 | 11 | GGGCAAACGC CAG GAT GGC GAA TGG GTG AGC TAT AGC GTG GCT GGC GAA AAC GTG GAT TATGAACGCTATCC |
| 280 | 11 | GGGCAAACGC CAG GAT GGC GAA TGG GTG AGC TAT AGC GTG TAT GCT GAA AAC GTG GAT TATGAACGCTATCC |
| 281 | 11 | GGGCAAACGC CAG GAT GGC GAA TGG GTG AGC TAT AGC GTG TAT GGC GCT AAC GTG GAT TATGAACGCTATCC |
| 282 | 11 | GGGCAAACGC CAG GAT GGC GAA TGG GTG AGC TAT AGC GTG TAT GGC GAA GCT GTG GAT TATGAACGCTATCC |
| 283 | 11 | GGGCAAACGC CAG GAT GGC GAA TGG GTG AGC TAT AGC GTG TAT GGC GAA AAC GCT GAT TATGAACGCTATCC |
| 284 | 11 | GGGCAAACGC CAG GAT GGC GAA TGG GTG AGC TAT AGC GTG TAT GGC GAA AAC GTG GCT TATGAACGCTATCC |
| 285 | 12 | GCGAAAACGTGGAT GCT GAA CGC TAT CCG GTG GCG CAT CTG CAG GAA GAA GCG GAT GCG TGC TATGAAAGCCTGG |
| 286 | 12 | GCGAAAACGTGGAT TAT GCT CGC TAT CCG GTG GCG CAT CTG CAG GAA GAA GCG GAT GCG TGC TATGAAAGCCTGG |
| 287 | 12 | GCGAAAACGTGGAT TAT GAA GCT TAT CCG GTG GCG CAT CTG CAG GAA GAA GCG GAT GCG TGC TATGAAAGCCTGG |
| 288 | 12 | GCGAAAACGTGGAT TAT GAA CGC GCT CCG GTG GCG CAT CTG CAG GAA GAA GCG GAT GCG TGC TATGAAAGCCTGG |
| 289 | 12 | GCGAAAACGTGGAT TAT GAA CGC TAT GCT GTG GCG CAT CTG CAG GAA GAA GCG GAT GCG TGC TATGAAAGCCTGG |
| 290 | 12 | GCGAAAACGTGGAT TAT GAA CGC TAT CCG GCT GCG CAT CTG CAG GAA GAA GCG GAT GCG TGC TATGAAAGCCTGG |
| 291 | 12 | GCGAAAACGTGGAT TAT GAA CGC TAT CCG GTG GGT CAT CTG CAG GAA GAA GCG GAT GCG TGC TATGAAAGCCTGG |
| 292 | 12 | GCGAAAACGTGGAT TAT GAA CGC TAT CCG GTG GCG GCT CTG CAG GAA GAA GCG GAT GCG TGC TATGAAAGCCTGG |
| 293 | 12 | GCGAAAACGTGGAT TAT GAA CGC TAT CCG GTG GCG CAT GCT CAG GAA GAA GCG GAT GCG TGC TATGAAAGCCTGG |
| 294 | 12 | GCGAAAACGTGGAT TAT GAA CGC TAT CCG GTG GCG CAT CTG GCT GAA GAA GCG GAT GCG TGC TATGAAAGCCTGG |
| 295 | 12 | GCGAAAACGTGGAT TAT GAA CGC TAT CCG GTG GCG CAT CTG CAG GCT GAA GCG GAT GCG TGC TATGAAAGCCTGG |
| 296 | 12 | GCGAAAACGTGGAT TAT GAA CGC TAT CCG GTG GCG CAT CTG CAG GAA GCT GCG GAT GCG TGC TATGAAAGCCTGG |
| 297 | 12 | GCGAAAACGTGGAT TAT GAA CGC TAT CCG GTG GCG CAT CTG CAG GAA GAA GGT GAT GCG TGC TATGAAAGCCTGG |
| 298 | 12 | GCGAAAACGTGGAT TAT GAA CGC TAT CCG GTG GCG CAT CTG CAG GAA GAA GCG GCT GCG TGC TATGAAAGCCTGG |
| 299 | 12 | GCGAAAACGTGGAT TAT GAA CGC TAT CCG GTG GCG CAT CTG CAG GAA GAA GCG GAT GGT TGC TATGAAAGCCTGG |
| 300 | 12 | GCGAAAACGTGGAT TAT GAA CGC TAT CCG GTG GCG CAT CTG CAG GAA GAA GCG GAT GCG GCT TATGAAAGCCTGG |
| 301 | 13 | CGGATGCGTGC GCT GAA AGC CTG GGC AAC ATG ATT ACC AGC CAG GTG CAG CCG TGC ACC CAGCGCGAAT |
| 302 | 13 | CGGATGCGTGC TAT GCT AGC CTG GGC AAC ATG ATT ACC AGC CAG GTG CAG CCG TGC ACC CAGCGCGAAT |
| 303 | 13 | CGGATGCGTGC TAT GAA GCT CTG GGC AAC ATG ATT ACC AGC CAG GTG CAG CCG TGC ACC CAGCGCGAAT |
| 304 | 13 | CGGATGCGTGC TAT GAA AGC GCT GGC AAC ATG ATT ACC AGC CAG GTG CAG CCG TGC ACC CAGCGCGAAT |
| 305 | 13 | CGGATGCGTGC TAT GAA AGC CTG GCT AAC ATG ATT ACC AGC CAG GTG CAG CCG TGC ACC CAGCGCGAAT |
| 306 | 13 | CGGATGCGTGC TAT GAA AGC CTG GGC GCT ATG ATT ACC AGC CAG GTG CAG CCG TGC ACC CAGCGCGAAT |
| 307 | 13 | CGGATGCGTGC TAT GAA AGC CTG GGC AAC GCT ATT ACC AGC CAG GTG CAG CCG TGC ACC CAGCGCGAAT |
| 308 | 13 | CGGATGCGTGC TAT GAA AGC CTG GGC AAC ATG GCT ACC AGC CAG GTG CAG CCG TGC ACC CAGCGCGAAT |
| 309 | 13 | CGGATGCGTGC TAT GAA AGC CTG GGC AAC ATG ATT GCT AGC CAG GTG CAG CCG TGC ACC CAGCGCGAAT |
| 310 | 13 | CGGATGCGTGC TAT GAA AGC CTG GGC AAC ATG ATT ACC GCT CAG GTG CAG CCG TGC ACC CAGCGCGAAT |
| 311 | 13 | CGGATGCGTGC TAT GAA AGC CTG GGC AAC ATG ATT ACC AGC GCT GTG CAG CCG TGC ACC CAGCGCGAAT |
| 312 | 13 | CGGATGCGTGC TAT GAA AGC CTG GGC AAC ATG ATT ACC AGC CAG GCT CAG CCG TGC ACC CAGCGCGAAT |
| 313 | 13 | CGGATGCGTGC TAT GAA AGC CTG GGC AAC ATG ATT ACC AGC CAG GTG GCT CCG TGC ACC CAGCGCGAAT |
| 314 | 13 | CGGATGCGTGC TAT GAA AGC CTG GGC AAC ATG ATT ACC AGC CAG GTG CAG GCT TGC ACC CAGCGCGAAT |
| 315 | 13 | CGGATGCGTGC TAT GAA AGC CTG GGC AAC ATG ATT ACC AGC CAG GTG CAG CCG GCT ACC CAGCGCGAAT |
| 316 | 13 | CGGATGCGTGC TAT GAA AGC CTG GGC AAC ATG ATT ACC AGC CAG GTG CAG CCG TGC GCT CAGCGCGAAT |
| 317 | 14 | TGCAGCCGTGCACC GCT CGC GAA TGC TAT GCG ATG GAT CAG AAA GTG TGC GCG GCG GTG GGC TTTAGCAGCGATGCGG |
| 318 | 14 | TGCAGCCGTGCACC CAG GCT GAA TGC TAT GCG ATG GAT CAG AAA GTG TGC GCG GCG GTG GGC TTTAGCAGCGATGCGG |
| 319 | 14 | TGCAGCCGTGCACC CAG CGC GCT TGC TAT GCG ATG GAT CAG AAA GTG TGC GCG GCG GTG GGC TTTAGCAGCGATGCGG |
| 320 | 14 | TGCAGCCGTGCACC CAG CGC GAA GCT TAT GCG ATG GAT CAG AAA GTG TGC GCG GCG GTG GGC TTTAGCAGCGATGCGG |
| 321 | 14 | TGCAGCCGTGCACC CAG CGC GAA TGC GCT GCG ATG GAT CAG AAA GTG TGC GCG GCG GTG GGC TTTAGCAGCGATGCGG |
| 322 | 14 | TGCAGCCGTGCACC CAG CGC GAA TGC TAT GGT ATG GAT CAG AAA GTG TGC GCG GCG GTG GGC TTTAGCAGCGATGCGG |
| 323 | 14 | TGCAGCCGTGCACC CAG CGC GAA TGC TAT GCG GCT GAT CAG AAA GTG TGC GCG GCG GTG GGC TTTAGCAGCGATGCGG |
| 324 | 14 | TGCAGCCGTGCACC CAG CGC GAA TGC TAT GCG ATG GCT CAG AAA GTG TGC GCG GCG GTG GGC TTTAGCAGCGATGCGG |
| 325 | 14 | TGCAGCCGTGCACC CAG CGC GAA TGC TAT GCG ATG GAT GCT AAA GTG TGC GCG GCG GTG GGC TTTAGCAGCGATGCGG |
| 326 | 14 | TGCAGCCGTGCACC CAG CGC GAA TGC TAT GCG ATG GAT CAG GCT GTG TGC GCG GCG GTG GGC TTTAGCAGCGATGCGG |
| 327 | 14 | TGCAGCCGTGCACC CAG CGC GAA TGC TAT GCG ATG GAT CAG AAA GCT TGC GCG GCG GTG GGC TTTAGCAGCGATGCGG |
| 328 | 14 | TGCAGCCGTGCACC CAG CGC GAA TGC TAT GCG ATG GAT CAG AAA GTG GCT GCG GCG GTG GGC TTTAGCAGCGATGCGG |
| 329 | 14 | TGCAGCCGTGCACC CAG CGC GAA TGC TAT GCG ATG GAT CAG AAA GTG TGC GGT GCG GTG GGC TTTAGCAGCGATGCGG |
| 330 | 14 | TGCAGCCGTGCACC CAG CGC GAA TGC TAT GCG ATG GAT CAG AAA GTG TGC GCG GGT GTG GGC TTTAGCAGCGATGCGG |
| 331 | 14 | TGCAGCCGTGCACC CAG CGC GAA TGC TAT GCG ATG GAT CAG AAA GTG TGC GCG GCG GCT GGC TTTAGCAGCGATGCGG |
| 332 | 14 | TGCAGCCGTGCACC CAG CGC GAA TGC TAT GCG ATG GAT CAG AAA GTG TGC GCG GCG GTG GCT TTTAGCAGCGATGCGG |
| 333 | 15 | GCGGCGGTGGGC GCT AGC AGC GAT GCG GGC GTG AAC AGC GCG ATG GTG GGC GAA GCG TAT TTTTATGCGTATGGCGGC |
| 334 | 15 | GCGGCGGTGGGC TTT GCT AGC GAT GCG GGC GTG AAC AGC GCG ATG GTG GGC GAA GCG TAT TTTTATGCGTATGGCGGC |
| 335 | 15 | GCGGCGGTGGGC TTT AGC GCT GAT GCG GGC GTG AAC AGC GCG ATG GTG GGC GAA GCG TAT TTTTATGCGTATGGCGGC |
| 336 | 15 | GCGGCGGTGGGC TTT AGC AGC GCT GCG GGC GTG AAC AGC GCG ATG GTG GGC GAA GCG TAT TTTTATGCGTATGGCGGC |
| 337 | 15 | GCGGCGGTGGGC TTT AGC AGC GAT GGT GGC GTG AAC AGC GCG ATG GTG GGC GAA GCG TAT TTTTATGCGTATGGCGGC |
| 338 | 15 | GCGGCGGTGGGC TTT AGC AGC GAT GCG GCT GTG AAC AGC GCG ATG GTG GGC GAA GCG TAT TTTTATGCGTATGGCGGC |
| 339 | 15 | GCGGCGGTGGGC TTT AGC AGC GAT GCG GGC GCT AAC AGC GCG ATG GTG GGC GAA GCG TAT TTTTATGCGTATGGCGGC |
| 340 | 15 | GCGGCGGTGGGC TTT AGC AGC GAT GCG GGC GTG GCT AGC GCG ATG GTG GGC GAA GCG TAT TTTTATGCGTATGGCGGC |
| 341 | 15 | GCGGCGGTGGGC TTT AGC AGC GAT GCG GGC GTG AAC GCT GCG ATG GTG GGC GAA GCG TAT TTTTATGCGTATGGCGGC |
| 342 | 15 | GCGGCGGTGGGC TTT AGC AGC GAT GCG GGC GTG AAC AGC GGT ATG GTG GGC GAA GCG TAT TTTTATGCGTATGGCGGC |
| 343 | 15 | GCGGCGGTGGGC TTT AGC AGC GAT GCG GGC GTG AAC AGC GCG GCT GTG GGC GAA GCG TAT TTTTATGCGTATGGCGGC |
| 344 | 15 | GCGGCGGTGGGC TTT AGC AGC GAT GCG GGC GTG AAC AGC GCG ATG GCT GGC GAA GCG TAT TTTTATGCGTATGGCGGC |
| 345 | 15 | GCGGCGGTGGGC TTT AGC AGC GAT GCG GGC GTG AAC AGC GCG ATG GTG GCT GAA GCG TAT TTTTATGCGTATGGCGGC |
| 346 | 15 | GCGGCGGTGGGC TTT AGC AGC GAT GCG GGC GTG AAC AGC GCG ATG GTG GGC GCT GCG TAT TTTTATGCGTATGGCGGC |
| 347 | 15 | GCGGCGGTGGGC TTT AGC AGC GAT GCG GGC GTG AAC AGC GCG ATG GTG GGC GAA GGT TAT TTTTATGCGTATGGCGGC |
| 348 | 15 | GCGGCGGTGGGC TTT AGC AGC GAT GCG GGC GTG AAC AGC GCG ATG GTG GGC GAA GCG GCT TTTTATGCGTATGGCGGC |
| 349 | 16 | GTGGGCGAAGCGTAT GCT TAT GCG TAT GGC GGC GTG GAT GGC GAA TGC GAT AGC GGC TAAGATATCTGTCTCTGCTTTTG |
| 350 | 16 | GTGGGCGAAGCGTAT TTT GCT GCG TAT GGC GGC GTG GAT GGC GAA TGC GAT AGC GGC TAAGATATCTGTCTCTGCTTTTG |
| 351 | 16 | GTGGGCGAAGCGTAT TTT TAT GGT TAT GGC GGC GTG GAT GGC GAA TGC GAT AGC GGC TAAGATATCTGTCTCTGCTTTTG |
| 352 | 16 | GTGGGCGAAGCGTAT TTT TAT GCG GCT GGC GGC GTG GAT GGC GAA TGC GAT AGC GGC TAAGATATCTGTCTCTGCTTTTG |
| 353 | 16 | GTGGGCGAAGCGTAT TTT TAT GCG TAT GCT GGC GTG GAT GGC GAA TGC GAT AGC GGC TAAGATATCTGTCTCTGCTTTTG |
| 354 | 16 | GTGGGCGAAGCGTAT TTT TAT GCG TAT GGC GCT GTG GAT GGC GAA TGC GAT AGC GGC TAAGATATCTGTCTCTGCTTTTG |
| 355 | 16 | GTGGGCGAAGCGTAT TTT TAT GCG TAT GGC GGC GCT GAT GGC GAA TGC GAT AGC GGC TAAGATATCTGTCTCTGCTTTTG |
| 356 | 16 | GTGGGCGAAGCGTAT TTT TAT GCG TAT GGC GGC GTG GCT GGC GAA TGC GAT AGC GGC TAAGATATCTGTCTCTGCTTTTG |
| 357 | 16 | GTGGGCGAAGCGTAT TTT TAT GCG TAT GGC GGC GTG GAT GCT GAA TGC GAT AGC GGC TAAGATATCTGTCTCTGCTTTTG |
| 358 | 16 | GTGGGCGAAGCGTAT TTT TAT GCG TAT GGC GGC GTG GAT GGC GCT TGC GAT AGC GGC TAAGATATCTGTCTCTGCTTTTG |
| 359 | 16 | GTGGGCGAAGCGTAT TTT TAT GCG TAT GGC GGC GTG GAT GGC GAA GCT GAT AGC GGC TAAGATATCTGTCTCTGCTTTTG |
| 360 | 16 | GTGGGCGAAGCGTAT TTT TAT GCG TAT GGC GGC GTG GAT GGC GAA TGC GCT AGC GGC TAAGATATCTGTCTCTGCTTTTG |
| 361 | 16 | GTGGGCGAAGCGTAT TTT TAT GCG TAT GGC GGC GTG GAT GGC GAA TGC GAT GCT GGC TAAGATATCTGTCTCTGCTTTTG |
| 362 | 16 | GTGGGCGAAGCGTAT TTT TAT GCG TAT GGC GGC GTG GAT GGC GAA TGC GAT AGC GCT TAAGATATCTGTCTCTGCTTTTG |
| **Saturation mutagenesis variants** | | |
| L293R | 293 | CATCGGTCTCA GCGAAAACGTGGAT TAT GAA CGC TAT CCG GTG GCG CAT **AGA** CAG GAA GAA GCG GAT GCG TGC TATGAAAGCCTGG TGAGACCTGAG |
| L293H | 293 | CATCGGTCTCA GCGAAAACGTGGAT TAT GAA CGC TAT CCG GTG GCG CAT **CAT** CAG GAA GAA GCG GAT GCG TGC TATGAAAGCCTGG TGAGACCTGAG |
| L293K | 293 | CATCGGTCTCA GCGAAAACGTGGAT TAT GAA CGC TAT CCG GTG GCG CAT **AAA** CAG GAA GAA GCG GAT GCG TGC TATGAAAGCCTGG TGAGACCTGAG |
| L293D | 293 | CATCGGTCTCA GCGAAAACGTGGAT TAT GAA CGC TAT CCG GTG GCG CAT **GAT** CAG GAA GAA GCG GAT GCG TGC TATGAAAGCCTGG TGAGACCTGAG |
| L293E | 293 | CATCGGTCTCA GCGAAAACGTGGAT TAT GAA CGC TAT CCG GTG GCG CAT **GAA** CAG GAA GAA GCG GAT GCG TGC TATGAAAGCCTGG TGAGACCTGAG |
| L293S | 293 | CATCGGTCTCA GCGAAAACGTGGAT TAT GAA CGC TAT CCG GTG GCG CAT **TCT** CAG GAA GAA GCG GAT GCG TGC TATGAAAGCCTGG TGAGACCTGAG |
| L293T | 293 | CATCGGTCTCA GCGAAAACGTGGAT TAT GAA CGC TAT CCG GTG GCG CAT **ACT** CAG GAA GAA GCG GAT GCG TGC TATGAAAGCCTGG TGAGACCTGAG |
| L293N | 293 | CATCGGTCTCA GCGAAAACGTGGAT TAT GAA CGC TAT CCG GTG GCG CAT **AAT** CAG GAA GAA GCG GAT GCG TGC TATGAAAGCCTGG TGAGACCTGAG |
| L293Q | 293 | CATCGGTCTCA GCGAAAACGTGGAT TAT GAA CGC TAT CCG GTG GCG CAT **CAA** CAG GAA GAA GCG GAT GCG TGC TATGAAAGCCTGG TGAGACCTGAG |
| L293C | 293 | CATCGGTCTCA GCGAAAACGTGGAT TAT GAA CGC TAT CCG GTG GCG CAT **TGT** CAG GAA GAA GCG GAT GCG TGC TATGAAAGCCTGG TGAGACCTGAG |
| L293G | 293 | CATCGGTCTCA GCGAAAACGTGGAT TAT GAA CGC TAT CCG GTG GCG CAT **GGT** CAG GAA GAA GCG GAT GCG TGC TATGAAAGCCTGG TGAGACCTGAG |
| L293P | 293 | CATCGGTCTCA GCGAAAACGTGGAT TAT GAA CGC TAT CCG GTG GCG CAT **CCA** CAG GAA GAA GCG GAT GCG TGC TATGAAAGCCTGG TGAGACCTGAG |
| L293A | 293 | CATCGGTCTCA GCGAAAACGTGGAT TAT GAA CGC TAT CCG GTG GCG CAT **GCT** CAG GAA GAA GCG GAT GCG TGC TATGAAAGCCTGG TGAGACCTGAG |
| L293V | 293 | CATCGGTCTCA GCGAAAACGTGGAT TAT GAA CGC TAT CCG GTG GCG CAT **GTT** CAG GAA GAA GCG GAT GCG TGC TATGAAAGCCTGG TGAGACCTGAG |
| L293I | 293 | CATCGGTCTCA GCGAAAACGTGGAT TAT GAA CGC TAT CCG GTG GCG CAT **ATT** CAG GAA GAA GCG GAT GCG TGC TATGAAAGCCTGG TGAGACCTGAG |
| L293L | 293 | CATCGGTCTCA GCGAAAACGTGGAT TAT GAA CGC TAT CCG GTG GCG CAT **TTG** CAG GAA GAA GCG GAT GCG TGC TATGAAAGCCTGG TGAGACCTGAG |
| L293M | 293 | CATCGGTCTCA GCGAAAACGTGGAT TAT GAA CGC TAT CCG GTG GCG CAT **ATG** CAG GAA GAA GCG GAT GCG TGC TATGAAAGCCTGG TGAGACCTGAG |
| L293F | 293 | CATCGGTCTCA GCGAAAACGTGGAT TAT GAA CGC TAT CCG GTG GCG CAT **TTT** CAG GAA GAA GCG GAT GCG TGC TATGAAAGCCTGG TGAGACCTGAG |
| L293Y | 293 | CATCGGTCTCA GCGAAAACGTGGAT TAT GAA CGC TAT CCG GTG GCG CAT **TAT** CAG GAA GAA GCG GAT GCG TGC TATGAAAGCCTGG TGAGACCTGAG |
| L293W | 293 | CATCGGTCTCA GCGAAAACGTGGAT TAT GAA CGC TAT CCG GTG GCG CAT **TGG** CAG GAA GAA GCG GAT GCG TGC TATGAAAGCCTGG TGAGACCTGAG |
| G357R | 357 | CATCGGTCTCA GTGGGCGAAGCGTAT TTT TAT GCG TAT GGC GGC GTG GAT **AGA** GAA TGC GAT AGC GGC TAAGATATCTGTCTCTGCTTTTG TGAGACCTGAG |
| G357H | 357 | CATCGGTCTCA GTGGGCGAAGCGTAT TTT TAT GCG TAT GGC GGC GTG GAT **CAT** GAA TGC GAT AGC GGC TAAGATATCTGTCTCTGCTTTTG TGAGACCTGAG |
| G357K | 357 | CATCGGTCTCA GTGGGCGAAGCGTAT TTT TAT GCG TAT GGC GGC GTG GAT **AAA** GAA TGC GAT AGC GGC TAAGATATCTGTCTCTGCTTTTG TGAGACCTGAG |
| G357D | 357 | CATCGGTCTCA GTGGGCGAAGCGTAT TTT TAT GCG TAT GGC GGC GTG GAT **GAT** GAA TGC GAT AGC GGC TAAGATATCTGTCTCTGCTTTTG TGAGACCTGAG |
| G357E | 357 | CATCGGTCTCA GTGGGCGAAGCGTAT TTT TAT GCG TAT GGC GGC GTG GAT **GAA** GAA TGC GAT AGC GGC TAAGATATCTGTCTCTGCTTTTG TGAGACCTGAG |
| G357S | 357 | CATCGGTCTCA GTGGGCGAAGCGTAT TTT TAT GCG TAT GGC GGC GTG GAT **TCT** GAA TGC GAT AGC GGC TAAGATATCTGTCTCTGCTTTTG TGAGACCTGAG |
| G357T | 357 | CATCGGTCTCA GTGGGCGAAGCGTAT TTT TAT GCG TAT GGC GGC GTG GAT **ACT** GAA TGC GAT AGC GGC TAAGATATCTGTCTCTGCTTTTG TGAGACCTGAG |
| G357N | 357 | CATCGGTCTCA GTGGGCGAAGCGTAT TTT TAT GCG TAT GGC GGC GTG GAT **AAT** GAA TGC GAT AGC GGC TAAGATATCTGTCTCTGCTTTTG TGAGACCTGAG |
| G357Q | 357 | CATCGGTCTCA GTGGGCGAAGCGTAT TTT TAT GCG TAT GGC GGC GTG GAT **CAA** GAA TGC GAT AGC GGC TAAGATATCTGTCTCTGCTTTTG TGAGACCTGAG |
| G357C | 357 | CATCGGTCTCA GTGGGCGAAGCGTAT TTT TAT GCG TAT GGC GGC GTG GAT **TGT** GAA TGC GAT AGC GGC TAAGATATCTGTCTCTGCTTTTG TGAGACCTGAG |
| G357G | 357 | CATCGGTCTCA GTGGGCGAAGCGTAT TTT TAT GCG TAT GGC GGC GTG GAT **GGT** GAA TGC GAT AGC GGC TAAGATATCTGTCTCTGCTTTTG TGAGACCTGAG |
| G357P | 357 | CATCGGTCTCA GTGGGCGAAGCGTAT TTT TAT GCG TAT GGC GGC GTG GAT **CCA** GAA TGC GAT AGC GGC TAAGATATCTGTCTCTGCTTTTG TGAGACCTGAG |
| G357A | 357 | CATCGGTCTCA GTGGGCGAAGCGTAT TTT TAT GCG TAT GGC GGC GTG GAT **GCT** GAA TGC GAT AGC GGC TAAGATATCTGTCTCTGCTTTTG TGAGACCTGAG |
| G357V | 357 | CATCGGTCTCA GTGGGCGAAGCGTAT TTT TAT GCG TAT GGC GGC GTG GAT **GTT** GAA TGC GAT AGC GGC TAAGATATCTGTCTCTGCTTTTG TGAGACCTGAG |
| G357I | 357 | CATCGGTCTCA GTGGGCGAAGCGTAT TTT TAT GCG TAT GGC GGC GTG GAT **ATT** GAA TGC GAT AGC GGC TAAGATATCTGTCTCTGCTTTTG TGAGACCTGAG |
| G357L | 357 | CATCGGTCTCA GTGGGCGAAGCGTAT TTT TAT GCG TAT GGC GGC GTG GAT **TTG** GAA TGC GAT AGC GGC TAAGATATCTGTCTCTGCTTTTG TGAGACCTGAG |
| G357M | 357 | CATCGGTCTCA GTGGGCGAAGCGTAT TTT TAT GCG TAT GGC GGC GTG GAT **ATG** GAA TGC GAT AGC GGC TAAGATATCTGTCTCTGCTTTTG TGAGACCTGAG |
| G357F | 357 | CATCGGTCTCA GTGGGCGAAGCGTAT TTT TAT GCG TAT GGC GGC GTG GAT **TTT** GAA TGC GAT AGC GGC TAAGATATCTGTCTCTGCTTTTG TGAGACCTGAG |
| G357Y | 357 | CATCGGTCTCA GTGGGCGAAGCGTAT TTT TAT GCG TAT GGC GGC GTG GAT **TAT** GAA TGC GAT AGC GGC TAAGATATCTGTCTCTGCTTTTG TGAGACCTGAG |
| G357W | 357 | CATCGGTCTCA GTGGGCGAAGCGTAT TTT TAT GCG TAT GGC GGC GTG GAT **TGG** GAA TGC GAT AGC GGC TAAGATATCTGTCTCTGCTTTTG TGAGACCTGAG |
